# Uncovering the features of Measles-targeting human antibodies elicited by the MMR vaccine

**DOI:** 10.1101/2025.09.09.675230

**Authors:** Marissa Acciani, Dawid Zyla, Gele Niemeyer, Stephanie Harkins, Diptiben Parekh, Emily Pawlack, Davide LaCarbonara, Stefan Niewiesk, Matteo Porotto, Kathryn M. Hastie, Erica Ollmann Saphire

## Abstract

Measles virus (MeV), a highly transmissible paramyxovirus, causes disease that can lead to severe complications and death, particularly in babies and young children. Deployment of the durable, highly effective, live-attenuated measles vaccine has saved an estimated 94 million lives in the past 50 years,^1^ yet the immunological explanation for this vaccine’s unique success and its landscape of antibody recognition remains unclear. Here we report the first panel of human monoclonal antibodies (mAbs) specific for the MeV hemagglutinin (H) and fusion (F) surface proteins, derived from the memory B cells of an MMR vaccinee. From over 100 cloned human mAbs, we mapped four major epitope clusters on H and another five major clusters on F, and structurally characterized 17 representative mAbs including one or more examples of each of the nine epitope clusters on the two surface antigens. We find that antibodies against both H and F can lead to potent virus neutralization and reduction of viral loads *in vivo*, including one mAb against F that reduces viral loads to below the limit of detection for all animals. High-resolution cryo-EM reveals contact sites of the most protective antibodies against both surface antigens. Discovery, characterization, and *in vitro* and *in vivo* success of these fully human mAbs now provide new avenues for prophylactic or therapeutic intervention against this re-emerging virus.

**Highlights:**

- A large panel of Measles-specific monoclonal antibodies was isolated from a human MMR vaccinee, years after vaccination.
- Structural and biochemical mapping paints a landscape of antibody recognition with nine major competition groups, including four major sites on Hemagglutinin (H) and five on the fusion protein (F).
- Antibodies against both H and F confer *in vitro* neutralization and *in vivo* protection, including, in one case, undetectable viral load after antibody treatment.
- The most protective H-specific mAbs, 4D08 and 1C08, target the receptor-binding site and the F-proximal outside of the H dimer, and likely function by interfering with receptor binding and H-F interactions, respectively.
- The most protective F-specific mAbs, 3A12 and 4F09, target the sides and apex of the prefusion F trimer, and likely function by locking F into its pre-fusion state

## Introduction

Measles virus (MeV), an enveloped negative-sense RNA paramyxovirus, has been endemic in the human population since around 600 BCE, coinciding with the shift to more densely populated communities with domesticated animals.^2^ MeV particles were first isolated in 1954, and MeV vaccines were developed shortly thereafter by serially passaging this clinical isolate for attenuation.^3,4^ Deployment of the MeV vaccine has now saved an estimated 94 million lives.^1^ This vaccine, derived from a genotype A MeV strain isolated in the 1960s, provides vaccinees with protective antibodies against all 24 MeV genotypes in circulation and lifelong immunity.^5^ Although the MeV vaccine has shown remarkable success in protecting the vaccinated global population from what was previously a universal childhood infection, there are only a few studies that have attempted to map the human antibody response from either measles vaccination or natural infection, and structural information on human anti-MeV antibodies is still lacking.

Measles virus spreads by respiratory droplets and is one of the most highly transmissible viruses known, with an estimated basic reproductive number of R_0_ of 12-18, although an R_0_ as high as 200 was recorded in one outbreak.^6^ Vaccine coverage of ∼95% is required to achieve herd immunity.^7^ In recent years, a drop in vaccination rates to ∼83% first dose coverage and 74% second dose coverage has resulted in surges of measles virus cases worldwide, with over 300,000 confirmed and suspected cases this year (as of July 2025), including at least 35 outbreaks in the United States alone.^8–10^ Using statistical modeling to capture further unobserved cases, Minta *et al*. estimated there were approximately 10 million cases for 2023 worldwide.^11,12^ The most vulnerable people in these outbreaks are the immunocompromised, including pediatric cancer patients and pregnant women, who are unable to receive a live-attenuated viral vaccine, as well as young children not yet vaccinated. The first dose of vaccine is typically given at 12-15 months of age and the second at 4-6 years of age, leaving these children susceptible to MeV disease and reliant on herd immunity.

MeV infection can cause a wide range of symptoms, from acute manifestations like high fever, runny nose, conjunctivitis, and a systemic skin rash, to severe and possibly fatal respiratory and neurological complications. In some cases, MeV can directly or indirectly induce encephalitis, resulting in permanent neurological sequelae or death. Further, long-term MeV replication in the brain (6-15 years after acute infection) may lead to subacute sclerosing panencephalitis (SSPE), which is almost always fatal.^13–16^ MeV is also a major cause of vision impairment and blindness in children in resource-limited areas.^17,18^ In nearly all patients, whether pediatric or adult, mild or severe, MeV infection increases risk for other infectious diseases like pneumonia as a result of “immune amnesia,” an immunosuppressive state lasting weeks to years after acute infection.^19–22^

There are currently no specific treatments available for measles virus infection, despite the numbers of people who aren’t vaccinated, can’t be vaccinated, or who experience breakthrough infections. The only available post-exposure intervention is inoculation of sera obtained from a pool of vaccinated individuals.^23^ Human monoclonal antibodies (mAbs) could offer valuable pre- or post-exposure protection for our most vulnerable populations, but none have been described to date.

Protective antibodies against measles virus may target one of two essential surface glycoproteins, Hemagglutinin (H) and Fusion (F), which together mediate virus-cell attachment, internalization, and fusion. MeV H is a 75 kDa single-pass transmembrane protein that forms disulfide-linked dimers or loosely assembled tetramers on the viral surface.^24–27^ The globular C-terminal head domains on wild-type H bind to the cellular receptors SLAMF1 (CD150) on alveolar macrophages, dendritic cells, and macrophages, and nectin4 on respiratory epithelial cells.^25,28–32^ Laboratory-adapted MeV H may also use CD46 as an additional receptor.^33,34^ Thus far, no *bona fide* MeV receptor has been identified in human brain cells. MeV F is a 58 kDa class I viral glycoprotein that exists as a metastable trimer on the virus surface, and catalyzes fusion of the virus and host membranes by refolding to a stable six-helix bundle. F is translated as a single polypeptide chain that gets cleaved into F1 and F2 disulfide-linked subunits by furin-like protease during transport through the Golgi. The F1 subunit contains the fusion peptide (FP) and N- and C-terminal heptad repeats (HRN and HRC), while the F2 subunit contains the stable signal peptide. Following receptor binding by H and triggering of F, the HRN refolds to transition F into an extended prehairpin intermediate state, allowing the FP to insert into the target membrane. The trimer then collapses into an HRN-HRC-mediated six-helix bundle, bringing the membranes together to form the fusion pore.^35^ H-F signaling that initiates F refolding and fusion relies on H-receptor interactions and signaling through the H stalk, via a mechanism that is still under investigation.^25,26,36–38^

Previous studies suggest that while human vaccinees generate antibodies against both MeV H and F, protection against measles infection correlates best with the level of neutralizing antibodies targeting H.^39–46^ The neutralizing activity of anti-F antibodies has been demonstrated only in individuals infected with wild-type virus.^43^ The structural basis underlying these protective antibodies remains unresolved.

Structures are available for the soluble monomeric and dimeric H head domain alone and in complex with receptors SLAMF1 and nectin-4, but there are not yet any structures of H bound to any antibody. In the absence of such structures, mapping of MeV immune recognition has been accomplished using murine mAbs and a combination of functional assays, mutagenesis, and peptide-based approaches. Combined, these studies describe residues in seven clusters on H, termed Ia, Ib, IIa, IIb, III, phi (ɸ), and the “noose” epitope.^44,47–58^ The human relevance of several of these sites has been evaluated using polyclonal human sera and reactivity to linear peptides or competition assays against representative murine mAbs.^44,47,59–64^ One study implicated sites IIa, III, and the noose as the most immunogenic in humans.^65^ Another study used detergent-solubilized antigens from the moderately-attenuated LEC strain to pan for human antibody fragments (Fabs), yielding two human Fabs targeting H.^66^ These two Fabs compete with murine group III mAbs shown to block H-receptor binding.^67^ None of the aforementioned studies, however, performed direct structural epitope mapping for H.

Fewer studies have defined immunogenic epitopes on F. Peptide scans and mutagenesis studies in the 1980s and 1990s indicated several immunodominant regions throughout the C-terminal F1 subunit and N-terminal F2 subunit, again relying on mouse mAbs and human polyclonal sera.^57,60,61,68–72^ The prefusion F ectodomain structure was solved alone and in complex with small molecular inhibitors^35,73^ and one murine Fab,^35,73^ but there are no structures of measles virus F in complex with any human antibody.

Here, we sought to map the human antibody response elicited in response to the measles virus vaccine. We developed mAbs from the peripheral memory B cells of a vaccinated 56-year-old female, resolved structures of mAb-H/F complexes, and mapped the epitopes associated with each competition group. We uncover potent and frequent neutralization by recognition of both H and F, including at least three neutralizing sites on H and four sites on F. *In vivo*, we find that antibodies targeting most of these sites (two on H and all four on F) drastically reduce infectious MeV titers in lungs when administered prophylactically. One anti-F mAb reduces viral titers to undetectable levels. Notably, the protective antibodies in this human panel appear durable, as they were isolated from circulating peripheral blood of an individual who had been first vaccinated over 50 years ago and boosted five years before blood donation (although the possibility of boost by an unknown wild-type viral exposure cannot be excluded). Therefore, the studies described here map a human antibody landscape against measles virus, and provide a framework to understand B cell recognition by a globally successful vaccine. Furthermore, the mAbs themselves open the possibility of therapeutic or prophylactic treatment in expanding outbreaks. Such treatments are needed for the growing populations of immunologically vulnerable people, including those who cannot be vaccinated and children too young to be vaccinated, where measles cases are surging.

## Results

### Isolation of antibodies from a thrice vaccinated person

We screened sera from a human donor pool in La Jolla, California for polyclonal antibody reactivity to wild-type full-length MeV H and full-length prefusion-stabilized F (with stabilizing mutations at E170G and E455G)^35,74^ glycoproteins to select a candidate for monoclonal antibody (mAb) discovery (Fig 1). Our donor pool contained 20 individuals who voluntarily completed a demographic and vaccination history survey. Fifteen donors indicated receiving the MMR vaccine (ages 19-61, median age 36, 10 female and 5 male), four donors did not report their MMR vaccination histories (ages 37-54, two ages unknown, three female and one sex unknown), and one donor indicated they did not receive the MMR vaccination (age 45, sex male). We sorted individuals by vaccination status and summed H/F sera reactivities and noted the most (#3920 and #0003) and least (#4168) reactive individuals (Fig 1A, outlined panel). As expected, no serum sample was completely negative for H/F-reactive polyclonal antibodies. It is highly likely that individuals in the “unknown” or ‘not vaccinated” categories were either vaccinated or exposed to wild-type MeV in their lifetimes and would therefore demonstrate some reactivity. We also performed correlative analyses of summed reactivity versus age and sex for vaccinated donors, as this was the only group with complete demographic information. We observed a significant effect of sex, with female donors having higher reactivity to measles antigens than male donors. For individuals with the highest (#3920 and #0003) and lowest (#4168) summed sera reactivities, we quantified the H- and F-specific memory B cells contained in peripheral blood mononuclear cells (PBMCs) using fluorescent wild-type H and pre-fusion stabilized F ectodomain probes (H_ECT_ residues 149-617 or F_ECT_ residues 1-495^35^)(Fig 1B). PBMCs from donor #3920 contained the most MeV+ memory B cells (302/43,512 or 0.69%). PBMCs from donors #0003 and #4168 contained similar quantities of MeV-specific memory B cells: 0.21% and 0.24%, respectively. Notably, we detected an approximately three-fold higher incidence of F-specific memory B cells relative to H-specific cells in all individuals.

**Fig 1.**
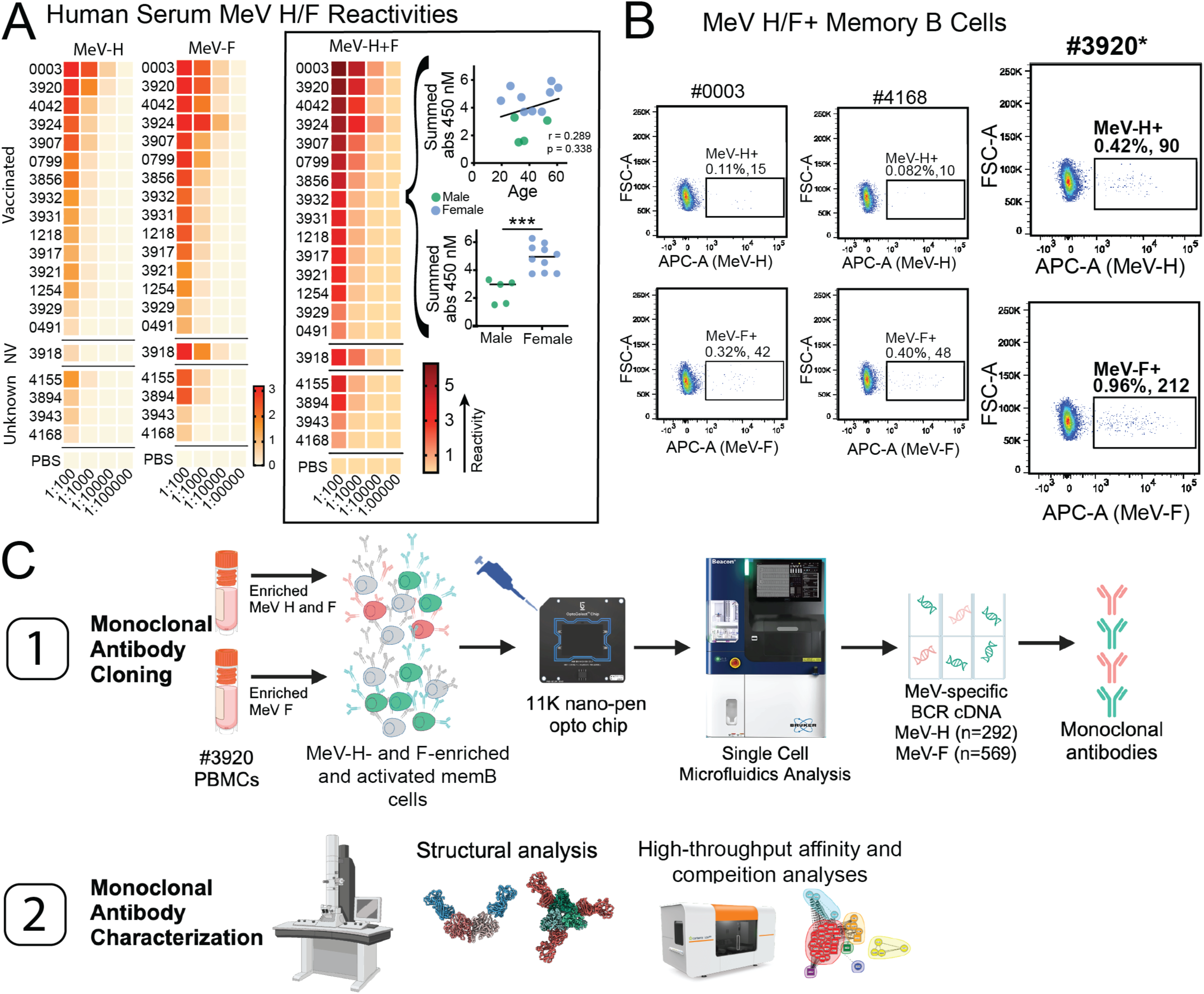
Vaccinee donor selection and MeV-specific memory B cell isolation. **A)** Sera from 20 individuals were screened for polyclonal antibody recognition of MeV H_FL_ or F_FL_ via ELISA (“NV” = not vaccinated). Reactivities for each individual and antigen were quantified by measuring optical density (OD) at 450nm, with data displayed in a heatmap (red = higher reactivity, beige = lower reactivity). Reactivities were summed across antigens (right outlined panel; H+F). Summed reactivities are also plotted by age and sex. In both plots, females are blue and males are green. **B)** PBMCs from three vaccinees were surface stained and single, live, MeV H+ or F+ memory B cells were quantified using flow cytometry. **C)** PBMCs from vaccinee #3920 were enriched for MeV H or F, activated into mAb-secreting cells, and loaded into a Beacon (Bruker) for high-throughput single cell analysis.

We selected donor #3920 for mAb discovery (Fig 1C). This 56-year-old donor reported that she had received two MeV vaccine doses in childhood, over 50 years before sample donation, and a third dose as an adult, five years before sample donation. PBMCs from donor #3920 were enriched for MeV antigen-specific memory B cells using biotinylated MeV H_ECT_ and F_ECT_ probes. Selecting with H_ECT_ initially introduced non-target cytotoxic lymphocytes that killed our enriched memory B cells; therefore, we eliminated cytotoxic cells by first pretreating PBMCs with L-Leucyl-L-Leucine methyl ester (LLME), and included a separate F_ECT_-alone enrichment as a precaution. MeV H- and F-specific memory B cells were activated for ten days, then loaded onto a Bruker Cell Analysis Beacon for analysis and selection of individual cells. We detected 292 and 569 single cells secreting MeV H- or F-specific mAbs, respectively, using fluorescent antigen probes (H_ECT_-AF555 and F_ECT_-AF647). The two-times greater number of F-specific memory B cells identified on the Beacon may be attributed to a higher incidence of F-specific memory B cells (Fig 1B), but could also be partially due to the addition of the aforementioned F_ECT_-only enriched/activated cells. We selected approximately 384 H- and F-specific cells for mAb discovery. Heavy and light variable regions were cloned and expressed as individual mAbs in expiCHO cells. Cloned mAbs were sequenced and produced in small quantities in cell supernatants, which were harvested and used for rough functional analyses (Fig S1A). We observed an inverse correlation between avidity and virus inhibition for F-specific mAbs, and no link between avidity/inhibition and heavy chain variable germline usage for H- and F-specific mAbs (Fig S1B).

### Defining human mAb epitopes on H and F

We combined high-throughput competition-based epitope binning and negative-stain electron microscopy (NS-EM) to roughly map mAb epitopes on MeV H and F (Figs 2-3). Fifty-two anti-H and 46 anti-F mAbs were purified and assessed for antigen-binding kinetics and epitope binning, then sorted into four and five epitope communities, respectively, where each community contains mAbs with similar competition behaviors (or a single mAb with unique competition behavior) (Fig 2A, 3A, Table S1). Representative mAbs from each community were selected for direct footprint mapping via NS-EM. High-quality micrographs of antigen-antibody complexes were used to construct 3D densities in cryoSPARC.^75^ Individual H_ECT_ and F_ECT_ models and Fabs were docked into densities in ChimeraX^76^ (Fig S2).

**Fig 2.**
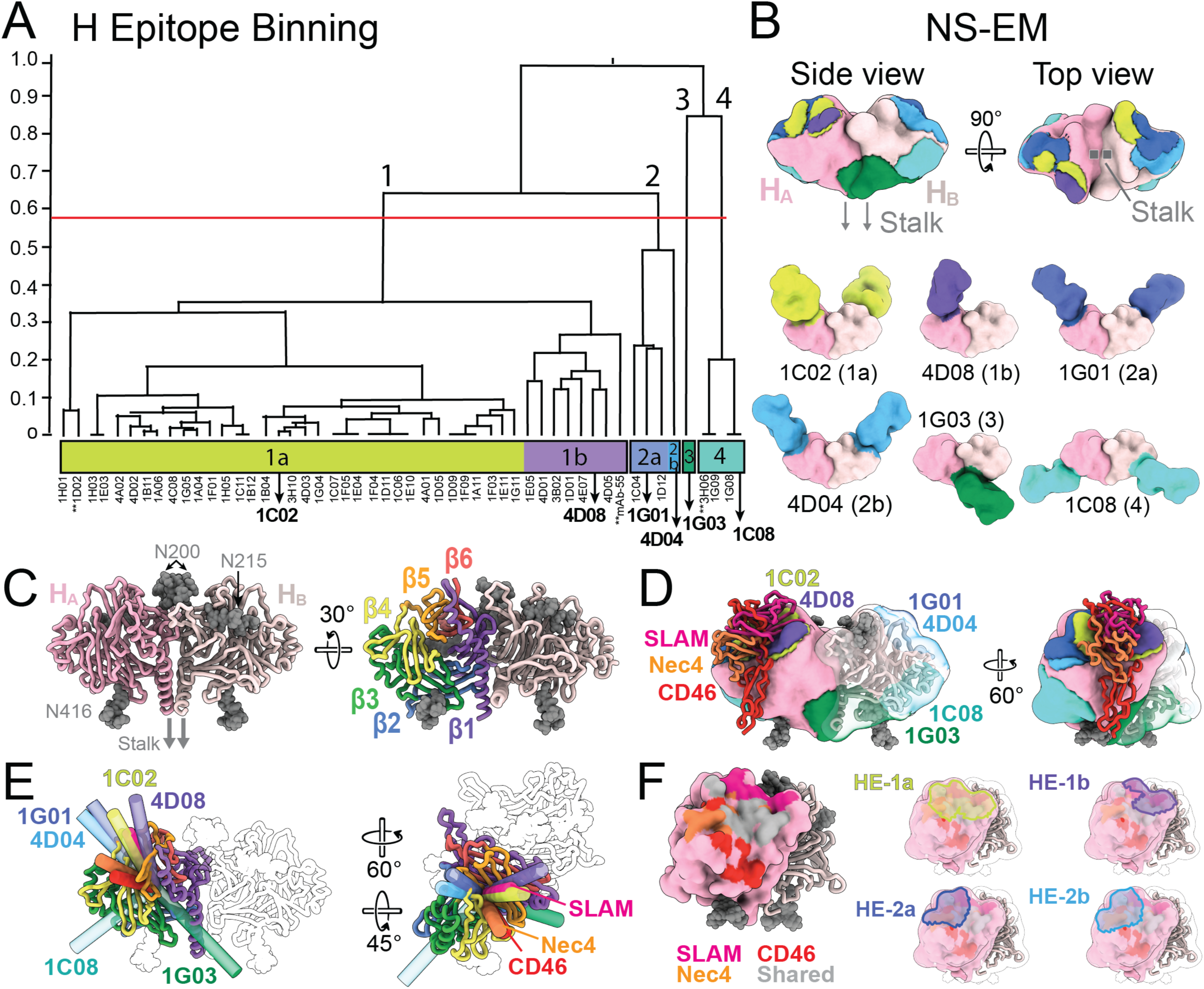
Monoclonal antibody epitope mapping on MeV H. **A)** High-throughput competition assays of purified mAbs with H_ECT_ were performed using a Carterra LSA. MAbs were assigned to bins based on blocking behaviors, as represented by a binning dendrogram, using Caterra Epitope software. Coarser epitope communities were generated by manually adjusting the dendrogram cut-heights (red lines). MAbs selected for NS-EM are bolded. The starred mAbs (**) were not directly visualized but were included in *in vitro* experiments. **B)** MAb antigen-binding fragment (Fab)-antigen complexes were subjected to negative-stain electron microscopy (NS-EM). Alphafold-derived structural models for H_ECT_ and Fab55 were docked into NS-EM densities, and Fab binding footprints were mapped onto H_ECT_ molecular surfaces. **C)** The H_ECT_ dimer modeled with homogeneous N-linked glycans^26,32^ (GlcNAc_2_Man_5_ gray spheres) and individual B propeller blades annotated. **D)** H-Fab footprint mapping combined with known H-SLAM, -nectin-4, and -CD46 structures (H_ECT_-SLAM PDB 3ALZ, H_ECT_-nectin-4 PDB 4GJT, H_ECT_-CD46 PDB 3INB). **E)** MAb and receptor approach angles relative to H_ECT_ were visually inspected by defining molecular axes. **F)** Surface-exposed residues on H critical for receptor binding are shown in the indicated colors, with individual epitopes HE-1a, HE-1b, HE-2a, and HE-2b overlaid on the H molecular surface for direct comparison. All structures were illustrated using Chimera X.^141^

**Fig 3.**
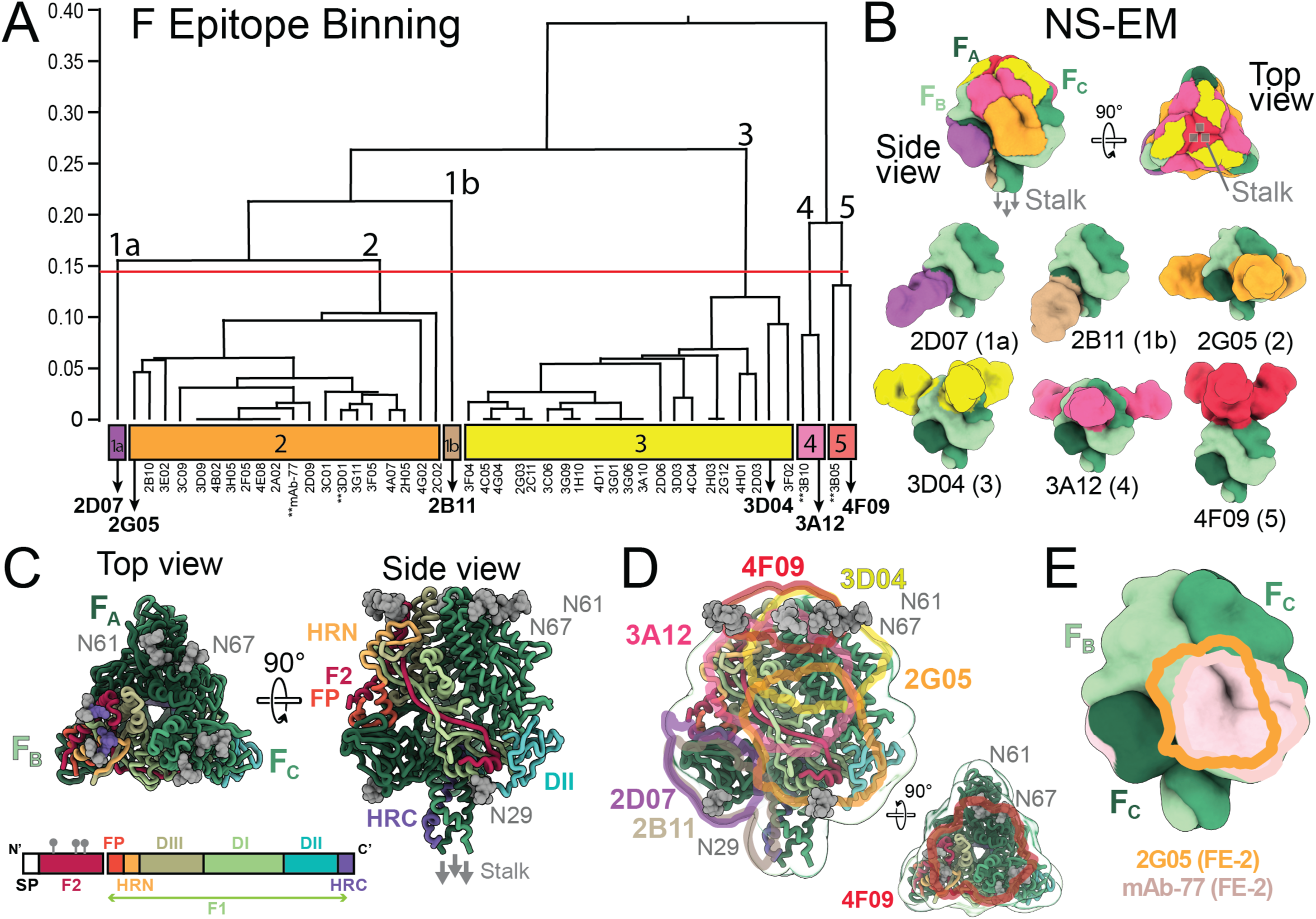
Monoclonal antibody epitope mapping on MeV F. **A)** High-throughput competition assays of purified mAbs with F_ECT_ were performed using a Carterra LSA. MAbs were assigned to epitope groups based on blocking behaviors, as represented by a binning dendrogram. Coarser epitope communities were generated by manually adjusting the dendrogram cut-heights (red lines). MAbs selected for NS-EM are bolded; starred mAbs (**) were not directly visualized but were included in *in vitro* experiments. **B)** Fab-antigen complexes were subjected to NS-EM. The cryo-EM-derived structural model for F_ECT_ (PDB 8UT2) and a Fab alphafold model were docked into NS-EM densities, and Fab binding footprints were mapped onto F_ECT_ molecular surfaces. **C)** The F_ECT_ trimer modeled with N-linked glycans (gray spheres), with functional and structural domains annotated for one protomer. **D)** F-Fab footprint mapping overlaid on the annotated F_ECT_ trimer, accompanied by a smaller apex view to better visualize epitope F6. **E)** Footprints for FE-2 mAbs 2G05 and mAb77 overlaid on the F_ECT_ trimer. All structures were illustrated using Chimera X.^141^

We formed Fab-H complexes using previously-characterized purified H_ECT_, which form disulfide-linked dimers of globular head domains.^26^ Each head domain contains a beta propeller of six blades (four antiparallel β-strands per blade) and contains three N-linked glycans at N200, N215, and N416.^26,32^ We identified four major epitope communities on H (HE-1 through HE-4), including several sub-epitopes designated HE-1a and HE-1b, and HE-2a and HE-2b, which target the apical surface of the dimer (Fig 2A-B). The majority (42/52, or 81%) of purified mAbs were sorted into HE-1, suggesting this could be a major immunogenic site. This epitope, consisting of sub-clusters HE-1a and HE-1b, overlaps with the binding sites for MeV receptors SLAMF1 and Nectin-4, and partially overlaps with that of MeV receptor CD46 (Fig 3C-F).^25,27,32^ HE-1a encompasses the inside and outer edges of a concave groove formed by H propeller blades β4, β5, and β6, which also contains the binding sites for SLAMF1^25^ and Nectin-4.^32^ HE-1b consists of a smaller region shifted towards the anterior edge of the groove, closer to blades β5 and β6 (Fig 3F). MAbs 1C02 and 4D08, used to define HE-1a and HE-1b, respectively, approach the groove at almost an identical angle relative to SLAMF1 (Fig 3E).

Epitope HE-2 maps to the posterior edge of this groove to a region containing blades β3, β4, and β5, and extends towards the posterior face of the dimer.

HE-3 and HE-4, defined by mAbs 1G03 and 1C08, respectively, approach the membrane-proximal undersides of the globular head domains. There, mAb 1G03 (HE-3), which binds one Fab per dimer, recognizes the dimer interface, engaging the β1 blade of both monomers and blades β1-β3 of a single monomer. This epitope appears to be most vulnerable to possible glycan shielding by the N-linked glycan at N416. MAb 1C08 (HE-4), which binds two Fabs per dimer, symmetrically recognizes the peripheral ends of each monomer at blades β2 and β3. Overall, we found that the majority of the exposed H dimer surface was targeted by mAbs, with a high incidence of mAbs targeting the receptor-binding regions.

We next complexed F-reactive mAbs to purified F_ECT_ trimers, which lack the transmembrane and cytoplasmic tail and contain naturally occurring mutations that slow the post-fusion transition.^35^ F is divided into F1 and F2 subunits, and contains N-linked glycans at positions N61 and N67 at the trimer apex and N29 at the membrane-proximal undersides of each monomer. We observe an even spread of six antibody recognition sites that ascend the prefusion F trimer vertically from the base to the apex, altogether nearly covering the entirety of the trimer’s surface (Fig 3B). Epitope FE-1, defined by mAbs 2D07 (FE-1a) and 2B11 (FE-1b), localizes to domain II of the F1 subunit at the corners of the trimer base. These mAbs were sorted into individual, separate epitope communities, likely owing to the fact that these mAbs did not fully saturate all epitopes on the F trimer, as observed in NS-EM 2D classes.

In contrast, most F-specific mAbs were grouped into FE-2 (19/46, 41%) and FE-3 (21/46, 46%), which were mapped to the lower and upper faces of the trimer, respectively. FE-2 mAbs target the N’ ɑ-helix and a large surface-exposed β strand in the F2 subunit (approximately residues 1-45) that stretches from the bottom right to upper left face of each monomer in the trimer, as well as a portion of DII. Murine mAb mAb-77^35,69,77^ also bins into this community, and footprint comparison analyses of human mAb 2G05 and murine mAb77^35^ indeed confirm highly overlapping epitopes (Fig 3E). FE-3 approaches the upper right edge of the trimer face, and likely encompasses the C’ F2 ɑ helix and the F1 HRN.

FE-4 lies more centrally on the trimer face, vertically positioned between epitopes F2 and F3. This epitope includes more membrane-distal regions of the N’ F2 β strand, and may include the C’ F2 ɑ helix. FE-5 is located at the apex of the trimer, and contains loops or hinge regions between secondary structures in the HRN, F2 and DIII domains. FE-5 seems most likely to be impacted by glycan shielding at N61 and N67, which may explain the presence of only two mAbs in this binning community (4F09 and 3B05).

Briefly, we evaluated the ability of AlphaFold3 (AF3) multimer modeling to predict mAb binding sites to within NS-EM accuracy (Fig S3). AF3 correctly predicted binding sites for 3 of the 12 complexes visualized experimentally: H_ECT_-4D04 (2b), H_ECT_-1C08 (H4), and F_ECT_-3D04 (F4), with 9-12 Å A-RMSD accuracy and 2-8 Å P-RMSD precision. Other anti-H and anti-F antibodies were predicted incorrectly with 58-77 Å A-RMSD, highlighting the importance of direct mAb-antigen complex visualization in epitope mapping.

### In vitro neutralization and in vivo mAb protection

We tested the potencies of H- and F-specific mAbs for *in vitro* neutralization using a recombinant MeV B3 strain bearing a fluorescent protein (mCherry) representing one of the two currently-circulating wild-type viruses^5^ (Fig 4). MeV-mCherry was pre-incubated with serially-diluted mAbs, then applied to Vero-SLAMF1 cells. Infected and uninfected cells were quantified 16 hours after infection.

**Fig 4.**
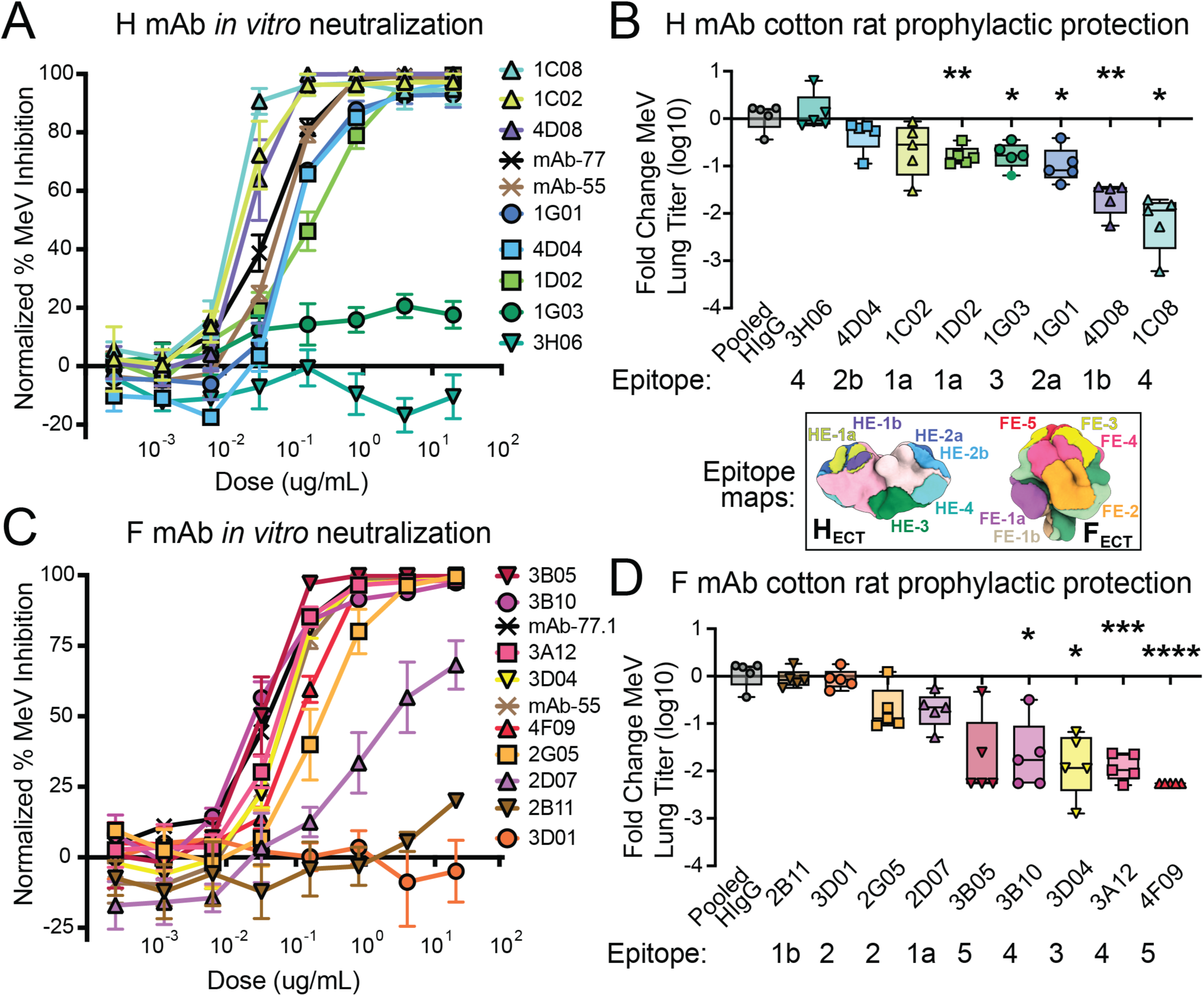
MeV mAb *in vitro* and *in vivo* efficacy. **A,C)** MAbs and MeV were combined and applied to cells to assess mAb neutralization efficiencies. Serially diluted mAbs (2.56×10^−4^ to 20 ug/mL) were mixed with 5×10^3^ PFU/mL MeV-mCherry for 1 h at room temperature (RT). MAb-virus mixtures were incubated with cells for 3 h, then cell infection media was replaced with media containing 500 nM FIP-HRC to inhibit cell-cell spread and syncytia formation. The next day, the proportions of mCherry-infected cells versus total cells were quantified using a Cellinsight CX5 plate reader (ThermoFisher). Shown are data normalized to % MeV inhibition using no mAb in infected wells, average ± SEM (n = 3-4 biological replicates). MAbs listed in the legends are arranged from lowest/best to highest/worst EC_50_ value. **B,D)** MeV titer reduction in the lungs (TCID_50_log10/g lung tissue), normalized and displayed as fold-change compared to intra-group PBS controls (Fig S5A). Cotton rats were injected intraperitoneally with each mAb (1mg/kg, 5 animals per mAb), PBS (3-4 animals), or pooled human IgG (20mg/kg, NT titer 640, 5 animals treated) 14 hours before infection. Animals were then inoculated intranasally with 2×10^5^ TCID_50_ MeV B3-GFP. Animal lungs were collected and homogenized 4 days after infection, and lung tissue homogenates were serially diluted and titrated on Vero-SLAMF1 cells. We subtracted average PBS-treated lung titers (TCID_50_log10/g lung tissue) from mAb-treated lung titers (TCID_50_log10/g lung tissue) and show the average reduction of lung titers (TCID_50_log10/g lung tissue) for each mAb treatment. * p<0.05, **p<0.005, ***p<0.0005, ***p<0.0001 by Brown-Forsythe and Welch ANOVA test (or Mann-Whitney test for mAb 4F09).

For mAbs against H, we found that those targeting HE-1a, HE-1b, HE-2a, and HE-2b at the receptor-binding site were highly neutralizing, with half-maximal effective concentration (EC_50_) values ranging from 0.019 to 0.176 µg/mL (Fig. 4A, Table 1). The sole member of epitope HE-3, mAb 1G03, which binds to the underside of H at the dimer interface, did not neutralize virus. MAbs from HE-4, which bind at the H periphery, varied: mAb 1C08 was the most potent virus inhibitor in the panel (EC_50_ 0.013 µg/mL), while 3H06 was non-neutralizing.

Overall, H-specific mAbs targeting the receptor-binding groove were neutralizing. MAb 1C08, which targets the side of H and may block F interactions, also neutralized. In contrast, those mAbs against the bottom of H did not neutralize.

For mAbs against F, we found that those targeting FE-3, FE-4, and FE-5 were the most neutralizing with EC_50_s ranging from 0.029 to 0.118 µg/mL (Fig 4C). FE-2 mAbs 2G05 and mAb 77 neutralized with EC_50_s 0.237 and 0.041 ug/mL respectively, while a third FE-2 mAb, 3D01, did not neutralize. FE-1 mAbs 2D07 and 2B11 demonstrated poor-to-moderate virus-inhibiting activity, with EC_50_ values of 3.35 and 64.84 µg/mL (Table S1).

Overall, F-specific mAbs binding the top half of the trimer were, in general, more neutralizing than those directed against the bottom half of the trimer. The most potent neutralizing F-specific mAbs, 3B10 and 3B05 (EC_50_s 0.029 and 0.031 µg/mL, respectively), target epitopes FE-4 and FE-5, both quaternary epitopes at the upper face and apex of F, respectively.

We next assessed the ability of these human mAbs to confer *in vivo* pre-exposure protection in the cotton rat model (*Sigmodon hispidus*), a semi-permissive animal model that replicates key aspects of human MeV infection, including viral replication in the lungs and MeV-induced immune suppression.^78^ Cotton rats were pre-treated with mAbs or PBS (1 mg/kg), then infected with 2×10^5^ TCID_50_ MeV-eGFP (genotype B3).^35^ Four days later, lung tissues were harvested and MeV titers were quantified. H- and F-specific mAbs were tested in four experimental groups, with four to six individual mAb treatments and one PBS control treatment per group, five animals per treatment (Fig S4A). One group also contained a pooled purified human IgG treatment (neutralizing titer 1:640). For each mAb treatment, we determined the fold-change (TCID_50_log10/g lung tissue) MeV lung tissue titer relative to group-specific PBS-only controls (Fig 4B-D).

In these experiments, two mAbs against H and five mAbs against F were highly protective, reducing MeV lung titers approximately 1.7-2.3 log₁₀/g lung tissue units relative to PBS controls, corresponding to a ∼50-200-fold reduction in viral load. Protective mAbs include 4D08 (HE-1b, receptor-binding site), 1C08 (HE-4, periphery), 3B05 (FE-5, apex), 3B10 (FE-4, upper center), 3D04 (FE-3, lower right), 3A12 (FE-4, upper center), and 4F09 (FE-5, apex). The most effective among them is the anti-F apex-binding mAb 4F09, which reduces lung titers to undetectable levels.

*In vitro* neutralization potency generally correlated with *in vivo* protection (r = 0.4964, p = 0.0623) for those mAbs that demonstrated quantifiable MeV neutralization. There were, however, mAbs which neutralized *in vitro* but failed to protect cotton rats at 1 mg/kg: 2G05 (F), 1C02 (H), 1G01 (H), and 4D04 (H). With 5 mg/kg dosing, two of these mAbs, 2G05 and 1C02 conferred 100% protection (titer < 10^3^ TCID50/g lung tissue) while the remaining two, 1G01 and 4D04, conferred only 20-60% protection (Fig. S4B).

### A detailed look into H and F complexed with protective mAbs using cryo-EM

We resolved structures of MeV H and F ectodomains complexed with the most protective mAbs for H (1C08 and 4D08) and F (4F09 and 3A12) using cryo-EM to more closely identify epitopes and illuminate possible structural bases for virus inhibition (Figs 5-6, Fig S5). We also describe the H-1C02 complex to examine structural discrepancies that may explain the reduced *in vivo* potency of 1C02 compared to mAb 4D08 of similar epitope. Detailed mAb-H/F interface information for every complex is listed in Table S2, and cryo-EM processing details are listed in Table S3.

**Figure 5.**
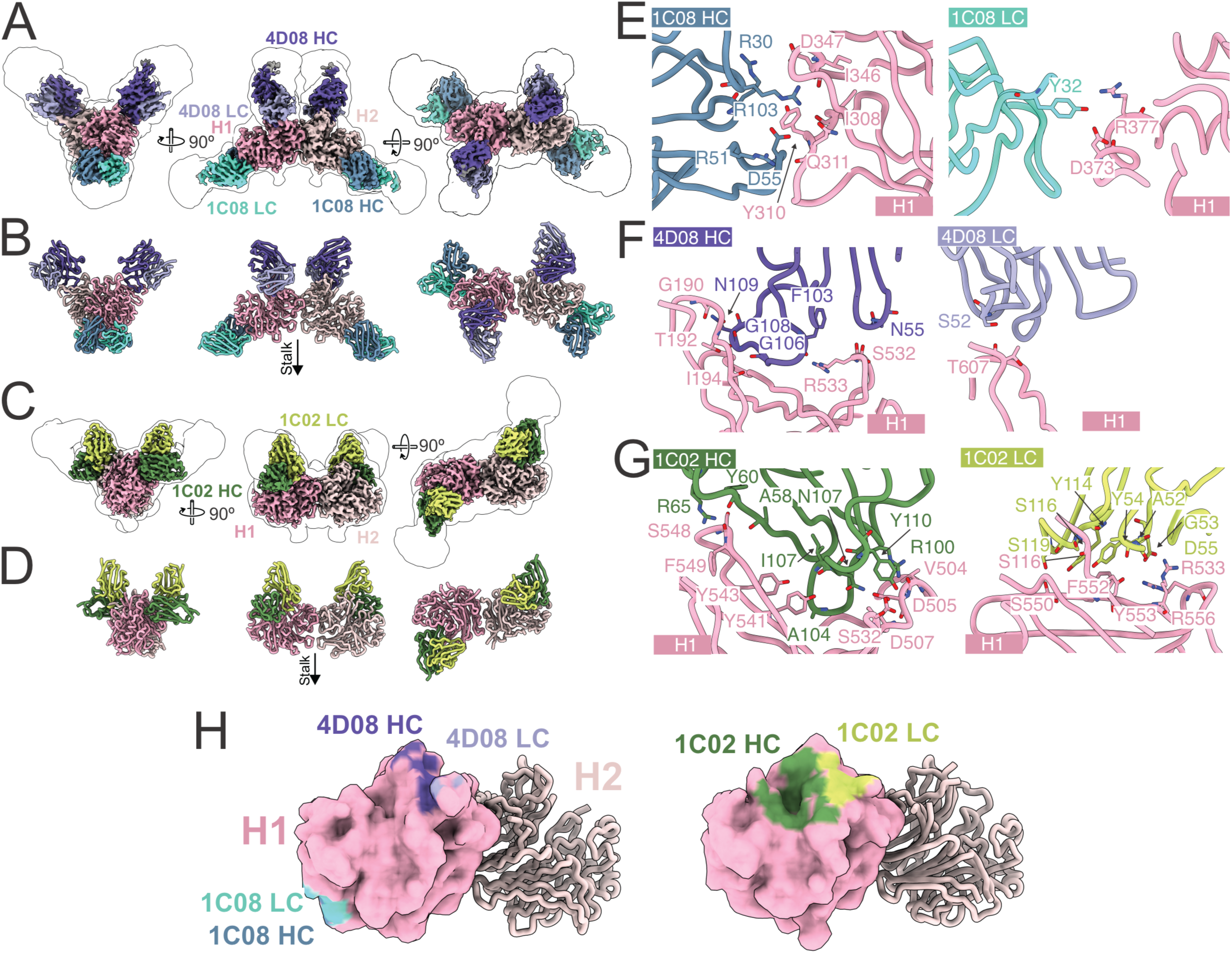
Cryo-EM structures of MeV H_ECT_ in complex with neutralizing antibodies. H_ECT_ monomers (pink shades) in complex with 4D08, 1C08, and 1C02 Fab fragments (light and heavy chains for each Fab are depicted in paired blue, purple, and green/yellow, respectively). The stalk region is indicated by an arrow, denoting the membrane-proximal orientation. The white silhouette outlines the low-pass filtered cryo-EM map (at 1.5 standard deviations), providing contextual density around the high-resolution maps. **(A)** Cryo-EM density map H_ECT_ in complex with Fabs 4D08 and 1C08. **(B)** Atomic model corresponding to (A), shown as ribbons. **(C)** Cryo-EM density map of H_ECT_ bound to antibody 1C02. **(D)** Corresponding ribbon model of the complex shown in (C). **(E)** Detailed view of the interaction interfaces between the heavy (left, dark blue) and light (right, cyan) chains of 1C08 and the H ectodomain (pink). **(F)** Interaction interfaces between the heavy (left, dark purple) and light (right, lavender) chains of 4D08 and H. **(G)** Interaction interfaces between the heavy (left, green) and light (right, yellow) chains of 1C02 and H. **(H)** Footprints of antibodies 1C08, 4D08, and 1C02 mapped onto the H dimer by coloring epitope residues within 3Å of each mAb. One H monomer is illustrated as a molecular surface and the second as a C alpha trace. Footprints by chain are colored as in E, F, and G.

**Figure 6.**
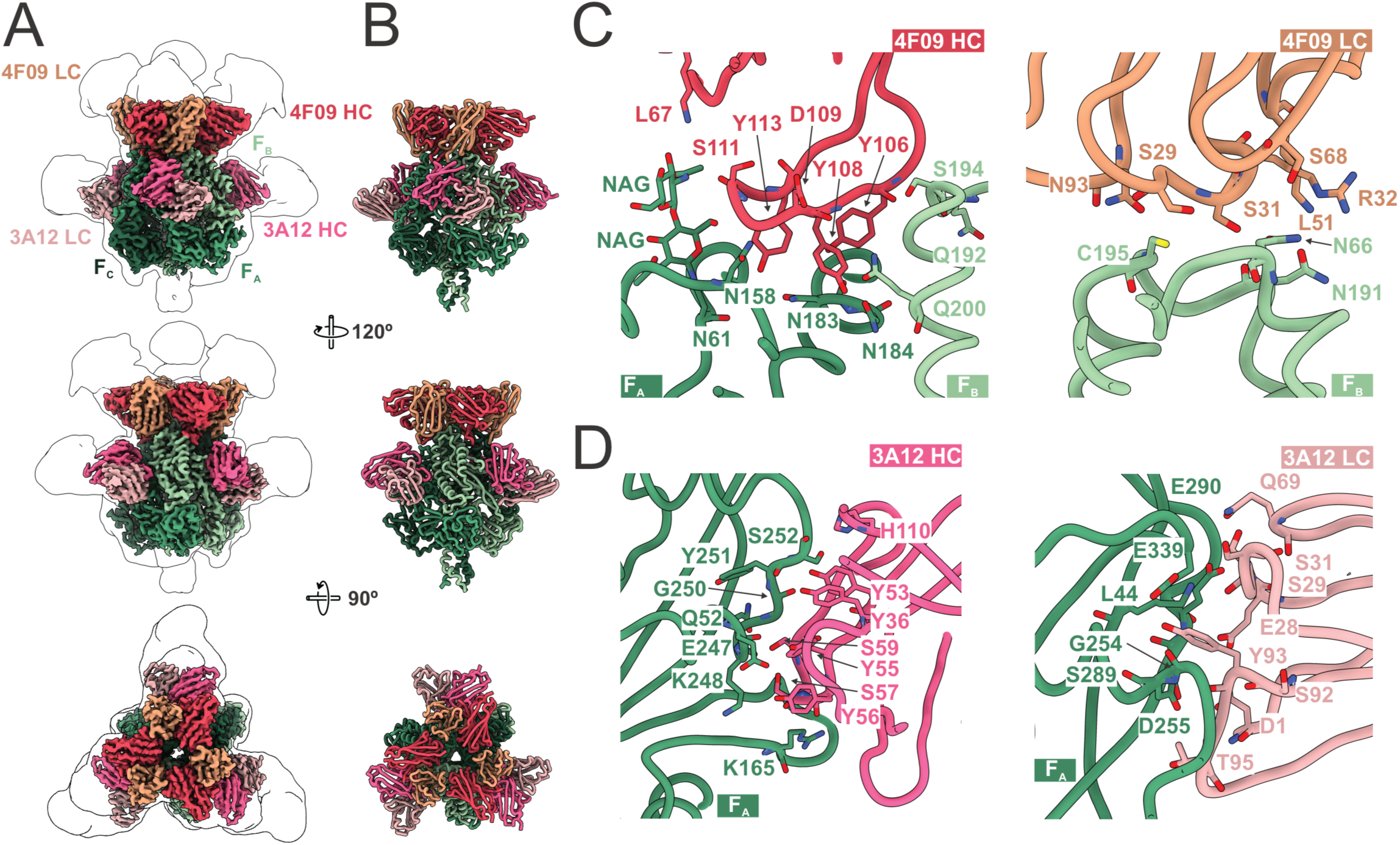
Cryo-EM structures of the measles virus F ectodomain in complex with neutralizing antibodies 4F09 and 3A12. F ectodomain protomers are shown in green, with Fab heavy and light chains distinguished by complementary colors. NAG denotes N-acetylglucosamine moieties. **(A)** Cryo-EM density map of the F_ECT_ trimer in complex with Fab fragments of 4F09 and 3A12, shown from three orthogonal views. The map silhouette represents a low-pass filtered density contoured at 1.5 standard deviations. **(B)** Corresponding ribbon models of the complex in the same orientations as in (A). The three F protomers are labeled as F_A_, F_B_, and F_C_. **(C)** Close-up views of the interface between 4F09 and the F ectodomain. Left: 4F09 heavy chain (red) engaging two protomers (F_A_ and F_B_), including interactions with glycans from N-glycosylated Asn61. Right: 4F09 light chain (orange) interactions with the F_B_ protomer. **(D)** Close-up views of the interface between 3A12 and the F ectodomain. Left: 3A12 heavy chain (magenta) binding a single protomer (F_A_), engaging both F1 and F2 fragments. Right: 3A12 light chain (light pink) interaction with F_A_, forming extensive contacts across the surface.

#### H Structures

Cryo-EM reconstructions of H ectodomain-mAb complexes revealed H_ECT_ homodimers throughout all stages of purification and structural analysis (Fig. 5A-D). As previously observed, each globular head domain within the dimer is tilted 60° relative to each other, and contains N-linked glycan moieties at positions N200 and N215 (the dimer interface) and position N416.^26,32^ We observed intrinsic flexibility in loops containing residues 283 (8.8 Å), 421 (5.2 Å), 534 (4.5 Å), and 589 (3.9 Å), consistent with prior structural data.^25,26,32,79^

#### Anti-H Antibody 1C08

We obtained a high-resolution structure of the H_ECT_ in complex with potently neutralizing and highly protective mAb 1C08 (3.2 Å), epitope group HE-4 (cyan, Fig 5A-B, E). MAb 1C08 anchors to the sides of the H dimer, binding symmetrically to the peripheral regions of each β-propeller, approaching the dimer from the membrane-proximal side. 1C08-H binding is predominantly mediated by the mAb heavy chain, with four residues in CDRs H1-3 engaging five residues in H blades β2 and β3 through hydrogen bonds and one salt bridge (H D347 - 1C08 R30). The mAb light chain contributes two CDRL1 residues to the interaction that are hydrogen bonded to two residues in blade β3 of H (Fig 5E). Structural comparison of the H dimer pre- and post-1C08 binding (using reference apo structure 2RKC,^79^ which exhibited the lowest global RMSD across all available H structures) revealed limited changes in the H dimer structure, with a minor, 1.3 Å conformational shift in a loop containing residue 311 away from the antibody paratope.

#### Anti-H Antibody 4D08

MAb 4D08, epitope group HE-1b, is also potently neutralizing *in vitro* and highly protective *in vivo*. While the mAb 1C08 complex described above was determined alone, structural analysis of the 4D08-bound complex was enabled via co-binding with 1C08, which mitigated preferred orientation observed with 4D08 alone and allowed high-resolution reconstruction (3.3Å) (Fig.5 A-B, F). 4D08 symmetrically engages epitope HE-1b on each monomer of the dimer, which is located proximal to the dimer interface on the apical surface. CDR H3 contributes most to the interaction, with a flattened loop of residues from 103-109 occupying a crevice formed by blades β1 and β5 (Figure 5F), much like the interaction mediated by SLAMF1 β strand residues 127-133.^25,32^ The 4D08 CDR L2 forms one hydrogen bond with H β6 residue T607.

Binding of 4D08 induces structural remodeling in H propeller blades β5 and β6, consisting of a two-residue register shift propagating from S544 to I564 (a 20-residue region containing blade β5, strand β4 and two unstructured loops on either side) (Fig S6A-C). 4D08 recognition of H B5 loop P545-R547 extends the loop ∼3.5 Å outwards, which pulls residues S544-P545 out of register (Fig S6D). The surrounding β strands accommodate this shift, likely due to sequence similarity between the unshifted and +2-shifted regions, allowing for local reorganization of the H fold.

#### Anti-H Antibody 1C02

A complex of H and mAb 1C02 (HE-1a) was also resolved by cryo-EM to 3.1 Å resolution (Fig 5C-D,G). Like 4D08 (HE-1b), 1C02 also symmetrically occupies this epitope on each monomer, on the apical surface near the H dimer interface (overlapping with HE-1b mAb 4D08). The 1C02 heavy chain forms ten hydrogen bonds with eight residues of H in blade β5, predominantly through the CDR H3, while the light chain CDRs L1 and 3 engage six H residues in blades β5 and β1 via nine hydrogen bonds. Of the fourteen H residues contacted, six also bind to SLAMF (D507, S532, R533, F552, Y553, and R556), one binds to nectin-4 (F549), and five interact with combination of SLAMF1, Nectin-4, and/or CD46 (D505, Y541, Y543, S548, and S550).^32^

Similar to 4D08, 1C02 also induces pronounced conformational changes in H protein blade β5 (Fig S6C, center right). These include the same two-residue register shift within blade β5 strand β4 (residues 546–555), as well as other remodeling. In the 1C02 complex, six loops designated by residues Q248, F284, D342, K460, V534, and S590 adopt alternate conformations relative to the apo structure. Additionally, the antibody loop containing S117 (CDR L3) introduces steric hindrance with H_ECT_, causing structural rearrangement of the N- and C-terminal β sheets of the β-propeller fold (blades β1 and β6, residues S189–S199 and H593–C606). The antibody-induced register shift in H observed in both complexes with 4D08 and 1C02 mirrors conformational changes previously reported in SLAMF1/CD150-bound H structures (PDB IDs: 3ALW, 3ALX, 3ALZ) (Fig S6C, right), but absent in the Nectin-4-bound (PDB ID: 4GJT) and CD46-bound (PDB ID: 3INB) conformations. These findings imply that 1C02 and 4D08 may mimic receptor-induced conformational states or were elicited against an H structure resembling that of the receptor-bound form.

The epitopes defined by 4D08 (HE-1b) and 1C02 (HE-1a) differ in the extent to which each mAb interacts with the receptor-binding groove (Fig 5H). The receptor-binding groove consists of an anterior wall composed mostly of blades β1 and β5, a posterior wall formed by blades β3 and β4, and a cavity formed by blades β4 and β5. MAb 1C02, which is less protective, more extensively interacts with the inside of this groove, contacting fifteen H residues in the anterior wall and throughout the inside of the cavity, five of which have been shown to be critical for SLAMF1 recognition (D505, D507, R533, F552, Y553).^80,81^ MAb 4D08, which is more protective, binds to fewer, more surface-accessible residues along the walls and edges of the cavity. All RBD contact residues for mAb 4D08 are also SLAMF1 contacts, including essential residue R533.

#### F structures

The cryo-EM structure of F complexed simultaneously to both 3A12 (FE-4) and 4F09 (FE-5) was resolved to 2.3 Å (Fig 6, Fig S5). Both 3A12 and 4F09 are attached at full occupancy, with three copies of each Fab per F trimer.

MAb 3A12 binds to the center of each protomer face, bridging the F1 and F2 subunits by engaging residues in the F1 HRN and DI and residues in F2 (Fig 6 A-B, D). The 3A12 HC and LC each form 11 hydrogen bonds with eight residues of F, mediated primarily by CDR H2 and CDRs L1 and L3. Binding of 3A12 causes a slight (3.0 Å) conformational shift in residues G250-G254, but otherwise the structure of 3A12-F_ECT_ and apo F_ECT_ are similar, with an RMSD of 0.7 Å.

MAb 4F09 (epitope FE-5) binds to the apex of F, with three Fab copies bound to the trimer, and each Fab simultaneously engaging two F protomers (Fig 6A-B, C). The 4F09 CDR H3 contains an eight amino acid tyrosine-rich (4 Tyr among 8 residues) loop from Y106 to Y113 that interacts with F_A_ and F_B_ residues in the F1 DIII domain, the F_B_ HR-N, and an F_B_ glycan in the F2 subunit (N61). This glycan also contacts framework FR H3 residue K67. For the 4F09 light chain, all contacts are limited to protomer F_A_. There, CDRs L1-3 bind to residues in F1 DIII and a glycan at F2 N66, and an additional contact is made between this glycan and residue S68 in the 4F09 FR H3 region.

#### Conservation of H epitopes

We assessed sequence conservation across H and F epitopes to evaluate the robustness of antibody binding in the context of natural viral diversity using a database of 2,350 sequences of H and 1,436 sequences of F (Fig 7, Table S2).

**Figure 7.**
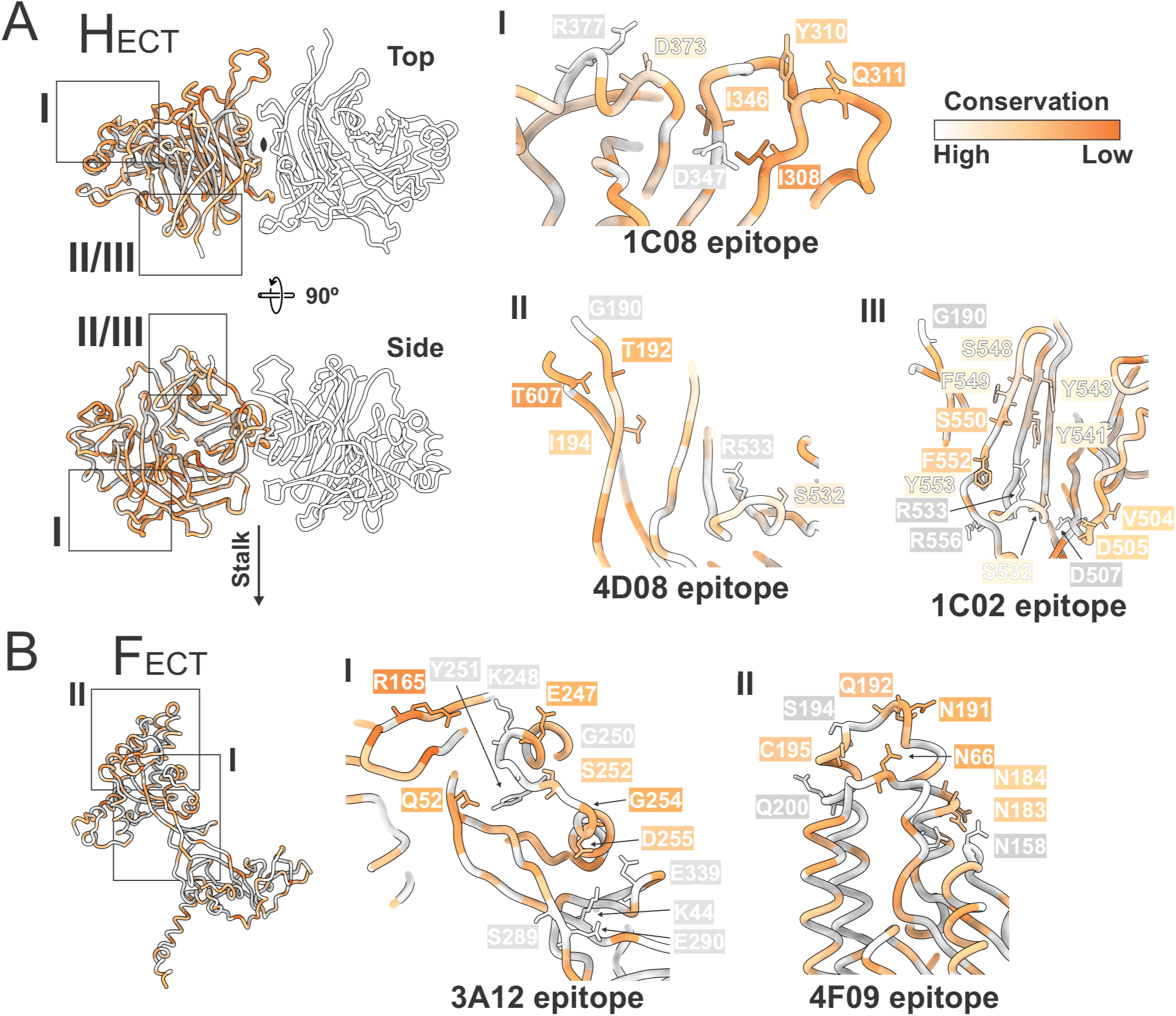
Epitope conservation mapping of measles virus H and F glycoproteins. Epitope residues are colored by conservation level (orange: low; white: high), based on multiple sequence alignments of 2,350 H and 1,436 F curated entries. Conservation scores were computed using AL2CO^142^ and visualized on representative PDB structures. Scores reflect both the frequency of each residue in the alignment and the presence of variants that would likely alter protein biochemistry at that position. Structural overviews in both panels indicate epitope regions for each antibody within the single chain or dimer context. **A)** Left: Ribbon model of the H dimer colored by per-residue conservation score derived from AL2CO, with orange indicating lower conservation and white indicating complete conservation. (I) the 1C08 epitope, composed primarily of conserved backbone-mediated contacts, with D373 as the only variant forming a side-chain hydrogen bond; most epitope residues are highly conserved. (II) The 4D08 epitope near the dimer interface, with residues T607 and I194 showing low conservation; the remaining epitope is conserved. (III) The 1C02 epitope exhibiting several variant residues, including D505, S532, Y541, and Y543, corresponding to naturally occurring point mutations in circulating strains. **B)** Left: Ribbon model of an F monomer colored by per-residue conservation score derived from AL2CO, with orange indicating lower conservation and white indicating complete conservation. (I, middle) Conservation profile across the 3A12 epitope on F, showing several moderately or poorly conserved residues, including R165, E247, S252, G254, and D255. (II, right): Close-up of the 4F09 epitope on F, highlighting residues N183 and N184 with low conservation.

The epitope of highly protective mAb 1C08 (HE-4) appears broadly conserved and likely robust against circulating variants (Fig 7A_I_). 1C08 forms key hydrogen bonds to side chains of fully or highly conserved residues Asp347, Asp373, and Arg377 (99.96-100% conserved), and backbone-only interactions to four moderate-to-highly conserved residues Ile308, Try310, Gln311, and Ile346 (84-99% conserved). Least-conserved residue Ile346 (84% conserved) is present as Val346 in circulating B3 strains, which would most likely not impact 1C08 recognition.

Highly protective mAb 4D08 (HE-1b) binds to a similarly conserved region on H, with S532 as the only residue with a variant (S532F) that might impact side chain-mediated interactions (Fig 7A_II_). Notably, this residue also directly contacts SLAMF1, and the variant at position 532 occurs only once out of 2,350 sequences.

Less-protective mAb 1C02 (HE-1a) binds to a larger epitope that, while highly conserved (98.26-100% conservation), may be more susceptible to escape mutations (Fig 7A_III_). Key contact residues with reported variants include Asp505 (D505G/Y), Ser532 (S532F), Tyr541 (Y541N), and Tyr543 (Y543H), which all form side chain hydrogen bonds with mAb residues. While these mutations are present in low frequency—each observed in only one to four sequences out of 2,305 sequences—they suggest a slightly higher susceptibility of 1C02 to escape mutations compared to protective mAbs 1C08 and 4D08. Variants S532F, Y541N, and Y543H were found in clinical isolates,^82–85^ while D505G/Y was found exclusively in vaccine or other lab-adapted strains.^86–88^

### Conservation of F Epitopes

Highly protective mAb 3A12 (FE-4, protomer face) contacts 16 residues, 15 of which are 99.4-100% identical across all 1,436 known sequences of F (Fig 7B_I_). The 16th residue is G254 which is 85.5% conserved among all historical sequences, but 100% conserved among currently circulating strains from genotypes B3 and D8. Highly protective mAb 4F09 (FE-5) contacts residues that are each 99.6-100% identical among all sequences (Fig 7B_II_).

In sum, we identified four neutralizing mAbs targeting distinct, highly conserved regions on MeV H and F proteins, which each demonstrate significant therapeutic potential *in vivo* against currently circulating MeV.

## Discussion

Two doses of the live-attenuated measles virus vaccine confer lifelong, durable protection against disease in 90-99% of vaccinees.^89,90^ We evaluated 15 MMR-vaccinated donors for their polyclonal MeV responses in order to identify individuals with vaccine-induced, protective, circulating antibodies. We observed small differences by age (stronger responses in older people) and larger differences by sex (stronger responses in all females). Previous findings vary, with some providing evidence of higher IgG titers in females,^91–93^ no sex-based difference,^94,9595–98^ or age-related sex-based differences in titers.^99^

One individual in our cohort, a 56-year-old female, who was vaccinated in childhood over 50 years prior to sample collection and boosted five years prior to sample collection, demonstrated the highest polyclonal response and the greatest numbers of H- and F-reactive memory B cells. Notably about 1% of her total circulating B cells were directed against MeV F, while ∼0.5% were against H, even five years after her last boost. This is the first study to directly quantify MeV H-versus F-specific memory B cells, but previous studies have shown that human polyclonal serum contains similar levels of MeV H-versus F-specific IgG.^40,100^

This individual’s PBMCs were used for mAb discovery using an optofluidic strategy and recombinant, fully glycosylated H and F ectodomain (H_ECT_ and F_ECT_) antigens. Hundreds of B cell clones against H_ECT_ and F_ECT_ were identified and used to engineer, express, and purify approximately 50 mAbs per antigen. MAbs were sorted into competition groups and assigned to epitopes HE1-4 or FE1-5.

### MeV H epitopes

Overall, we find that mAbs against the receptor-binding groove (epitope groups HE-1 and HE-2) universally neutralize (five of five neutralize with an EC_50_ of at least 0.176 µg/mL). However, another mAb, 1C08 which recognizes the peripheral ends of H (epitope group HE-4) and not the receptor-binding site also neutralizes potently: EC_50_ of 0.013 µg/mL.

Anti-H mAbs targeting the receptor-binding site (epitopes HE-1b) and 1C08 against the periphery (HE-4) protect against wild-type MeV B3 in the cotton rat model, reducing titers in the lung 50-fold (mAb 4D08) to 160-fold (mAb 1C08). Similarly to clinical viral pathogenesis in humans, wild-type MeV in cotton rats replicates in the lower and upper respiratory tracts and the lungs and spreads to lymphoid organs, mediated by SLAMF1 interactions.^101^

mAb 4D08 recognizes conserved residues of H also essential for SLAMF1 receptor binding. Interestingly, binding of both mAbs 4D08 and 1C02 (HE-1) induces conformational changes in H, including register shifts, that are similar to those induced by binding of SLAMF1 itself.^25^

The position of 1C08 at the periphery of H, combined with cryo-ET modeling of the H-F interaction of the related human parainfluenza virus 3,^102^ suggest that 1C08 may function by preventing interactions of MeV H and F. This supposition is not without precedent: some early studies described murine mAbs that pulled down H and blocked hemolysis, but didn’t block H-mediated red blood cell aggregation (through NHP CD46).^31,103^ Residues in the epitope of 1C08 are also highly conserved, particularly in currently circulating MeV genotypes. Importantly, no H contact residue identified for 1C08 or 4D08 has been named as a positive selection site in MeV H evolution.^104,105^

### MeV F epitopes

F epitopes, consisting of clusters FE-1 through FE-5, encompass most of the trimer surface, from the membrane-proximal pyramid base to the trimer apex. MAbs targeting the trimer’s upper half and apex (FE-3, 4, and 5), universally neutralize, with an EC_50_ of at least 0.118 µg/mL. Those against the bottom half (epitope groups FE-1 and 2) neutralize rarely or poorly. The most protective anti-F antibodies in this study, 4F09 (group FE-5) and 3A12 (group FE-4), recognize conformational and quaternary epitopes and binding would likely block transition of F to the postfusion form. 4F09, for example, the most protective mAb in this study, bridges two monomers together in the prefusion conformation, anchoring the F1 and F2 subunits together with the major HRN heptad repeat, a portion of F which would otherwise drive the conformational transitions of membrane fusion.

Notably, in this study, we find abundant antibodies against F (twice as many as against H), and that mAbs against F both neutralize potently and fully protect in the cotton rat model when delivered individually. One anti-F antibody, 4F09, was the most effective of any mAb in this study and reduced viral titers to undetectable levels (although interestingly was not the most neutralizing). These findings contrast with the previously accepted notion that neutralization is associated mainly with H and poorly with F. A major difference between this study and prior work, however, is that here, we specifically selected B cells using F containing substitutions that support maintenance of the pre-fusion conformation,^35^ whereas earlier work used depletion of F antibodies using vaccine F.^55,69,103,106–115,57,58^ Our use of pre-fusion F to identify antibodies likely favored identification of neutralizing antibodies, and was certainly critical to identification of pre-fusion specific antibodies that neutralize by preventing the conformational rearrangements associated with membrane fusion.

Additionally, the mAbs in our study are from one individual with high circulating anti-MeV-H/F IgG, and may not be representative of the average response to vaccination.

### Comparing these antibodies to previous work

Now that we have 3D structural information for the landscape of H and F recognition, we can reconcile studies performed prior to the availability of antibody structures (Fig S7). Previous work used pepscan^56^ and escape mutations^41,44,49,52^ to identify residues important for binding of antibodies against H and F elicited by immunization of mice. For H, residues previously clustered into groups III and the ‘noose’ generally map to epitope groups HE-1a/1b and HE-2a/b, respectively, while residues in murine group Ia map to epitope group HE-4 (Fig S7A). For F, previously implicated residue R73^70^ maps to human epitope footprint FE-3, residues G240-258^116^ to FE-4, and residues A388 and C420^7111768,118^ to FE-1a/b (Fig S7B).

We can also compare the epitopes observed here for MeV with those thus far known for related viruses, including the henipaviruses Nipah (NiV) and Hendra (HeV) and human parainfluenza virus 3 PIV3 (Fig. S7C). The PIV3 attachment protein, termed HN, recognizes its primary receptor via two sites, one of which is in a similar location on HN as MeV-H.^119,120^ Accordingly, the epitope of anti-PIV3 neutralizing mAb rPIV3-28^121^ is similar to those of mAbs 1C02 and 4D08, which target the receptor-binding site of MeV H. However, rPIV3-28 does not cause the same conformational changes and register shift upon binding to HN. Other mAbs against the attachment proteins also find neutralizing epitopes outside receptor-binding regions, including the dimer interface (like HENV32,^122^ n425,^123^ and PIV3HN-13^124^) and opposing ends of the dimer (like HE-4 for MeV and mAb hAh1.3 for HeV), emphasizing the possible significance of these regions in the less-understood fusion triggering process for the paramyxovirus family. In contrast, the henipavirus attachment protein achieves receptor binding in a different location, not at the side of the H beta propeller as in MeV, but instead, at the center of the beta propeller.^119^ This site also serves as a second receptor binding site for PIV3 HN^120^ and is targeted by several neutralizing henipavirus- and PIV3-specific mAbs, including m102.3,^125^ 41-6,^126^ 1E5,^127^ HENV 26,^122^ rPIV3-23,^121^ PIV3HN-05, PIV3HN-13.^124^ No mAbs against this site are yet known for MeV.

Previous work has identified three main antigenic regions on the prefusion F protein for other paramyxoviruses, including the apex and lateral face of the trimer, which together contain the fusion machinery, and the trimer base.^128^ Multiple neutralizing apex-directed antibodies have been identified,^121,128–132^ despite conserved apical glycosylation in MeV and henipaviruses.^129,130^ Apex mAbs are thought to stabilize the metastable prefusion conformation by sterically hindering the refolding of the heptad repeat N (HRN) domain and potentially disrupting the F-attachment protein interaction, and include 3D04 (group FE-3) and 4F09 (group FE-5) for MeV, mAbs 4H3,^128^ 66,^133^ and 92^130^ for NiV,^130,133^ and PIV-18 for PIV3.^121^ The lateral face of the head domain is largely unglycosylated and exposed, and undergoes extensive conformational rearrangement in the pre- to post-fusion transition.^35^ Antibodies against the lateral face are believed to lock the F protein in its prefusion state, thereby aborting the fusion process. Lateral face antibodies include measles group FE-4 (mAb 3A12; this study) as well as NiV mAb 5B3^134^ and HeV mAb 1F5,^129^ among others.^128,135,136^ Finally, the base of the F protein, epitope groups FE-1 and FE-2, presents a more complex antigenic site where neutralization is highly epitope-dependent. While some antibodies to the outer surface of domain II are poorly neutralizing, other antibodies like mAb 77^35^ target the interface between the F1 and F2 subunits, and potently neutralize by preventing completion of the structural changes required for membrane fusion. Other mAbs against the base include 2D07, 2B11, and 2G05 for MeV (FE-1a, −1b, and −2; this study) and 1H1 and 2B12 for NIV.^128^

Compassionate use of mAb m102.4^137^ against HeV, and recent FDA-approval of nirsevimab^138^ and clesrovimab^139^ against the related respiratory syncytial virus, open the door to use of mAbs against measles as well. For measles virus, such use of recombinant human mAbs for pre- or post-exposure therapy is of high interest as measles cases rise and outbreaks become more frequent. These antibodies would be particularly useful in the immunocompromised and people not yet vaccinated or fully vaccinated, including very young children. Indeed, subacute sclerosing panencephalitis, a progressive and nearly always fatal complication of measles infection, occurs more often in children infected before age 2.^140^ Further, the same immunocompromised people for whom a live-attenuated vaccine like MMR is contraindicated, are the most vulnerable to severe infection and have no current options beyond reliance on herd immunity. A combination of monoclonal antibodies may finally provide a complementary protective or therapeutic option.

Although the MMR vaccine has been used in hundreds of millions of people since its introduction in 1963, and has saved an estimated 94 million lives, antibodies elicited by this vaccine in humans are still very poorly understood. The work presented here now maps the landscape of antibody recognition against measles virus in a human recipient of the globally successful MMR vaccine, and illuminates which recognition sites on both H and F antigens lead to *in vitro* neutralization and *in vivo* protection. Further, these human mAbs themselves, which recognize conserved sites and inhibit measles by complementary mechanisms, now provide a needed treatment option against the resurging global burden of measles virus infection. With measles cases surging globally, the absence of an effective therapeutic is no longer acceptable— it is an urgent, unmet medical need.

## Acknowledgments

The authors thank Ruben Diaz Avalos with cryo-EM facility and Cheryl Kim with the flow cytometry core of La Jolla Institute for Immunology for assistance with data collection; and the GHR Foundation and private donors for support of instrumentation. Molecular graphics were performed with UCSF ChimeraX, developed by the Resource for Biocomputing, Visualization, and Informatics at the University of California, San Francisco, with support from National Institutes of Health R01-GM129325 and the Office of Cyber Infrastructure and Computational Biology, National Institute of Allergy and Infectious Diseases.

## Author contributions

M.D.A.: Designed, cloned, expressed, and purified viral protein constructs; cloned, expressed, purified, and digested mAbs; performed PBMC cell-staining and cytometric data collection, memory B cell enrichment and single-cell Beacon analysis, mAb cloning and sequencing, and high-throughput SPR binding kinetics and epitope binning; performed *in vitro* virus neutralization experimentation and data analysis; prepared figures and wrote the manuscript. D.S.Z.: Designed, cloned, expressed, and purified viral protein constructs, performed mAb epitope prediction with alphafold3, recorded cryo-EM data and processed and solved structures, analyzed cryo-EM data and epitope conservation, prepared figures, and assisted in writing and revising the manuscript. G.N.: Cloned, expressed, purified, and digested mAbs; expressed and purified viral proteins, collected and processed H_ECT_-Fab and F_ECT_-Fab NS-EM data and built 3D reconstructions, screened H_ECT_-Fab and F_ECT_-Fab complexes for cryo-EM, recorded cryo-EM data and processed and solved structures. S.S.H.: Performed donor serum reactivity screening; assisted with PBMC cell-staining and cytometric data collection, memory B cell enrichment and single-cell Beacon analysis, mAb cloning and sequencing, and high-throughput SPR binding kinetics and epitope binning. D.P.: Cloned, expressed, and purified mAbs and produced and purified Fabs. E.P.: Performed *in vivo* MeV protection experiments and data analysis. D.L.C.: Performed *in vitro* MeV experimentation and data analysis. S.N.: Performed *in vivo* MeV protection experiments and data analysis. M.P.: Performed *in vitro* MeV experimentation and data analysis, supplied reagents, provided supplies and assistance in establishing *in vitro* MeV assay protocols in the EOS lab, manuscript writing. K.M.K.: Directed single-cell Beacon analysis, mAb cloning and sequencing, and high-throughput SPR binding kinetics and epitope binning and data analysis, manuscript writing. E.O.S.: Project conception and direction, results analysis and interpretation, supervision, funding, manuscript writing.

## Data and materials availability

Materials are available upon request to. All structural data supporting the findings of this study have been deposited in the Electron Microscopy Data Bank (EMDB) and the Protein Data Bank (PDB). Negative-stain electron microscopy (EM) maps of the Measles virus fusion (F) glycoprotein ectodomain in complex with neutralizing antibodies are available under the following EMDB accession numbers: EMD-72182 (F + 3A12), EMD-72181 (F + 2G05), EMD-72184 (F + 4F09), EMD-72180 (F + 2D07), and EMD-72179 (F + 2B11). Negative-stain EM maps of the Measles virus hemagglutinin (H) glycoprotein ectodomain in complex with neutralizing antibodies have been deposited under the following accession numbers: EMD-72195 (H + 1C02), EMD-72196 (H + 1C08), EMD-72197 (H + 1G01), EMD-72198 (H + 1G03), EMD-72201 (H + 4D04), and EMD-72203 (H + 4D08 and 1C08). Cryo-EM structures of the Measles virus glycoprotein ectodomains in complex with neutralizing antibodies have been deposited in the PDB and EMDB under the following paired accession numbers: 9Q26 / EMD-72151 (F + 4F09 and 3A12), 9Q20 / EMD-72147 (H + 1C08), 9Q24 / EMD-72149 (H + 1C08 and 4D08), and 9Q1Z / EMD-72146 (H + 1C02). All data are publicly available and can be accessed through the EMDB (https://www.ebi.ac.uk/emdb) and PDB (https://www.rcsb.org) repositories.

## Supplementary Figures

**Figure S1. H- and F-specific mAb sequence analysis, kinetics, and MeV inhibition.** A) MAb variable heavy (VH) chains from vaccinee #3920 were sequenced, aligned, and arranged into phylogenetic trees to visualize mAb evolution. Horizontal branch lengths represent the estimated fraction of residue changes between mAbs. MAbs were produced in small volumes in expiCHO cells, and mAb-containing supernatants were assessed for antigen affinity and MeV inhibition *in vitro*; these data are shown in heat maps to the right of each mAb name. MAb VH gene designations are also listed for reference. H- and F-specific mAbs with the highest antigen affinities (▴, lowest KD values) or MeV inhibition abilities (*, highest % inhibition values) are denoted. B) MAb VH sequence mutations and CDR H3 lengths were quantified, and evaluated for correlation to antigen affinity or virus inhibition. Antigen affinity vs virus inhibition was also examined. Spearman nonparametric correlation coefficients (r) and P-values are listed above each graph. MAbs are color-coded according to the VH gene family (IGHV1= purple, IGHV 2= orange, IGHV 3= red, IGHV4= green, IGHV5= blue). ***P<0.0005.

**Figure S2. Approximating Fab-H/F interfaces using densities generated with NS-EM.** A) H_ECT_ and 1C02 Fabs, B) H_ECT_ and 4D08/1C08 Fabs, C) H_ECT_ and 1G01 Fabs, D) H_ECT_ and 4D04 IgG, E) H_ECT_ and 1G03 Fabs, F) H_ECT_ and 1C08 Fabs, G) F_ECT_ and 2D07 Fabs, H) F_ECT_ and 2G05 Fabs, I) F_ECT_ and 2B11 Fabs, J) F_ECT_ and 3D04 IgG, K) F_ECT_ and 3A12 Fabs, L) F_ECT_ and 4F09 Fabs.

I) NS-EM was performed on Fab-antigen complexes, and single particles extracted from the resulting micrographs were sorted into 2D classes in cryoSPARC.

II) Shown is the Gold Standard Fourier Shell Correlation (GSFSC) plot for each mAb-antigen 3D model. Each plot displays correlation values between two half maps in the final refinement used for complex reconstruction when none (blue), loose (green), tight (red), and corrected (purple) masks are applied to complex volumes. Masks reflect the degree to which noise is removed from regions surrounding single particles in an extraction box. The GSFSC resolution of the structure is the value at which the corrected mask curve crosses the 0.143 threshold (black line).

III) The viewing direction plot shows the number of particles with a viewing direction at a particular elevation/azimuth (equivalent to longitude) bin, and provides a general representation of particle orientation diversity.

IV) Visual rendering of fab footprints on H_ECT_ or F_ECT_. Left and center left panels: fab-antigen complexes were modeled by rigid-body fitting of H_ECT_, F_ECT_, and fab atomic models into cryoSPARC densities using ChimeraX. Center right and right panels: H_ECT_, F_ECT_, and fab surfaces were depicted using molecular maps (16 Å resolution, 2Å pixel separation) of the docked atomic models. Antigen surfaces within 12 Å of a Fab atomic model were colored with the fabs corresponding color to visualize Fab footprints.

**Figure S3. Alphafold multimer modeling vs NS- and cryo-EM.** The AlphaFold3 multimer framework was used to generate 5 antibody-antigen complex predictions per mAb per H (A) or F (B) glycoprotein. Model predictions (depicted in orange for H and blue for F) were aligned and visually assessed for convergence of antibody binding sites. Confidence in interface prediction was inferred from the consistency of binding location across models and local prediction confidence scores (pLDDT).

**Figure S4. H and F mAb protection at 1 mg/mL and 5 mg/mL.** A) Cotton rats were injected intraperitoneally with each mAb (1mg/kg, 5 animals per mAb), PBS (3-4 animals), or pooled human IgG (20mg/kg, NT titer 640, 5 animals treated) 14 hours before infection. Animals were then inoculated intranasally with 2×10^5^ TCID_50_ MeV B3-GFP. Animal lungs were collected and homogenized 4 days after infection, and lung tissue supernatants were serially diluted and titrated on Vero-SLAMF1 cells. Shown are MeV titers in the lungs (TCID_50_log10/g lung tissue), compared to intra-group PBS controls. The dotted line represents the TCID_50_ titration assay threshold of detection (1.7 TCID_50_log10/g lung tissue). B) MeV fold-change in lung titer (TCID_50_log10/g lung tissue), normalized to intra-group PBS controls. We subtracted average PBS-treated lung titers (TCID_50_log10/g lung tissue) from mAb-treated lung titers (TCID_50_log10/g lung tissue) and show the average reduction of lung titers (TCID_50_log10/g lung tissue) for each mAb treatment. * p<0.05, **p<0.005, ***p<0.0005, ***p<0.0001 by Brown-Forsythe and Welch ANOVA test (or Mann-Whitney test for mAb 4F09).

**Figure S5. Cryo-EM data processing workflow for antibody-bound measles virus H and F glycoprotein complexes.** Overview of the cryo-EM processing pipelines for four datasets: H_ECT_-TS + 1C08 (far left), H_ECT_+ 1C02 (center left), H_ECT_-TS + 1C08 + 4D08 (center right), and F_ECT_2M + 3A12 + 4F09 (far right). The top row indicates the number of micrographs acquired (total and those used in the final analysis). The middle panels show representative 2D class averages obtained after particle picking and initial classification. Bottom panels show Gold-standard Fourier shell correlation (GSFSC) curves for each refined reconstruction, indicating final resolution estimates (3.25 Å, 3.05 Å, 3.13 Å, and 2.29 Å, respectively), and angular distribution heatmaps representing particle orientation distributions used during 3D reconstruction.

**Figure S6. Structural comparison of the β-strand register shift in the measles virus H glycoprotein induced by antibody and receptor binding.** A) Location of the β-strand 5, encompassing residues S544–564 (highlighted in pink) within the head domain of the H ectodomain, shown in the context of the dimer. B) Side view of a single H protomer with the shifted β-strand (residues 544–564) highlighted to illustrate its relative position within the fold. C) Structural overlay of the β-strand segment (residues 544–564, shown as sticks) across four structures: apo form (AlphaFold3 model closely resembling the 2RKC structure), bound to antibody 4D08, bound to antibody 1C02, and bound to the CD150 receptor (PDB: 3ALZ, receptor peptide shown in blue). The surrounding H surface is rendered to provide context. D) Linear representation of the β-strand segment in the apo (left) and 4D08-bound (right) conformations, with side chains shown and residue identities labeled. Dashed lines indicate corresponding alignment positions between the apo and shifted conformations, emphasizing the two-residue register shift observed upon antibody binding.

**Figure S7. Human mAb epitopes against MeV, compared to prior analysis.** Pepscan and escape mutation profiling for previous murine antibodies are mapped on H_ECT_ (A, left panel) and F_ECT_ (B, left panel) and compared to the direct human mAb epitope mapping performed in this study (middle and right panels. C, left) Approach angles of anti-MeV-H_ECT_ mAbs compared to those targeting paramyxovirus relatives Hendra virus G (HeV-G), Nipah virus G (NiV-G), and human parainfluenza virus HN (PIV3-HN), determined by aligning the globular head domains for each glycoprotein-Fab complex to MeV-H_ECT_. C, right) A table summarizing each Fab’s species of origin, target antigen, epitope on the MeV H_ECT_ dimer, and Fab-antigen complex source PDBs.

## Tables

**Table S1. Properties of MeV mAbs used to define human epitopes on H and F. ^a^**VH= heavy chain variable region, VL= light chain variable region. **^b^**MAb kinetics were determined by measuring mAb binding interactions with MeV H_ECT_ or F_ECT_ via surface plasmon resonance, and data was fit to a 1:1 Languir model to determine the Ka (on-rate), Kd (off-rate), and KD (equilibrium dissociation rate) for each mAb using the Kinetics software package (Carterra). **^c^**EC_50_= half-maximal effective dose. **^d^**The amino acid sequence of the complementarity-determining region-3 on the mAb heavy chain.

**Table S2. Summary of H- and F-mAb contact residues as determined by cryo-EM.** Shown are the interacting residues between H_ECT_ and protective mAbs 1C08 and 4D08, and less potent mAb 1C02, and F_ECT_ and protective mAbs 3A12 and 4F09. Contact residues for each mAb are separated into heavy chain (HC) and light chain (LC) residues. The first letter of each residue refers to the chain, followed by amino acid and position. Interaction types, atom interaction state (glycoprotein - antibody), and conservation scores are also provided. Conservation scores were calculated and normalized using AL2CO. The maximum score for a fully conserved residue is 1.48 for H residues and 0.83 for F residues.

**Table S3. Cryo-EM and model refinement statistics.**

## Materials and Methods

### Experimental model and subject details

The La Jolla Institute for Immunology (LJI) Clinical Core recruited healthy adults for pre- and post-SARS-CoV2 vaccination blood donations and collected blood draws under IR-B approved protocols (IB-233-0820 A3). All donors gave informed consent at the time of study enrollment and provided information on gender, ethnicity, age, and MMR vaccination history (including a date approximation). Vaccinated subjects did not report exposure to authentic measles virus, although we cannot rule out unknown, subclinical infections. Pre-SARS-CoV-2 vaccine serum and PBMC samples were used in this study. Samples were coded, and then de-identified prior to analysis.

### Plasmids

We used full-length Measles virus (MeV) hemagglutinin (H) and fusion (F) constructs for donor serum IgG screens, and ectodomain constructs for all downstream experiments and analyses. All MeV-H and -F constructs were from strain IC323 (derived from the highly virulent wild type Ichinose-B95a strain rescued in B95a cells).^143^ Full-length MeV-H was codon optimized, synthesized, and sub-cloned into the mammalian expression vector pCAGGS^35^ (Epochlifescience) with a C-terminal 6x histidine tag. The H ectodomain (H_ECT_, residues 149-617) was subcloned to a pCG vector using the NEB High Fidelity Phusion polymerase and Gibson Assembly Master Mix (NEB). The pCG_H_ECT_ gene was flanked by a 5’ Ig Kappa leader sequence and a C-terminal twin StrepII tag. We also cloned an untagged version of pCG-H_ECT_ for structural studies. The full-length MeV fusion protein (F) and ectodomain (F_ECT_, residues 1-494) from MeV strain Ichinose-B95a were cloned into pMT-puro or pCG expression vectors. All F constructs contained two (“2M”: E170G and E455G) stabilizing mutations, which maintain F in the pre-fusion conformation, and a C-terminal twin Strep-TagII. The pMT-puro plasmid encoding F_ECT_ 2M was synthesized as previously described.^35^

### Cells

Vero (African green monkey kidney), and Vero-SLAM cells (Vero cells stably expressing MeV receptor human SLAMF1) were grown in Dulbecco’s modified Eagle’s medium (DMEM; Life Technologies; Thermo Fisher Scientific) supplemented with 10% (vol/vol) fetal bovine serum (FBS, Life Technologies; Thermo Fisher Scientific) and 1% penicillin-streptomycin (pen/strep, 100 U/mL penicillin and 100 μg/mL streptomycin) at 37°C and 5% CO_2_. Vero-SLAM cell culture medium also included 1 mg/ml geneticin (Thermo Fisher Scientific). ExpiCHO cells were maintained in ExpiCHO Expression Medium (Gibco) supplemented with 8 mM L-glutamine in a humidified incubator at 37°C with 8% CO_2_. HEK 293F cells were maintained in FreeStyle293 Expression Medium (Gibco) at 37°C with 5% CO_2_. *Drosophila* S2 cells were cultured at 25°C (no CO_2_) in complete Schneider’s *Drosophila* Medium (Gibco) with 10% heat-inactivated FBS and 1% pen/strep or in serum-free Insect Xpress Medium (Lonza).

### MeV Antigen Production and Purification

#### Full-length H purification via 6x his tag

HEK293F cells were expanded to a density of approximately 1 × 10^6^ cells/mL in at least 500 mL media on the day of transfection. Transfection was performed using 1μg/mL pCAGGS-H-F_FL_-6xhis and polyethylenimine (PEI, Polysciences) at a ratio of 1:3 DNA:PEI. The cells were cultured for five days, then centrifuged at 4000 x g and 10°C. The cell pellet was resuspended in a volume equivalent to one-tenth of the initial culture (50 mL) in ice-cold HBS buffer (50 mM HEPES, pH 8.0, 150 mM NaCl). Protease inhibitor cocktail (200 μL) and 1.5 μL of Benzonaze were added to the suspension, and the mixture was kept on ice. The suspension was then homogenized gently via a douncer. NP-40 (ThermoFisher) was added to the homogenized cells at a final concentration of 2%. The cells were incubated at 4°C for 2 hours with gentle rocking at approximately 1 Hz. The solubilized solution was centrifugated at 38,000 x g for 30 min at 4°C. The supernatant was incubated overnight with 1 mL of equilibrated NiNTA beads (Qiagen) at 4°C with rocking. This mixture was then diluted with HBS to reduce the final NP-40 concentration. The resin was washed with 0.06% NP-40 HBS, 0.06% NP-40 HBS with 20 mM imidazole (Fisher Scientific), then 0.06% NP-40 HBS with 50 mM imidazole, followed by elution with 0.06% NP-40 HBS 250 mM imidazole. The eluted medium was mixed with amphipol (Anatrace) (5 mg amphipol to 1 mg protein) for 20 to 30 min, then mixed with Bio-Beads (Bio-rad) to remove NP-40. Biobeads were removed using gravity filtration and the imidazole was removed by applying the filtered medium to a Hiload 16/600 superdex column (Cytiva) equilibrated with HBS buffer on an ÄKTA Pure system. Purified protein was flash-frozen in liquid nitrogen. Protein concentration was estimated using the molar extinction coefficient at 280 nm (97,750 M-1 cm-1).

#### Full-length F purification via Twin StrepII tag

Expression of full-length MeV-F in HEK293F cells and supernatant harvesting was performed as mentioned above using the pCG-F-FL-2M plasmid. Supernatant from homogenized cells was incubated overnight with 1 mL of equilibrated Streptactin resin (Qiagen) at 4°C with rocking. This mixture was then diluted with HBSE (50 mM HEPES ph8, 150 mM NaCl, 1mM EDTA) to reduce the final NP-40 concentration, then washed with 10 mL HBSE containing 0.06% NP-40. Amphipol was added to the resin and incubated for 2 h at 4°C with rocking. The resin was then washed three times with 2 mL HBSE, and protein was eluted with HBSE containing 5 mM d-desthiobiotin. Purified protein was flash-frozen in liquid nitrogen. Protein concentration was estimated using the molar extinction coefficient at 280 nm (57,700 M-1 cm-1).

#### H ectodomain purification via Twin StrepII tag

ExpiCHO cells were transiently transfected with plasmid encoding the H ectodomain construct (pCG_IgK_H_ECT__TwinStrep) using the ExpiFectamine CHO Transfection Kit (Gibco) in accordance with the manufacturer’s “high titer” protocol. Following expression, cells were removed by centrifugation at 4,000 × g for 15 min. The clarified supernatant was incubated with 300 μL Biolock (IBA GmbH) and the pH was adjusted to 8.0 with 1 M HEPES (final concentration 50 mM). After 30 min, the medium was further clarified by centrifugation at 4,000 × g for 10 min and sterile-filtered using a 0.22 μm filter. The filtered medium was applied to a Strep-Tactin HP column (Cytiva) equilibrated with HBS buffer (20 mM HEPES, pH 8.0, 150 mM NaCl) on an ÄKTA Pure system. After extensive washing with HBS buffer to remove nonspecifically bound proteins, tagged H_ECT_ was eluted using 5 mM desthiobiotin (Sigma) in HBS. Fractions containing the protein of interest were pooled, concentrated to 0.8 mg/mL, and flash-frozen in liquid nitrogen. Protein concentration was estimated using the molar extinction coefficient at 280 nm (84,800 M-1 cm-1).

#### H ectodomain purification via ion exchange

Due to potential conformational constraints imposed by the twin-strep tag in close proximity to the stalk region, an untagged H_ECT_ variant was also expressed and purified. Following removal of cells by centrifugation, the supernatant was adjusted to pH 9.0 using 1 M Bis-Tris-Propane (Sigma) based on the calculated isoelectric point (pI ≈ 6.6) for H_ECT_. The medium was filtered through a 0.22 μm membrane and loaded onto a 6 mL Resource Q anion exchange column (Cytiva). Protein was eluted using a 10-column volume linear gradient up to 100% of elution buffer. Eluted fractions containing Hecto were concentrated and subjected to size-exclusion chromatography using a Superdex 200 Increase 10/300 GL column (Cytiva). Fractions corresponding to the H_ECT_ peak were pooled, concentrated to ∼0.4 mg/ml, and flash-frozen.

#### F ectodomain purification via Twin StrepII tag

F ectodomain was produced in Drosophila S2 cells as previously described.^35^ To summarize, cells were seeded in a six well plate in complete Schneider’s medium, then transfected with a F ectodomain plasmid the following day using the Effectene (Qiagen) transfection protocol. Cells were transferred to a T25 flask and cultured with puromycin (6 μg/ml) on day 5, then expanded to a T75 flask and cultured with Insect Cell Expression Culture Media (Lonza) with 6 μg/ml puromycin. Cells were expanded to 2 L shaking flasks, induced with 500 μM CuSO_4_ at 1 × 10^7^ cells/mL, and harvested four days after induction. Cells cleared via centrifugation at 5000 x *g* for 20 min. The supernatant was adjusted to pH 8.0 with 1 M HEPES, and incubated with Biolock (IBA) and NaCl (final concentration 800 mM). The supernatant was loaded onto a pre-washed StrepTrap HP 5 mL column (Cytiva). Purified F ectodomain protein was eluted with high-salt HBS (20mM HEPES, 800 mM NaCl) containing 5 mM d-desthiobiotin. The purified protein buffer was exchanged for high-salt HBS-only buffer using a HiPrep 26/10 desalting column (Cytiva). Purified protein was flash-frozen in liquid nitrogen. Protein concentration was estimated using the molar extinction coefficient at 280 nm (48,820 M-1 cm-1).

#### H_ECT_ and F_ECT_ labeling

We used biotinylated or fluorescently-labeled H/F constructs for antigen-specific memory B cell enrichment, antigen-specific PMBC quantification via flow cytometry, and single-cell memory B cell identification on the Bruker Cell Analysis Beacon. Non-specific biotinylations were performed using EZ Link NHS-biotin (20217, Thermo Scientific) according to the manufacturers protocol. Proteins were incubated with NHS-biotin for 2 hours at 4°C while gently shaking. Reactions were quenched with 50uL Tris HCl (1M, pH 8) for 10 min. For fluorescent detection, proteins were labeled with succinimidyl ester (NHS)-linked alexa fluor 555 (NHS-AF555, A20009, Thermo Fisher Scientific) or alexa fluor 647 (NHS-AF647, A20006, Thermo Fisher Scientific) according to the manufacturer’s protocol. Proteins were labeled in 0.1M sodium bicarbonate at 4°C with periodic vortexing. Biotinylated and fluorescent proteins were desalted using PD G-25 miditraps (Cytive, 28918008) into HBS, then aliquoted, flash-frozen in liquid nitrogen, and stored at −80°C. Protein concentration was estimated using the molar extinction coefficient at 280 nm (84,800 M-1 cm-1 for H_ECT_, 48,820 M-1 cm-1 for F_ECT_).

### Virus

Recombinant Measles virus strain B3 encoding the mCherry reporter gene (MeV-mcherry) was obtained via viral rescue. Briefly, HEK293T cells were transfected using Lipofectamine 2000 with cDNA constructs encoding the T7 polymerase (GS58929), MeV-mCherry genome (GS68838, with mCherry), and measles proteins N (GS6558900-1), P (GS59482), and L (GS58944) (MeV proteins sourced by Epoch Life Sciences). Cells were incubated overnight at 37°C in Opti-MEM medium, then the medium was replaced with DMEM 10% FBS and pen/strep. Cells were heat-shocked by incubating them in a 42°C water bath for 3 hours, then returned to 37°C for 48 hours. Next, transfected HEK293T cells were detached and overlaid with Vero-SLAM cells in T-75 flasks to facilitate rescued virus amplification. The virus underwent four passages and was titrated to a concentration of 10^6^ plaque-forming units per milliliter (PFU/ml). RNA from the virus sample was then sequenced and the virus stock used for subsequent *in vivo* experiments.

For virus stock propagation, Vero-SLAM cells were infected with 0.05 to 0.01 PFU/mL at 32°C in DMEM 10% FBS 1% pen/strep. When extensive cell-cell fusion was observed, the media was replaced with fresh DMEM 10% FBS 1% pen/strep, and cells were scraped into the new media. Cells were freeze-thawed at −80°C and 25°C twice to release cell-associated virus particles, then spun at 3000 to 4000 RPM. Media was replaced, and the resuspended cell pellet was aliquoted, and stored at −80°C.

Virus stock titers were determined using end-point dilutions and fluorescent cell quantification via the CellInsight CX5 HCS Platform (Thermo Fisher). Because cells infected with MeV form large multi-nucleated syncytia, automated cell quantification was simplified by treating infected cells with 500nM FIP-HRC, a F-specific peptide that inhibits MeV cell-cell spread and syncytia formation.^144^

### Serum ELISAs

Donor serum samples were heat-inactivated for 30 min at 55°C. Half-area high-binding 96 well plates were coated overnight with 200 ng/well of purified full-length MeV H and F at 4°C. Wells were blocked with 3% milk in PBS for 3 hours, followed by incubation with donor sera diluted 1:100, 1:1000, 1:10,000, and 1:100,000 in PBS with 0.05% Tween-20 (PBS-T) and 1% milk for 1 hour. Plates were washed five times with PBS-T, then incubated with goat anti-human IgG-HRP (Invitrogen 31413), diluted 1:7500 in 1% milk PBST for 1 hour. Plates were washed five times with PBS-T, then developed using 1-Step Ultra TMB-ELISA substrate (Thermo Fisher 34029). Reactions were stopped with 1M H_2_SO_4_, and plate signals at 450 nm were measured using a Tecan Spark 10M plate reader. Donor serum reactivities to MeV H and F individually and summer are displayed as heatmaps (red = higher reactivity, beige = lower reactivity).

### MeV Memory B cell detection via flow cytometry

Cryopreserved PBMCs from donors #3920, #0003, and #4168 were prepared and stained as previously described,^145^ with some modifications as follows: PBMCs were thawed at 37°C and then resuspended in prewarmed DMEM 10% FBS 1% pen/strep with Benzonase (Sigma 70746-4). Cell counts and viability after washing were assessed on a Countess II FL (Invitrogen). PBMCs were placed in U-bottom 96-well plates and stained with 200 ng H_ECT_-AF647 or F_ECT_-AF647 in Brilliant Stain Buffer (BSB, BD Biosciences cat#563794) for 1 hour at 4°C, protected from light. Cells were then washed in BSB and resuspended in a staining cocktail containing surface antibodies diluted in BSB for 30 min at 4°C in the dark. Viability staining was performed using LIVE/DEAD fixable aqua dead cell stain (Thermo cat#L34965), and cells were incubated at 4°C for 30 min. Cells were washed twice in PBS and resuspended in FACS sort buffer (0.5 M EDTA, 25 mM HEPES pH 8, 1% FBS, in Hank’s balanced salt solution [Thermo Fisher, cat#14175095]). The acquisition was performed using a FACSSymphony S6 cell sorter. The frequency of antigen-specific PMBCs was expressed as a percentage of total memory B cells (Live, Lymphocytes, Singlets, CD3^−^ CD14^−^ CD16^−^ CD56^−^CD8a^−^, CD19^+^ CD20^+^, CD20^+^ CD38^int/–^, IgD^−^ and/or CD27^+^).

### mAb Identification and Production

#### MemB Enrichement

Cryopreserved PBMCs were thawed and washed twice in DMEM 20% FBS and Benzonase (50 U/mL) via centrifugation (500g for 10 min). Cells were resuspended in DMEM 20% FBS and assessed for density and viability, filtered through mesh, pelleted, and resuspended in cold PBS (0.5% BSA, 2mM EDTA, pH ∼7.3). Cells were mixed with biotinylated H_ECT_ and F_ECT_ (2 ng/uL) for 30 min at 4°C, then washed and blocked with Human FC block (BD #564220) for 15 min at 4°C. Magnetic anti-biotin microbeads (Miltenyi Biotec #130-090-485) were then added to cells for 15 min at 4°C. Cells were washed and resuspended in PBS (0.5% BSA, 2mM EDTA, pH ∼7.3), then passed through magnetic MS columns (Miltenyi Biotec 130-042-201). Antigen-specific PBMCs bound to antigen-coated anti-biotin beads were eluted from the column. We performed two parallel PBMC enrichments, one with biotinylated H_ECT_ and F_ECT_ combined, and one with biotinylated F_ECT_ alone, as we previously observed the ability for H_ECT_ to bind to and introduce non-target toxic lymphocytes to our enriched cell populations. To mitigate this, pre-H+F-enriched cells were first treated with 1 mM L-Leucyl-L-Leucine methyl ester (LLME, Cayman Chemicals 16008) to eliminate cytotoxic cells.

#### MemB activation and Beacon Analysis

H_ECT_- and F_ECT_-enriched cells were activated and loaded onto a Bruker Cellular Analysis Beacon for single-cell identification as previously described.^146^ Enriched cells were overlaid on feeder mitomycin-treated MS40L cells in 6-well plates in complete Iscove’s medium (cIMEM, 20% Low IgG FBS, 1% pen/strep, 5 μg/mL insulin, and 50 μg/mL human transferrin) supplemented with 100 U/ml Hu rIL-2 (GenScript), and 100 ng/mL Hu rIL-21 (Stemcell Technologies). Seven days later, cell densities were readjusted to 250,000 cells/mL and culture media was changed to cIMEM supplemented with 500 U/mL Hu rIFN-α 2b (PBL Assay), 50 ng/mL Hu rIL-6 (Stemcell Technologies) and 10 ng/mL Hu rIL-15 (BioLegend). Three days later (10 days after enrichment), cells were loaded onto OptoSelect 11K chips (Bruker Cellular Analysis), wherein they were isolated as single cells in nanoliter pens using OEP light cages. Antigen-specific antibodies were detected in a 30-minute time course assay using anti-human IgG beads (amount, Spherotech cat#HUP5-60-5) that were resuspended in media containing H_ECT_-AF555 (6 μg/mL), F_ECT_-AF647 (6 μg/mL) and goat anti-human IgG-AF488 (1 μg/mL, Invitrogen cat# A11013).

#### mAb Cloning

Cellular RNA to cDNA synthesis was performed on-chip using the Bruker OptoSeq BCR kit, according to the manufacturer’s instructions and previously described.^146^ Total cDNA was amplified using the Platinum SuperFiII polymerase (Invitrogen), and cleaned up enzymatically. Antibody heavy chain (HC) and light chain (LC) variable domains were amplified with one or two rounds of nested PCR using the PlatinumII Hot-Start polymerase (Invitrogen) and previously published primer sets.^147^ PCR products were assessed using 96-well E-gells (ThermoFisher), and paired wells were sequenced and analyzed using the International ImMunoGeneTics Informations System (IMGT)/V-Que.^148,149^ Unique variable heavy (VH) and variable light (VL) domains were cloned into linearized antibody expression vectors (human IgG1 and relevant light chain) using Gibson Assembly (NEB) according to the manufacturer’s protocol. Ligation reactions were transformed into 5-alphaF’Iq competent E. coli cells (NEB). QIAprep 96 Turbo Miniprep Kits (Qiagen 27191) were used to purify plasmids. Plasmids were sequenced to ensure that genes were in-frame, and the cloned VH and VL domains were compared to PCR sequences.

#### mAb and Fab Production and Purification

Test expressions of each mAb were carried out in 2.5 mL cultures of ExpiCHO cells cultured in 24-well blocks. Cells were transiently transfected and incubated at 37°C, 5% CO_2_ for five days. Supernatants were clarified and either 1) used directly in preliminary kinetics and neutralization screens (Fig S1) or 2) applied to PreDictor MabSelect SuRe 20uL 96-well plates (Cytiva Cat# 28925824) to isolate mAbs. Purified mAbs were used for H_ECT_ and F_ECT_ epitope binning to further select mAbs of interest. A panel of sixteen mAbs was produced in larger (25-100 mL) expiCHO cell cultures and purified from cell supernatants using Praesto AP resin (Purolite PR00300-310AP) or a HiTRap Fibro PrismA column (Cytiva). MAbs were then dialyzed into PBS and stored at −20°C.

For structural analyses, purified mAbs were digested into antibody fragments (fabs) by incubating antibodies with papain (2.5 mg/mL) in the presence of 10mM L-cysteine for 3 hours at 37°C. Digestions were stopped by adding iodoacetamide (50mM). Fabs were separated from FC fragments using Praesto AP resin, then dialyzed into PBS and stored at −20°C. mAb and Fab concentrations were determined using the absorbance at 280nM and the general IgG extinction coefficient 210,000M^−1^ cm^−1^.

### High-throughput SPR binding kinetics

Binding kinetics measurements were performed on the Carterra Lodestar Array (LSA) platform using HC30M sensor chips (Carterra) at 25°C. Two microfluidic modules, a 96-channel print-head (96PH) and a single flow cell (SFC), were used to deliver samples onto the sensor chip. Sensor chips surfaces were conditioned using 25 mM MES pH 5.5 with 150 mM NaCl and 0.05% Tween 20 and activated with a freshly prepared solution of 130 mM 1-ethyl-3-(3-dimethylaminopropyl)carbodiimide hydrochloride (EDC) + 33 mM N-hydroxysulfosuccinimide (Sulfo-NHS) in 0.1 M MES pH 5.5 using the SFC. MeV mAbs were either captured from cell supernatant on to chips using goat anti-Human IgG Fc secondary antibody (50ug/mL, VWR, 103255–066) or purified, diluted to 5 (MeV-H mAbs) or 2.5 μg/mL (MeV-F mAbs) in 10 mM NaOAc (pH 4.5, 0.05% Tween 20), and directly immobilized on to chip surfaces for 20 min. All IgG immobilization steps were followed by a 7-min injection of 1 M ethanolamine-HCl (pH 8.5) using the SFC to quench unreactive esters.

A seven-fold dilution series of MeV-H_ECT_ or -F_ECT_ was prepared in 1xHBSTE-BSA (10 mM HEPES pH 7.4, 150 mM NaCl, 3 mM EDTA and 0.01% Tween 20) buffer. Each antigen was then injected onto the chip surface using the SFC, from the lowest to the highest concentration, without regeneration in between, using 1xHBSTE-BSA running buffer. Five to six injections of buffer before the lowest non-zero concentration were used for signal stabilization. For each concentration, baseline data were collected for 120 s, association data for 300 s and dissociation data for 900 s. After the titration of each analyte, the chip surface was regenerated with two pulses (20-30 s per pulse) of 10 mM Glycine, pH 2.0.

Titration data were processed with the Kinetics software package (Carterra), including reference subtraction, buffer subtraction and data smoothing. H_ECT_ or F_ECT_ binding time courses for each antibody were fitted to a 1:1 Langmuir model to derive k_a_, k_d_ and K_D_ values.

### High-throughput SPR epitope binning

Premix epitope binning was performed on the Carterra Lodestar Array (LSA) platform using HC30M sensor chips (Carterra) at 25°C in 1xHBSTE-BSA running buffer.^150^ The chip was activated as described above. Each mAb was immobilized on the chip’s surface and separately mixed with H_ECT_ or F_ECT_ at a 4x molar excess of IgG to antigen in 1xHBSTE-BSA. MAb-antigen solutions were injected over the chip’s surface for 5 min. Chip surfaces were regenerated after every cycle using single pulses (30 s) of 10 mM Glycine, pH 2.0. Data was processed and analyzed with Epitope software (Carterra).^150^ Briefly, data was referenced using unprinted locations on the array. The binding level of the analyte antibody-antigen complex was compared to that of an antigen-alone injection. Signals that were significantly increased relative to the antigen-alone controls are described as sandwiches and represent non-blocking behavior. Competition results were visualized as a heatmap that depicts blocking relationships of analyte/ligand pairs. Antibodies with similar patterns of competition are clustered together in a dendrogram and are assigned to shared communities.

### Epitope mapping using NS-EM

#### Fab-antigen complex formation, grid preparation, NS-EM, and 3D volume reconstruction

MeV H_ECT_ or F_ECT_ were incubated with an excess of fabs for 15-30 min at RT and applied to freshly glow-discharged, carbon-coated 400 mesh copper grids (C-flat, Electron Microscopy Sciences). Following a rapid water wash, grids were stained with 1% uranyl acetate for 1 min. Excess stain was removed and grids were air dried. TEM images were collected using SerialEM on a Titan Halo 300kV electron microscope using a Gatan K3 direct electron detector. Motion correction, CTF estimation, particle picking, 2D classification and 3D reconstruction were performed using cryoSPARC software algorithms. Reconstructions were evaluated using Gold Standard Fourier Shell Correlation (GSFSC) metrics to prevent overfitting and determine the final map resolution at an FSC value of 0.143.

#### Fab-antigen complex modeling

Fab-antigen complexes were modeled by rigid-body fitting of individual H_ECT_, F_ECT_ and fab models into cryoSPARC densities in ChimeraX. Atomic H_ECT_ and fab models were generated using AlphaFold2 and AlphaFold3 predictions. N-linked glycans (Man_5_GlcNAc_2_) were modeled on H_ECT_ residues N200, N215, and N416 using alphafold3.^26,32^ The cryo-EM structure of Fecto2M (PDB ID: 8UUP) was used as a starting model.^35^ Density handedness was determined via visual inspection of docked antigen models.

#### Fab-antigen epitope mapping

We generated 3D molecular map surfaces (molmaps) from the docked H_ECT_ or F_ECT_ atomic models at 16 Å using ChimeraX command (molmap #model 16 gridspacing 2) and rendered rough representations of fab interfaces by coloring regions of these surfaces that fell within 12 Å of docked fabs.

### AlphaFold3 Modeling of Antibody–Antigen Complexes

We generated antibody–antigen complex predictions for both the H and F glycoproteins in order to evaluate whether AlphaFold3 multimer modeling could accurately predict experimental NS and cryo-EM observations and support interface identification in the absence of high-resolution structural data. For the F glycoprotein, the ectodomain (residues 24–495) was modeled as a single continuous chain and replicated three times to model the trimeric complex. Similarly, two copies of the H ectodomain (residues 149–617) were used to model its dimeric architecture. Each antibody was modeled in complex using the variable regions (VH and VL) of the heavy and light chains as input. Predictions were generated using the AlphaFold3 multimer framework. To account for glycosylation, AlphaFold3-generated JSON files were generated using the UniProt2AlphaFold3 software (https://github.com/dzyla/AF3_JSON_generator), which annotated sequences with experimentally determined glycosylation sites annotated in the Uniprot database. Predicted models (five per antibody–antigen pair) were aligned and assessed for convergence of antibody binding sites (precision) and their agreement with experimental data (accuracy) using root-mean-square deviation (RMSD) analyses

Accuracy and precision RMSD values (A-RMSD/P-RMSD) were calculated by first defining centroids- or the XYZ coordinates-of each predicted and experimental fab model in ChimeraX. For A-RMSDs, we then quantified the average Euclidean distance from model centroids to the experimental centroid. For P-RMSDs, we quantified the average Euclidean distance from model centroids to a theoretical “centroid of centroids” coordinate, which reflects the spatial spread of the centroids around their collective center. Because the typical resolution achievable using NS-EM is approximately 20Å, we considered predicted models sets to be accurate/precise if the overall A-RMSD and P-RMSD values for each antigen-mAb set were below 20Å.

### *In vitro* viral neutralization

Confluent Vero-SLAMF1 cells were diluted 1:20 and seeded in 96-well plates (100 uL/well). The next day, mAbs and virus were combined and applied to cells for infection. MAbs were serially diluted 1:5 (final concentration range 2.56×10^−4^ to 20 ug/mL) and pre-incubated with 5×10^3^ FFU/mL MeV-mCherry for 1 hour at room temperature (RT). Vero-SLAM supernatants were removed, and mAb-virus mixtures were added to cells and incubated at 37C 5% CO2 for three hours. Because adherent cell lines infected with MeV fuse together to form giant multi-nucleated syncytia that are difficult for standard fluorescent detectors to define and quantify, we treated infected cells with FIP-HRC, a F-specific peptide that inhibits MeV cell-cell spread and syncytia formation.^144^ After three hours, cell infection media was replaced with media containing 500 nM FIP-HRC. Cells were then returned to the incubator for infections to proceed overnight. The next day, cells were fixed and stained using 3.7% formalin in PBS containing Hoescht (1:1000) for 10 minutes at RT. MCherry-infected cells were counted using a Cellinsight CX5 plate reader (ThermoFisher). The samples were tested in three to four experiments. The measurements from each plate were normalized using mAb^−^/MeV^+^ wells. Data were analyzed in GraphPad Prism 10 and are shown as normalized percent MeV inhibition. We used the nonlinear regression agonist vs normalized response (variable slope) model to determine half-maximal effective concentrations (EC50s) for each mAb.

### *In vivo* mAb prophylactic protection in cotton rats

Inbred cotton rats (*Sigmodon hispidus*, cotton rat strain “Hsd:Cotton Rat” without genetic modifications) were purchased from Envigo, Inc., Indianapolis. Male and female cotton rats aged 4 to 10 weeks were used for experiments. Isoflurane-anesthetized rats were injected intraperitoneally with each mAb (1mg/kg, 5 animals per mAb), PBS (3-4 animals), or human IgG (5 animals) purified from pooled sera (20mg/kg, Carimune, CSL Behring, NT titer 640) 14 hours before infection. Anesthetized rats were then inoculated intranasally with 100 μl 1×10^5^ TCID_50_ MeV B3-GFP. Four days after infection (when animals typically reach peak MeV titer in the lung tissue,^151^ animals were euthanized by CO_2_ inhalation, and their lungs were collected and homogenized. Lung tissue homogenates were serially diluted and titrated on Vero-SLAMF1 cells. Plates were scored microscopically for cytopathic effect (CPE) after 7 days, and the TCID_50_ was calculated. Average raw titers are shown in Fig S5A grouped in the chronological order animals were assessed. For Fig4B-D, we subtracted average PBS-treated lung titers (TCID_50_log10/g lung tissue) from mAb-treated lung titers (TCID_50_log10/g lung tissue) and show the average reduction of lung titers (TCID_50_log10/g lung tissue) for each mAb treatment. For both raw and normalized data, we used a one-way ANOVA (not assuming equal standard deviations) to compare mAb vs control (PBS) lung titer reductions (log10/g lung tissue), with the Dunnett’s multiple comparisons correction. Because lung titers for animals treated with mAb 4F09 all fell below the 10^2^ TCID_50_/g lung tissue threshold of detection, the nonparametric Mann-Whitney test was used. * p<0.05, **p<0.005, ***p<0.0005, ***p<0.0001.

### Sample Vitrification for Cryo-EM

Cryo-EM grids were prepared with the following sample combinations: H_ECT_-TS with 1C08 and H_ECT_ with 1C02 were applied to Quantifoil R2/2 copper grids coated with an in-house prepared graphene oxide film, following a previously established protocol.^35^ The H_ECT_-TS + 1C08 + 4D08 complex was vitrified on C-flat 2/1 grids, while the F_ECT_2M + 3A12 + 4F09 complex was applied to Quantifoil R2/1 grids, also coated with a graphene oxide layer.

For vitrification, protein complexes were diluted to a final absorbance of 0.1–0.2 at 280 nm. A 3 μL aliquot of each sample was applied to glow-discharged grids using a Vitrobot Mark IV (Thermo Fisher Scientific) operated at 4 °C and ∼100% relative humidity. After a 15-s incubation, grids were blotted for 1.5–3 s using blot force setting 0 and a drain time of 0.5 s. The grids were immediately plunge-frozen in liquid ethane, cooled by liquid nitrogen, and stored under cryogenic conditions until imaging.

### Data Acquisition, Processing, and Model Building

Cryo-EM data were collected on a FEI Titan Krios transmission electron microscope operated at a nominal magnification of 130,000×, equipped with a Gatan K3 direct electron detector and a 20 eV energy filter. The calibrated physical pixel size was 0.66 Å/pixel. All datasets were processed using cryoSPARC v4.6.2 (Structura Biotechnology Inc.). Initial particle picking was performed in live sessions using the blob picker in over-picking mode (particle diameter range: 80–120 Å; minimum distance: 0.5×diameter). Approximately 5 million particles were initially extracted per dataset and Fourier-binned to 64 pixels to accelerate fast classification steps. Particles corresponding to the target complex were identified using Deep2D classification.^35^

Subsequent rounds of non-uniform and heterogeneous refinement were performed to isolate high-quality particles, from which ∼10,000 particles were selected for training a Topaz particle picker (10 training epochs). Particles picked by Topaz were also binned to 64 pixels and subjected to iterative refinement and classification. High-quality particles were then re-extracted at 160 pixels for improved alignment, achieving a resolution limit of ∼5 Å, which enabled further discrimination of conformational heterogeneity via 3D classification.

A final round of particle re-extraction was performed with binning adjusted to produce a Fourier shell correlation (FSC) of 0.143 at approximately 75% of Nyquist resolution, corresponding to Fourier-cropped particle box sizes of 320–360 pixels. Final refinements used non-uniform refinement with symmetry: C2 for H_ECT_ complexes and C3 for Fecto2M. For H_ECT_ datasets, symmetry expansion (C2) was followed by 3D classification and local refinement.

Initial structural models were generated using AlphaFold2 and AlphaFold3 predictions for the H_ECT_ and Fab fragments. The cryo-EM structure of F_ECT_-2M (PDB ID: 8UUP) was used as a starting model. All initial models were rigid-body fitted into the density maps and further refined using ISOLDE. Manual adjustments were performed in Coot, and final real-space refinement was done in Phenix v2.0-rc.

### Structural Analysis

Antibody–protein interfaces were analyzed using the Protein Interfaces, Surfaces and Assemblies (PISA) tool within the CCP4 suite. Local root-mean-square deviation (RMSD) analyses were performed using a custom Python script. In this approach, structurally aligned chains from two models were superimposed using the Superimposer module from Biopython, and residue-resolved deviations were calculated based on Cα–Cα distances derived from the sequence alignment. This allowed quantification of localized conformational differences across distinct structural states.

The most representative H ectodomain structure was identified through complete pairwise structural comparisons and global RMSD calculations. The structure exhibiting the lowest average RMSD relative to all others was selected as the reference, corresponding to PDB ID: 2RKC.

To model the β-sheet register shift, a full-length H ectodomain model was generated using AlphaFold3. This model was then compared to the reference structure (2RKC) and compared to the determined cryo-EM structures of H in complex with antibodies or previously described X-ray structures.

### Sequence Conservation Analysis

Sequence conservation analysis for both H and F glycoproteins was performed using curated datasets obtained via the SPACE (https://github.com/dzyla/SPACE). Sequences were filtered against the respective reference entries from UniProt (H: Q786F2; F: Q786F3) and aligned using FAMSA (Fast and Accurate Multiple Sequence Alignment), a tool optimized for large datasets. H and F sequences in the dataset include naturally occurring and lab-adapted MeV strains.

Per-residue conservation scores were computed using the AL2CO program.^142^ First, amino acid frequencies at each position were estimated using the “estimated independent counts method,” and a conservation index was calculated using the frequencies and the “entropy-based measure strategy” formula:

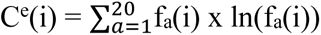

C^e^(i) = entropy-based conservation score at position i
f_a_(i) = frequency of amino acid a at position i in the multiple sequence alignment

Conservation indices were normalized using the mean conservation across positions and the standard deviation of conservation scores, generating Z-scores. Z-scores were mapped onto the corresponding PDB structures via the B-factor column for structural visualization. Fully conserved residues were indexed at 1.482 for H_ECT_ and 0.829 for F_ECT_, owing to differences in the number of sequences available for each protein.

### Statistics

All statistical analyses were performed using Graphpad Prism 10.

#### Sequence analyses

CDRH3 lengths, VH residue mutation counts, affinities, and % MeV infection were tested for normality. Correlations were examined by determining the Spearman nonparametric correlation coefficients (r) and P-values for each comparison.

#### Prophylactic protection

We analyzed both raw and normalized titers using the Brown-Forsythe and Welch ANOVA test assuming non-equal standard deviations. We compared each treatment group to the PBS control, and used the Dunnett T3 multiple comparisons correction. Because all data points for 4F09 fell below detection threshold, we used the Mann-Whitney test to compare raw and normalized titers to controls.

