## Supplemental Figures 1-7 for "Uncovering the features of Measles-targeting human antibodies elicited by the MMR vaccine"

### Figure S1.

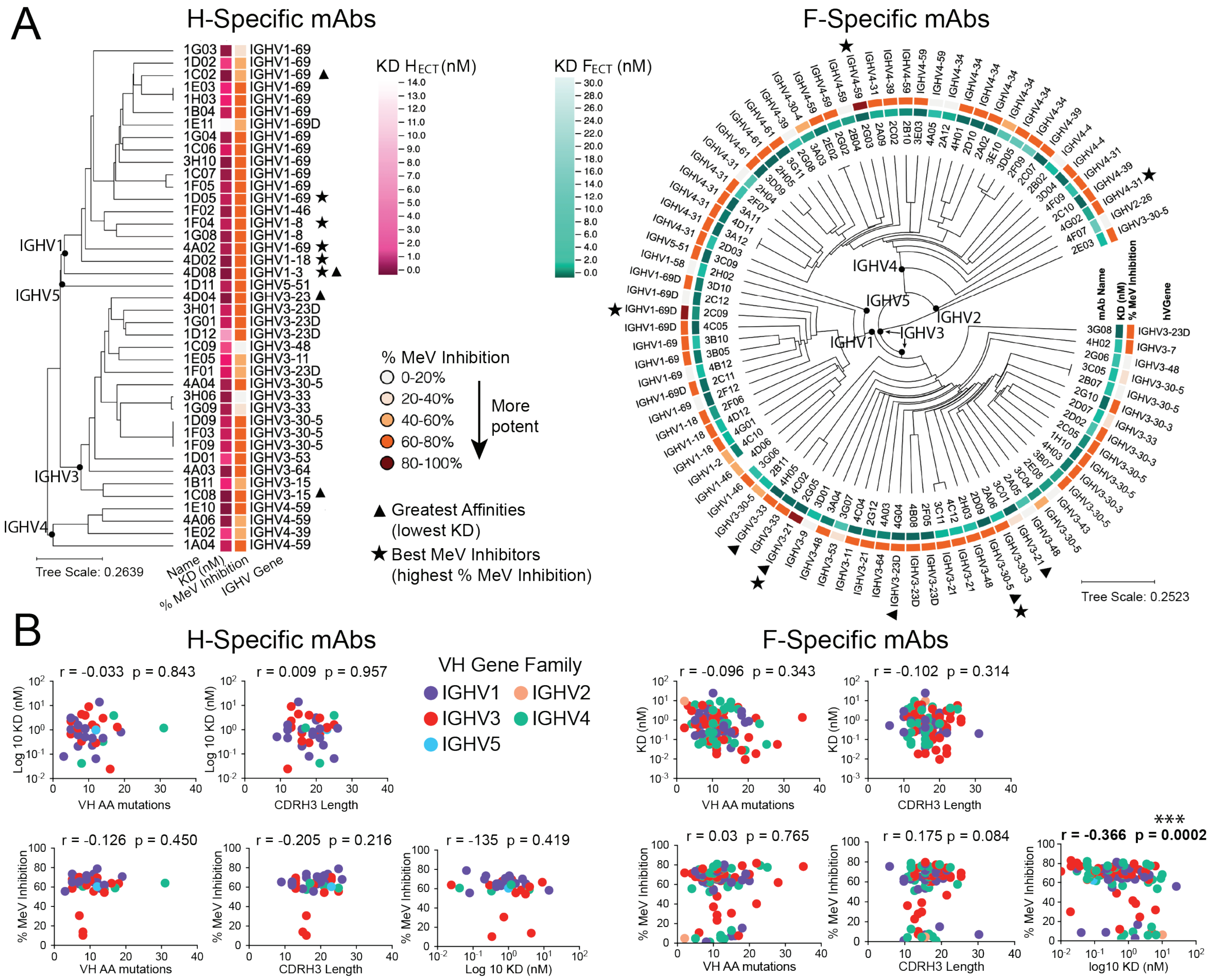
A
H-Specific mAbs
F-Specific mAbs
KD HECT (nM)
KD FECT (nM)
% MeV Inhibition
0-20%
20-40%
40-60%
60-80%
80-100%
More potent
Greatest Affinities (lowest KD)
Best MeV Inhibitors (highest % MeV Inhibition)
Name
KD (nM)
% MeV Inhibition
IGHV Gene
Tree Scale: 0.2639
Tree Scale: 0.2523
IGHV1
IGHV2
IGHV3
IGHV4
IGHV5
mAb Name
hVGene
B
H-Specific mAbs
F-Specific mAbs
VH Gene Family
IGHV1
IGHV2
IGHV3
IGHV4
IGHV5
r = -0.033 p = 0.843
r = 0.009 p = 0.957
r = -0.096 p = 0.343
r = -0.102 p = 0.314
r = -0.126 p = 0.450
r = -0.205 p = 0.216
r = -135 p = 0.419
r = 0.03 p = 0.765
r = 0.175 p = 0.084
***
r = -0.366 p = 0.0002
Log 10 KD (nM)
KD (nM)
% MeV Inhibition
VH AA mutations
CDRH3 Length
Log 10 KD (nM)
log10 KD (nM)

### Figure S2.

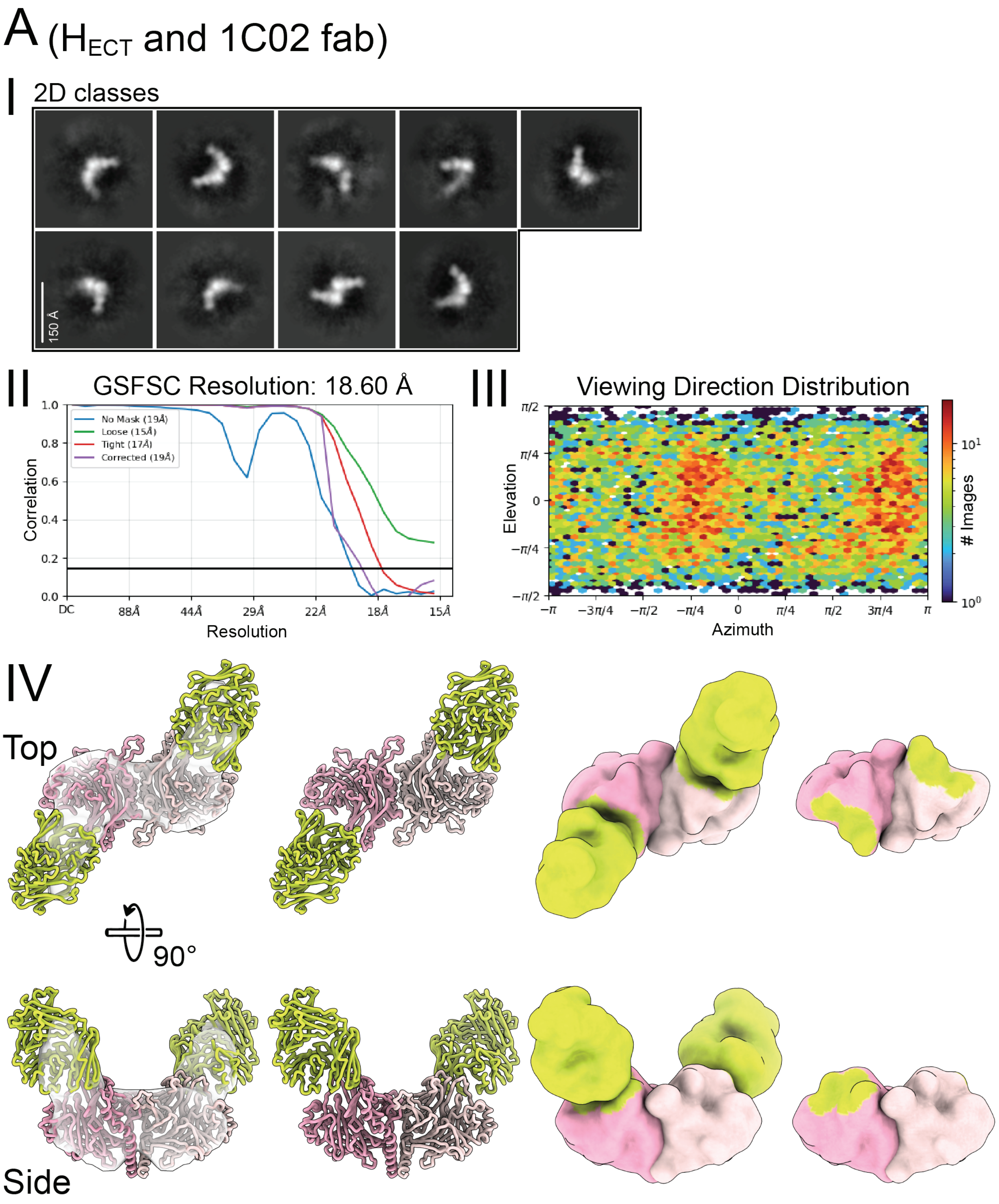

A (H$_{ECT}$ and 1C02 fab)
I
2D classes
150 Å
II
GSFSC Resolution: 18.60 Å
No Mask (19Å)
Loose (15Å)
Tight (17Å)
Corrected (19Å)
Correlation
Resolution
DC
88Å
44Å
29Å
22Å
18Å
15Å
III
Viewing Direction Distribution
Elevation
Azimuth
### Images
IV
Top
90°
Side

### Figure S2.

#### B (H$_{ECT}$ and 4D08 fab with 1C08 fab)

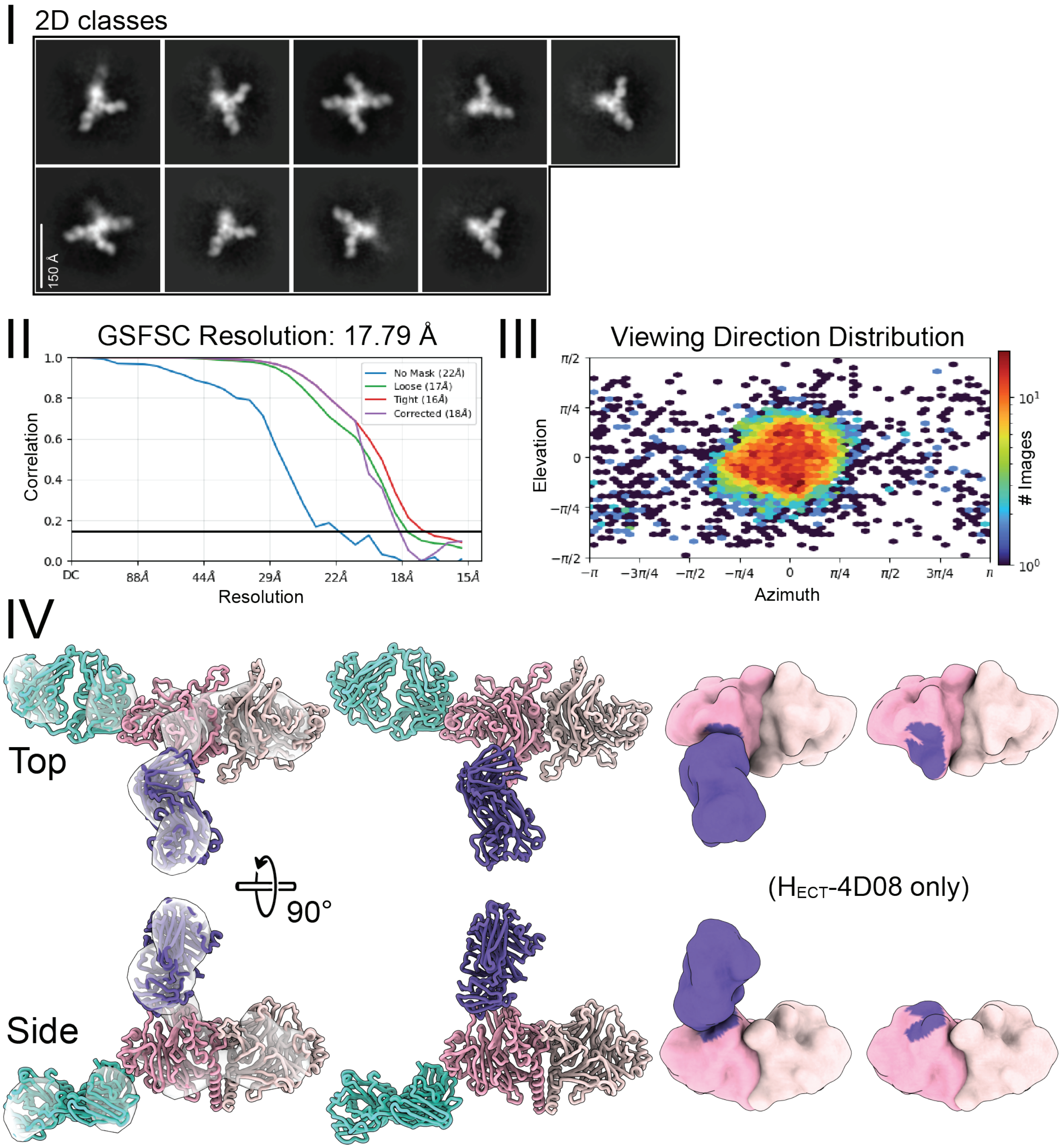

### Figure S2.

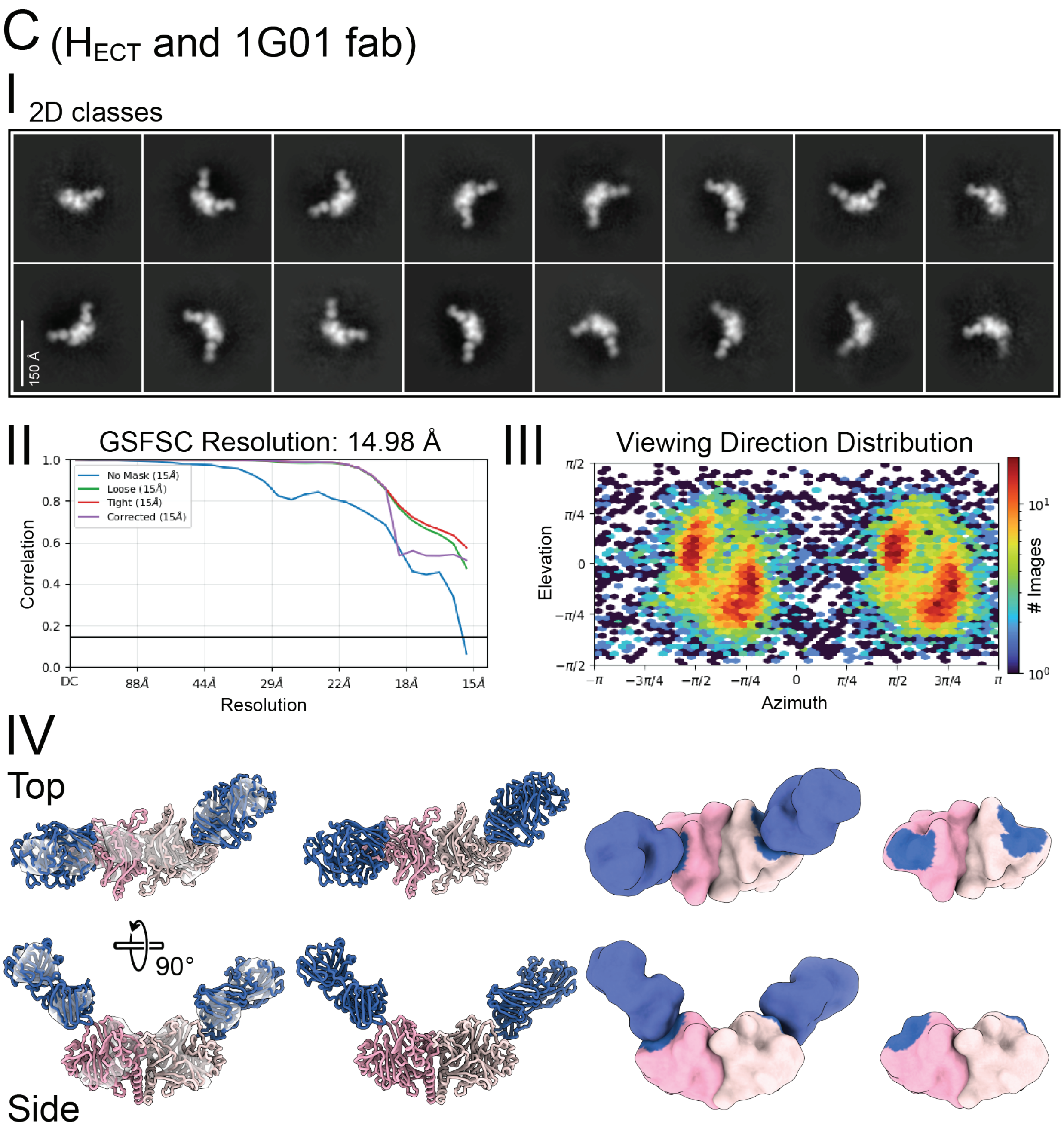

C (HECT and 1G01 fab)
I 2D classes
150 Å
II GSFSC Resolution: 14.98 Å
No Mask (15Å)
Loose (15Å)
Tight (15Å)
Corrected (15Å)
Correlation
Resolution
DC 88Å 44Å 29Å 22Å 18Å 15Å
III Viewing Direction Distribution
Elevation
Azimuth
### Images
IV
Top
90°
Side

### Figure S2.

#### D (H$_{ECT}$ and 4D04 IgG)

I 2D classes

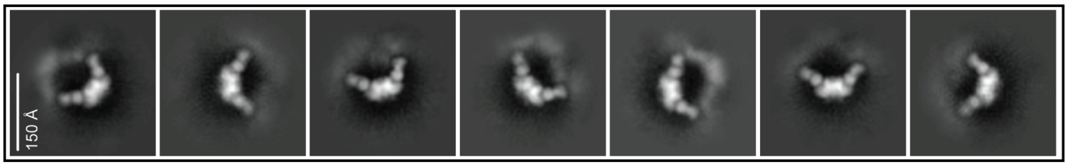


II

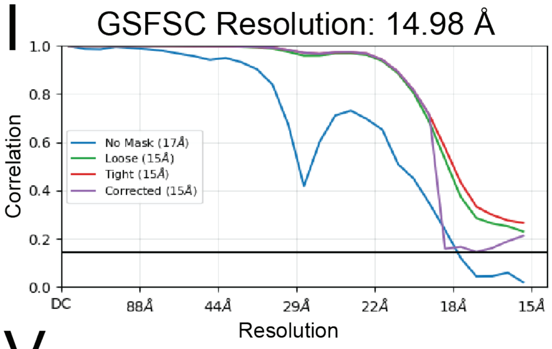


III

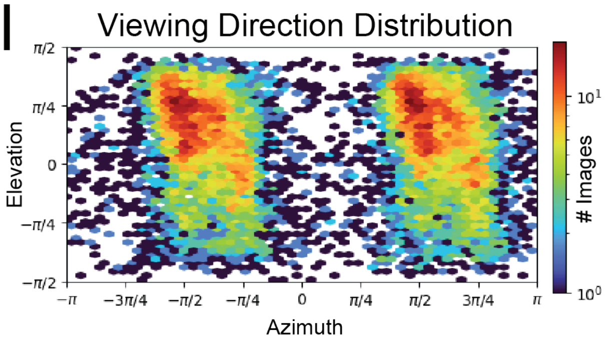


IV

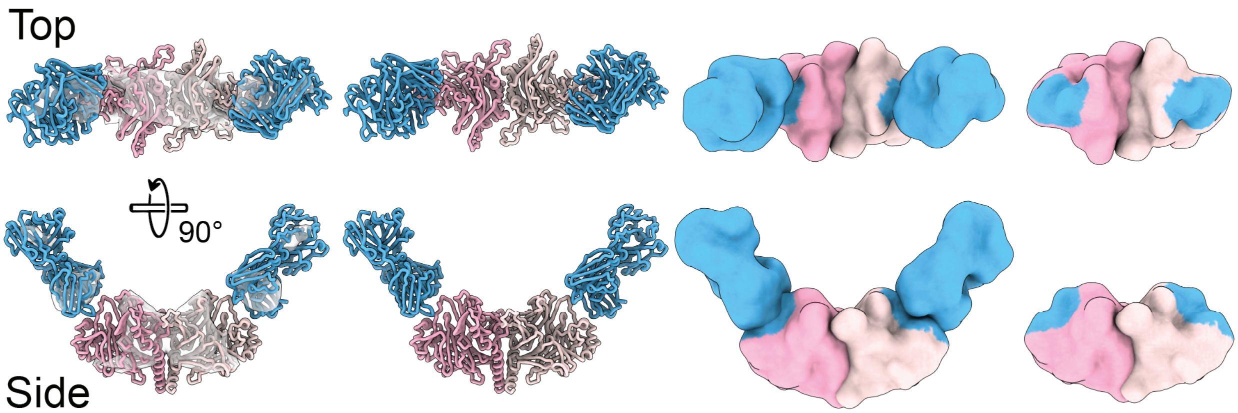

### Figure S2.

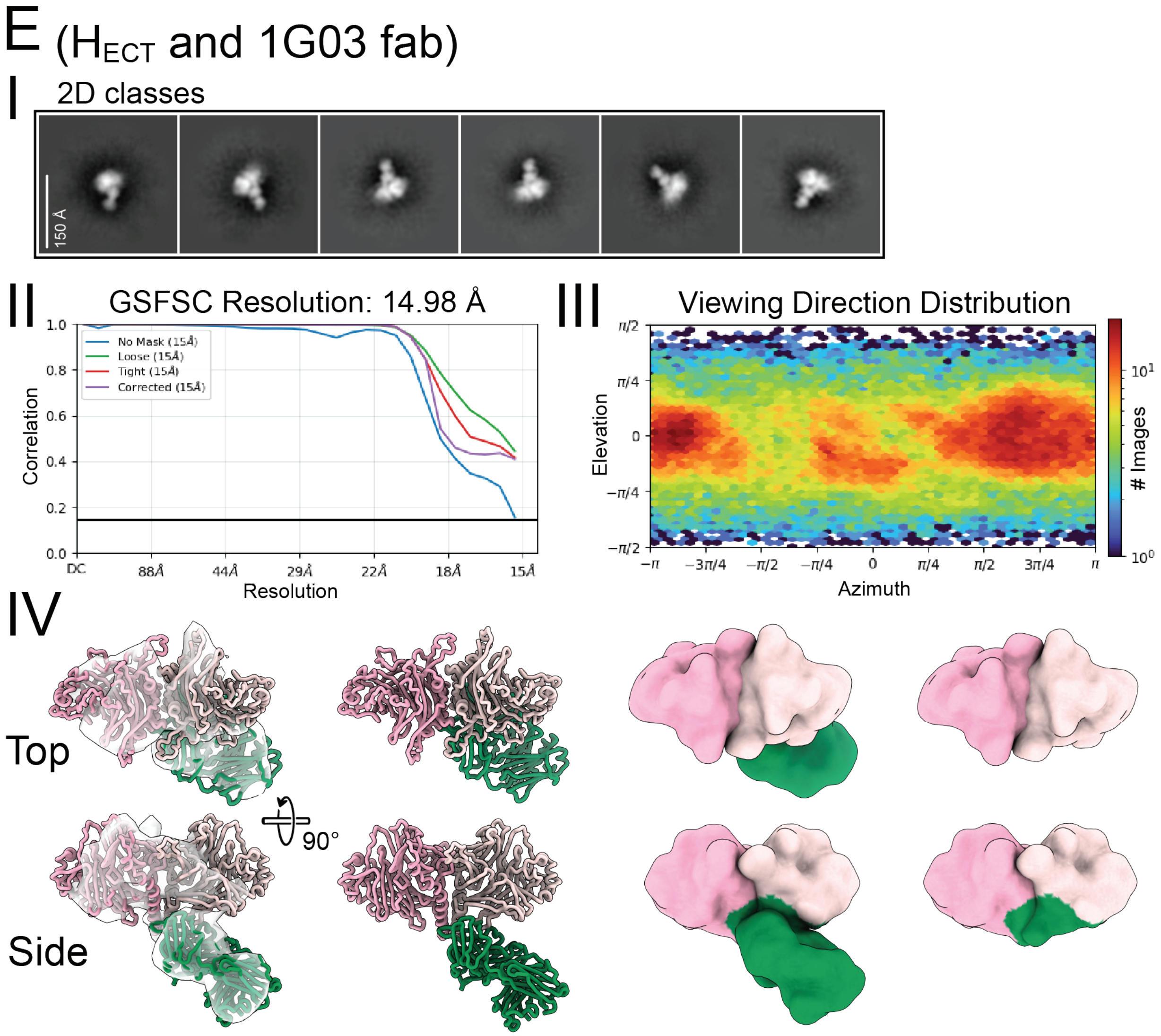
E (H$_{ECT}$ and 1G03 fab)
I
2D classes
150 Å
II
GSFSC Resolution: 14.98 Å
No Mask (15Å)
Loose (15Å)
Tight (15Å)
Corrected (15Å)
Correlation
Resolution
DC
88Å
44Å
29Å
22Å
18Å
15Å
III
Viewing Direction Distribution
Elevation
Azimuth
### Images
IV
Top
Side
90°

### Figure S2.

#### F ($H_{ECT}$ and 1C08 fab)

##### I 2D classes

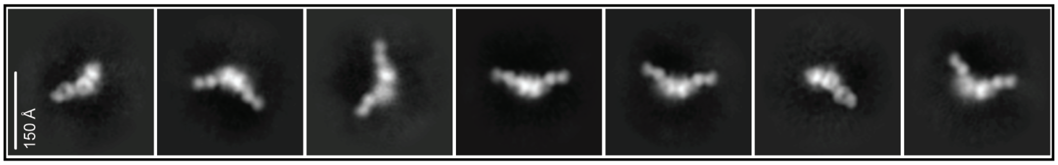


### II

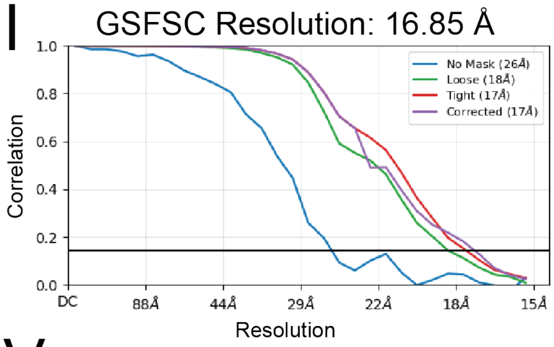


##### III

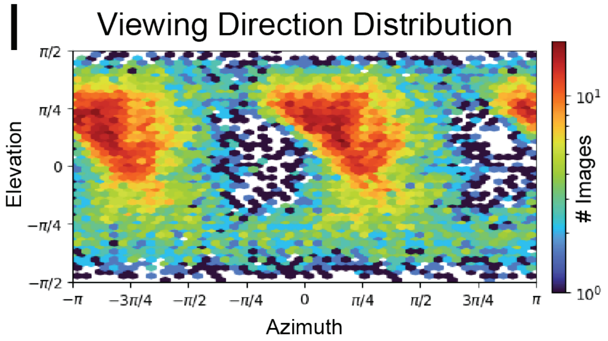


### IV

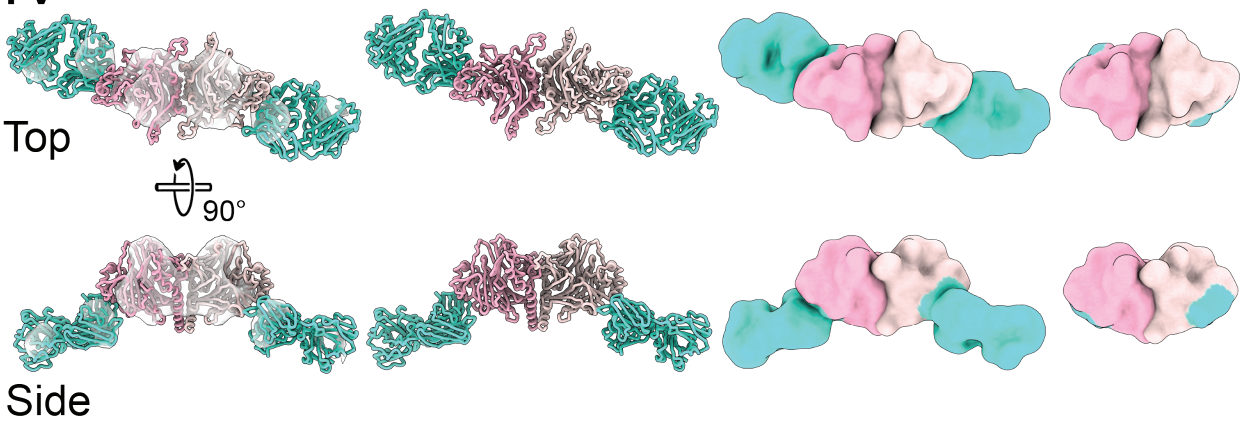

### Figure S2.

### G (F$_{ECT}$ and 2D07 fab)

I 2D classes

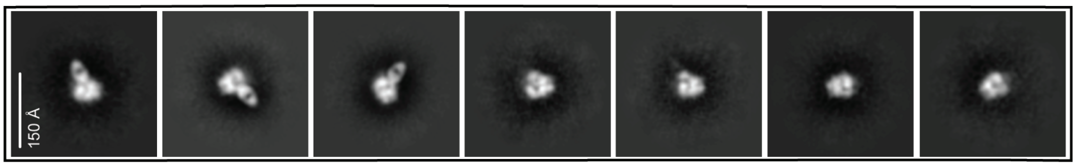


II

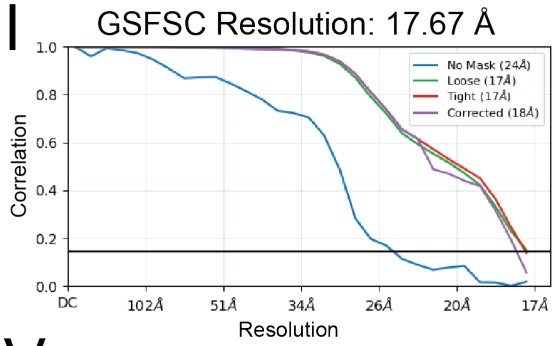


III

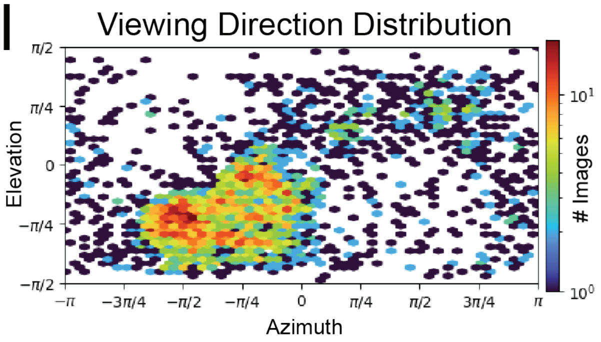


IV

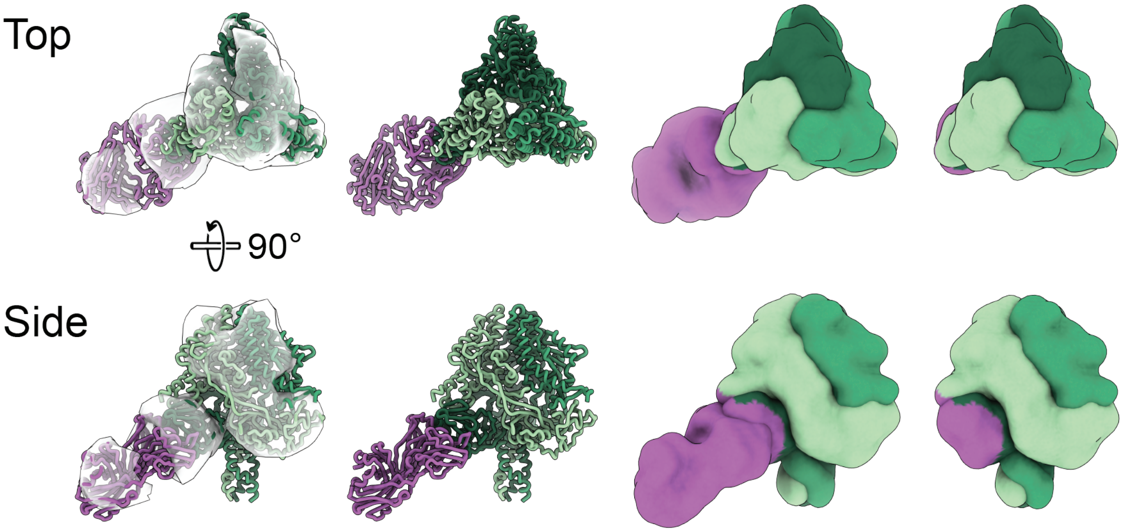

### Figure S2.

#### H (F$_{ECT}$ and 2G05 fab)

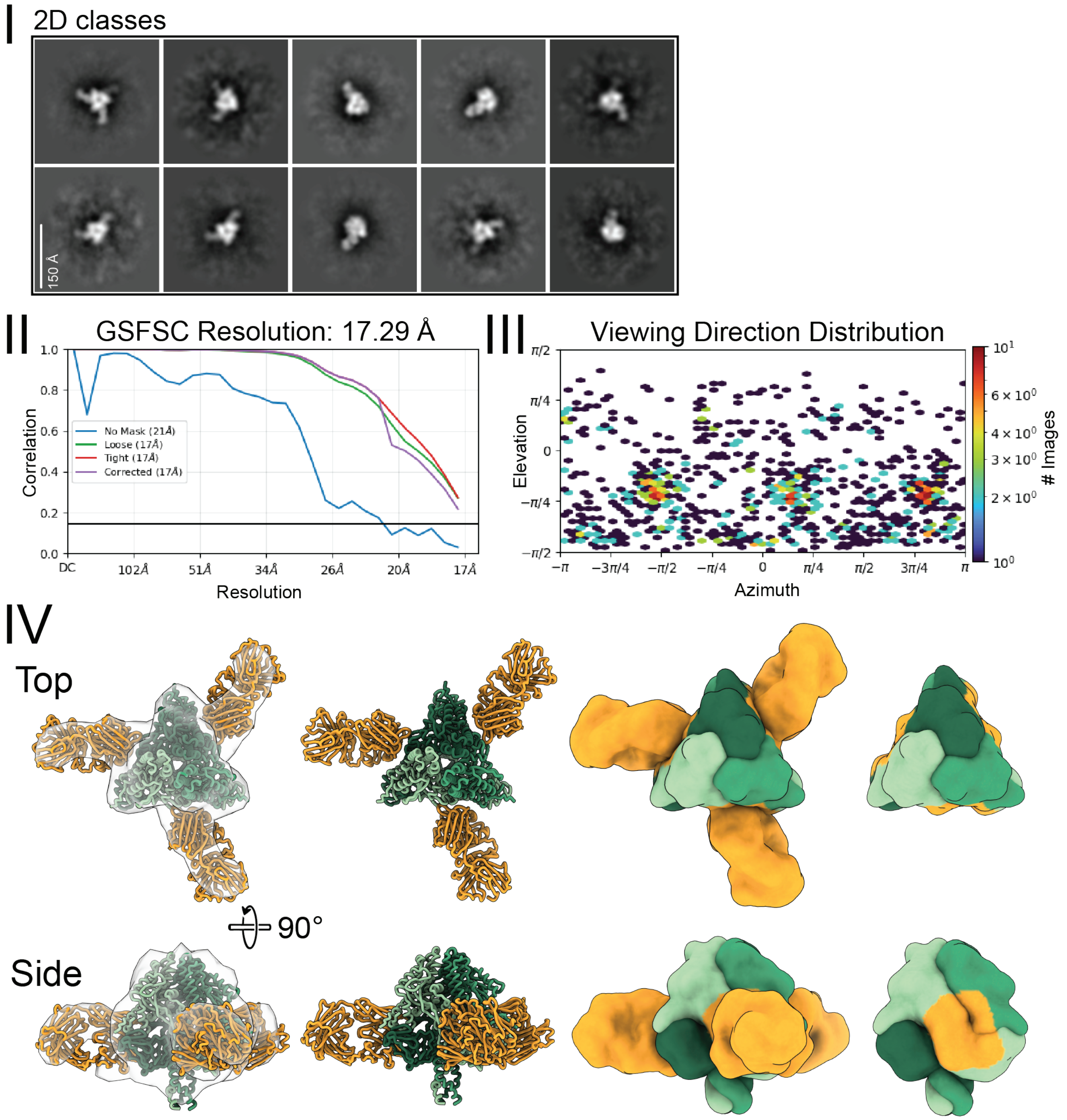

### Figure S2.

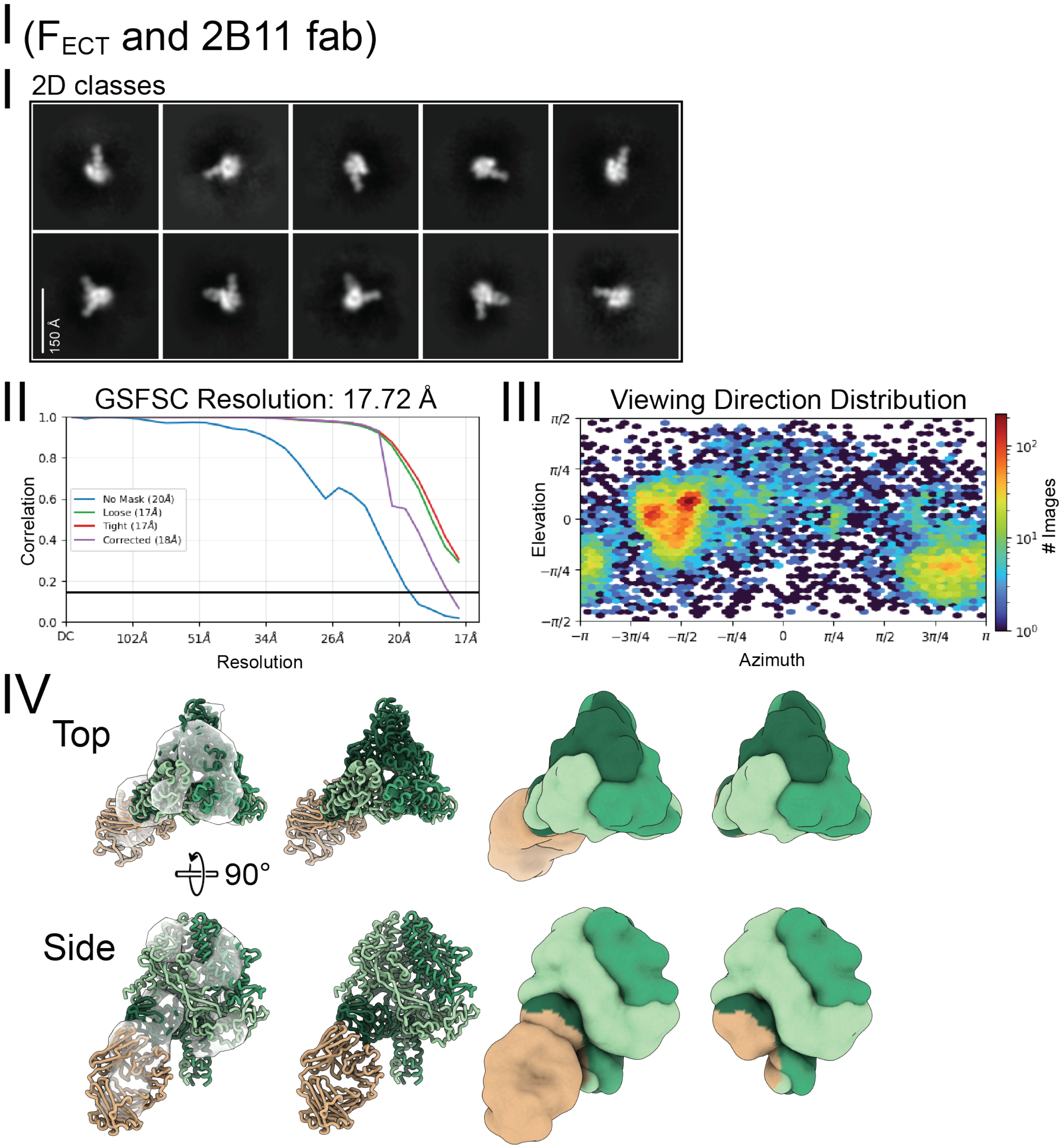

I (FECT and 2B11 fab)
2D classes
150 Å
II
GSFSC Resolution: 17.72 Å
No Mask (20Å)
Loose (17Å)
Tight (17Å)
Corrected (18Å)
Correlation
Resolution
DC
102Å
51Å
34Å
26Å
20Å
17Å
III
Viewing Direction Distribution
Elevation
Azimuth
### Images
IV
Top
90°
Side

### Figure S2.

#### J ($F_{ECT}$ and 3D04 fab)

I 2D classes

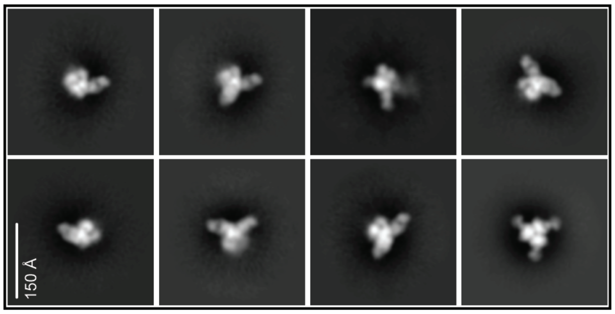


II

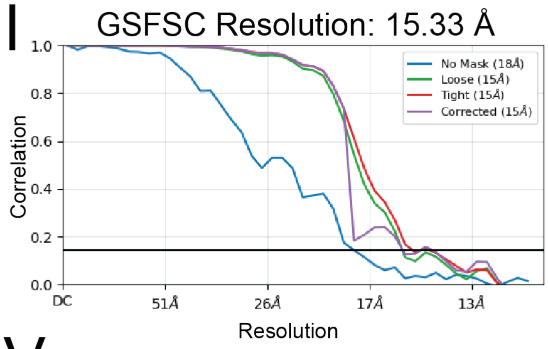


III

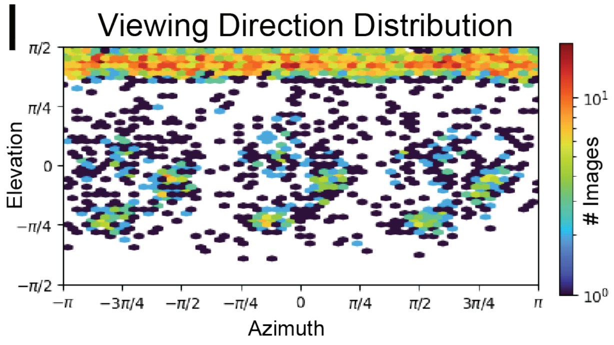


IV

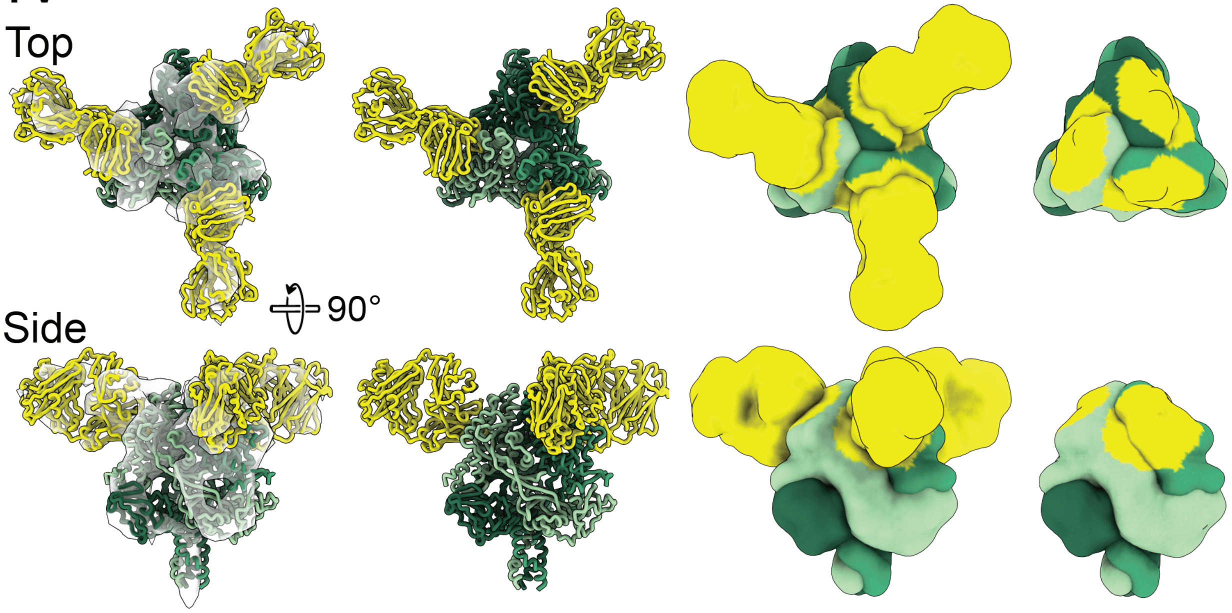

### Figure S2.

#### K (F$_{ECT}$ and 3A12 fab)

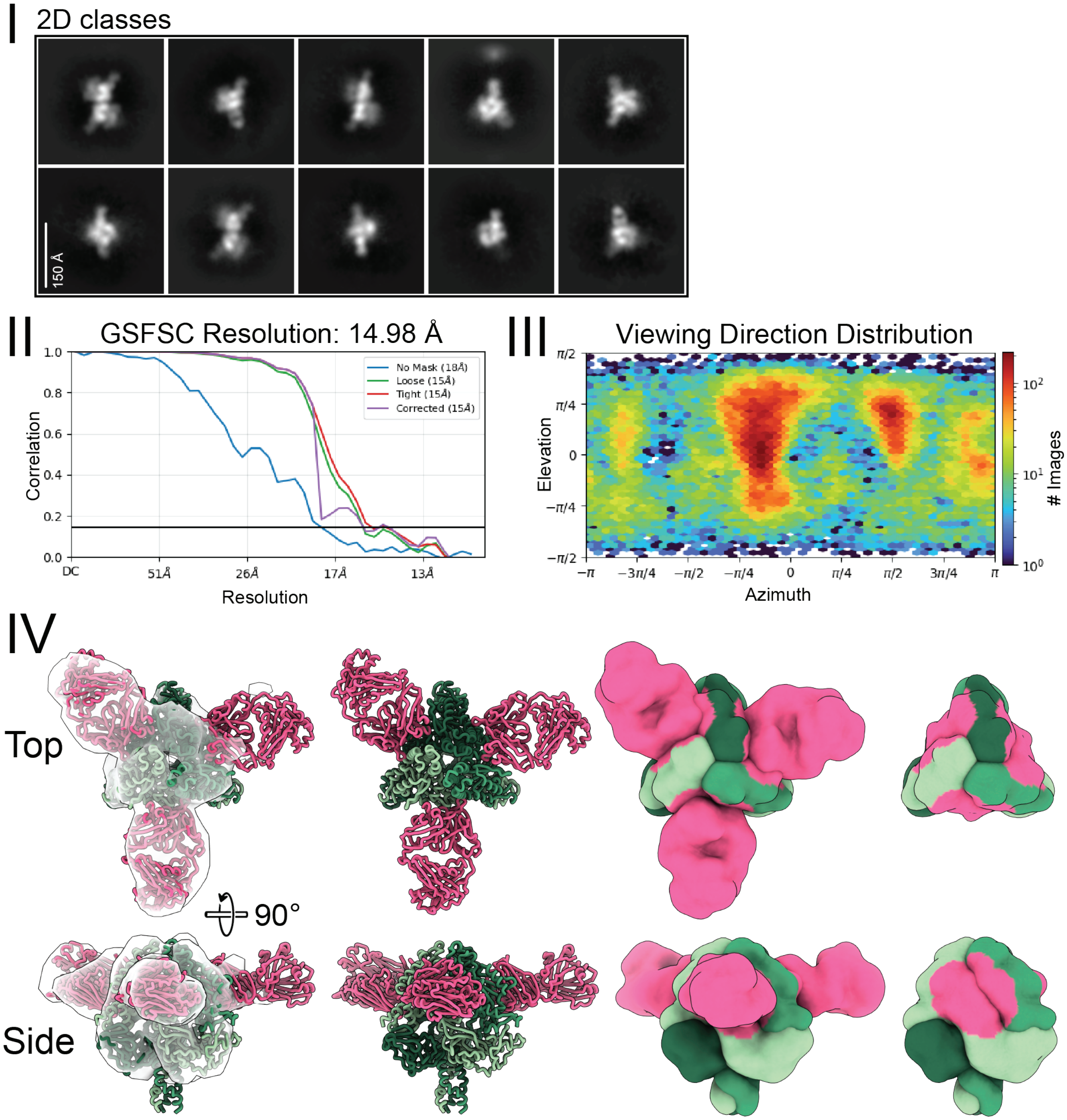

### Figure S2.

#### L ($F_{ECT}$ and 4F09 fab)

##### I 2D classes

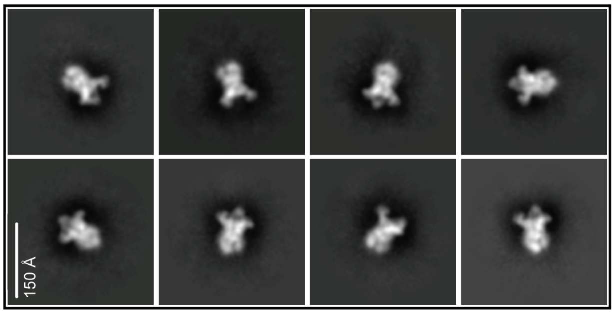


### II

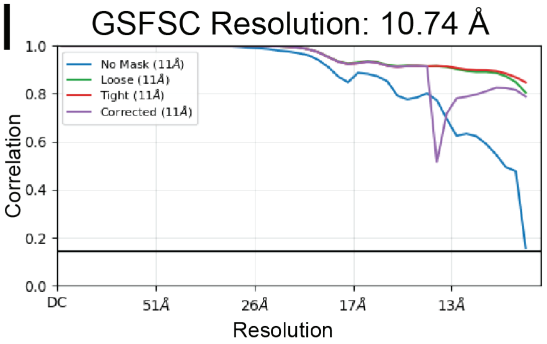


##### III

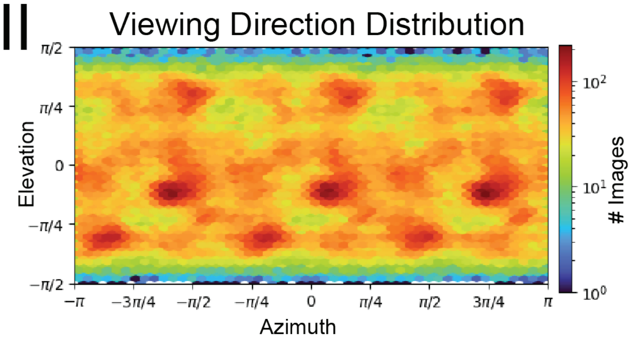


### IV

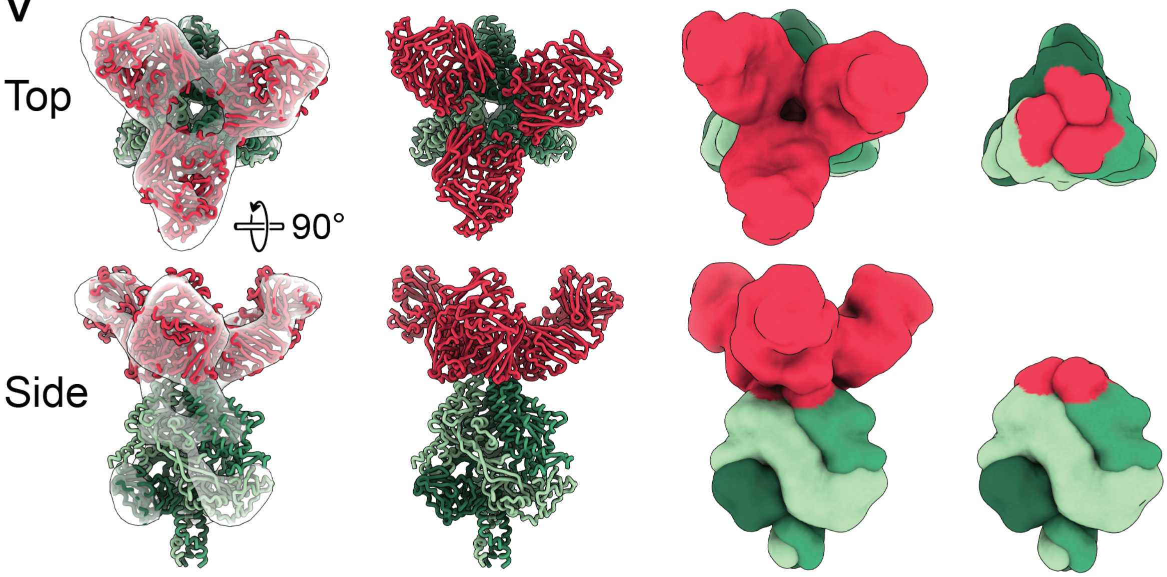

### Figure S3.

A

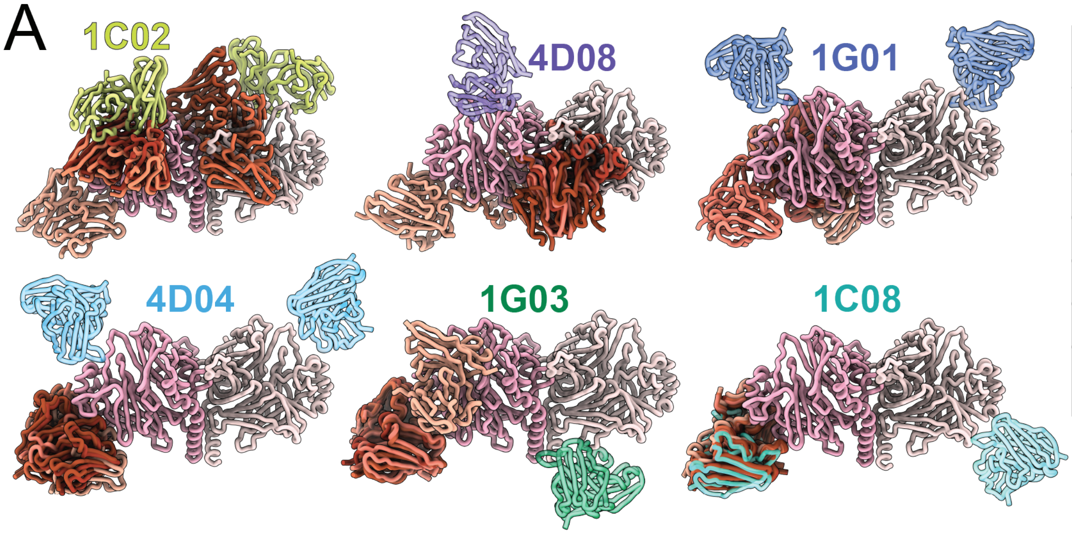


| mAb Name | Accuracy RMSD (Å) | Precision RMSD (Å) |
|---|---|---|
| 1C02 | 53.27 | 37.37 |
| 4D08 | 76.99 | 30.92 |
| 1G01 | 68.86 | 19.55 |
| 4D04 | 57.79 | 8.43 |
| 1G03 | 82.18 | 20.99 |
| **1C08** | **8.67** | **2.09** |
| AVG ± STD | 57.96 ± 26.53 | 19.89 ± 13.24 |

B

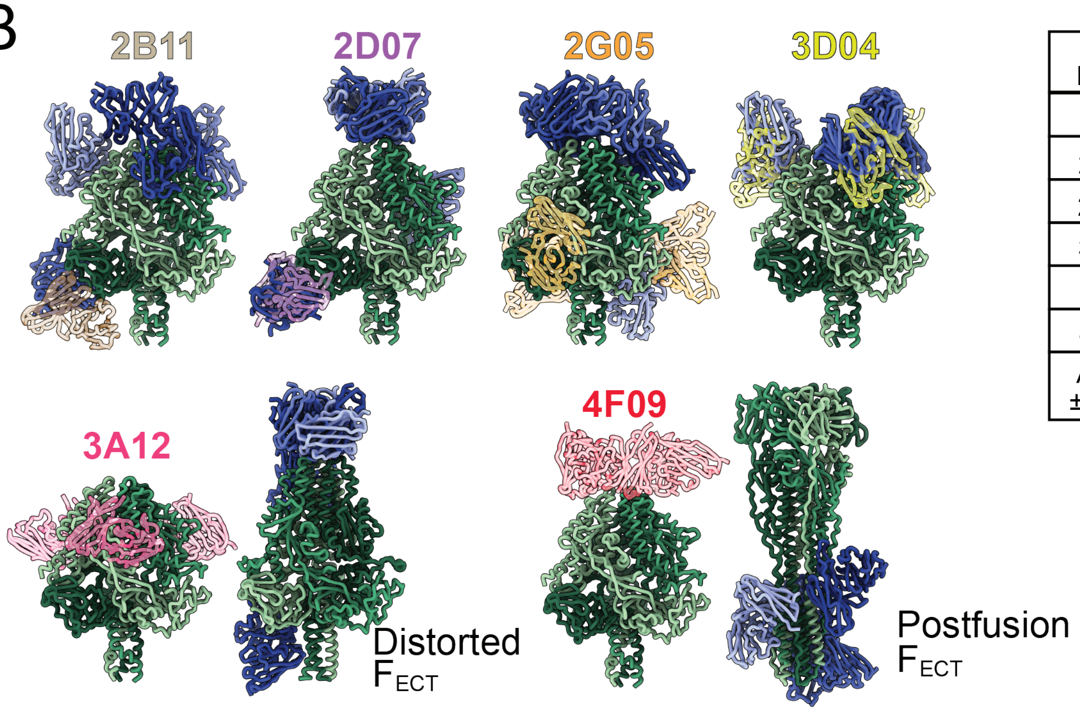


| mAb Name | Accuracy RMSD (Å) | Precision RMSD (Å) |
|---|---|---|
| 2B11 | 105.50 | 42.11 |
| 2D07 | 96.64 | 49.55 |
| 2G05 | 87.55 | 46.52 |
| **3D04** | **12.24** | **1.55** |
| 3A12 | 66.58 | 50.46 |
| 4F09 | 94.64 | 28.09 |
| AVG ± STD | 77.22 ± 34.43 | 36.39 ± 18.92 |

### Figure S4.

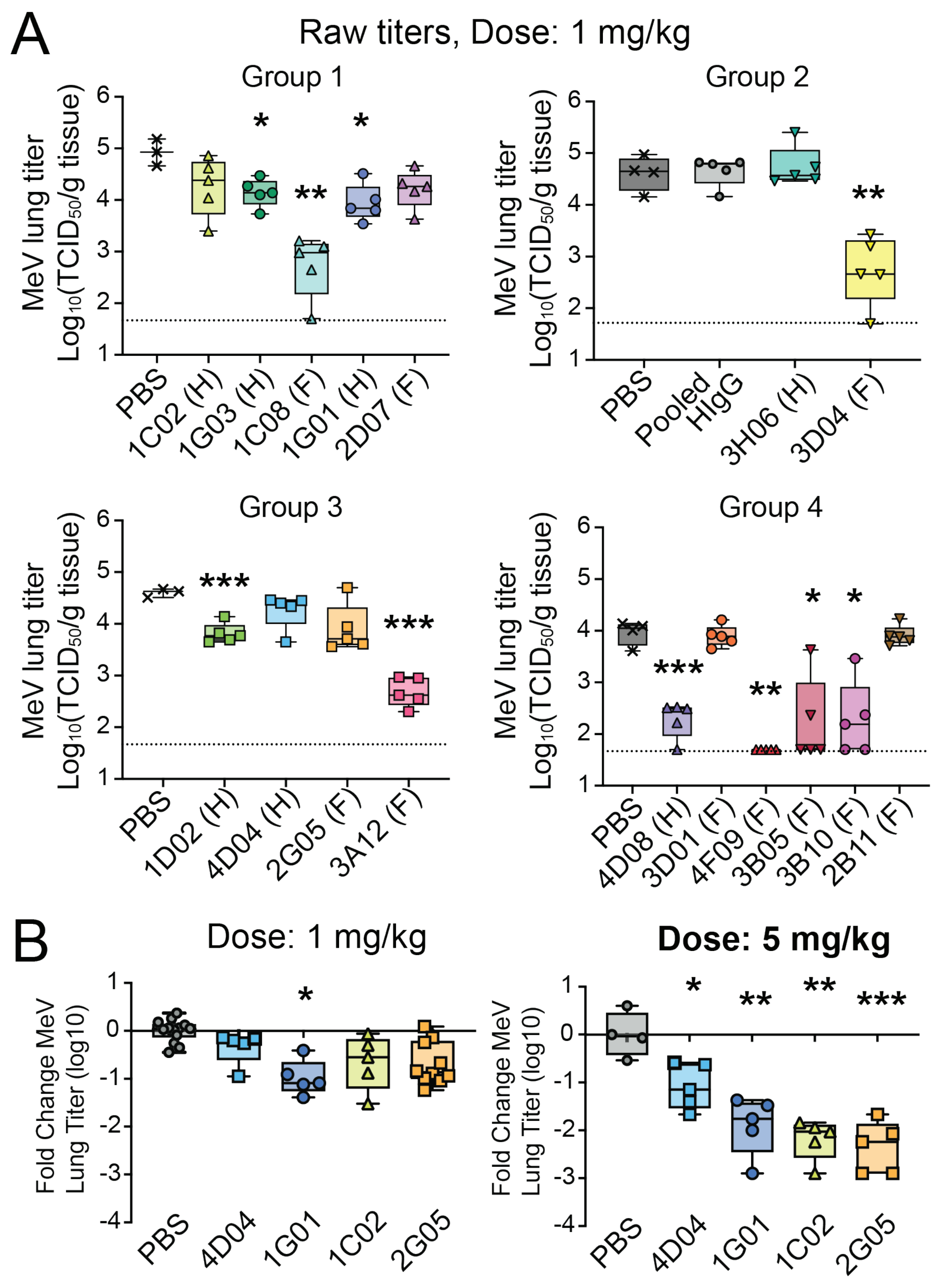
A
Raw titers, Dose: 1 mg/kg
Group 1
Group 2
Group 3
Group 4
MeV lung titer
Log10(TCID50/g tissue)
PBS
1C02 (H)
1G03 (H)
1C08 (F)
1G01 (H)
2D07 (F)
Pooled HIgG
3H06 (H)
3D04 (F)
1D02 (H)
4D04 (H)
2G05 (F)
3A12 (F)
4D08 (H)
3D01 (F)
4F09 (F)
3B05 (F)
3B10 (F)
2B11 (F)
B
Dose: 1 mg/kg
Dose: 5 mg/kg
Fold Change MeV
Lung Titer (log10)
PBS
4D04
1G01
1C02
2G05

### Figure S5.

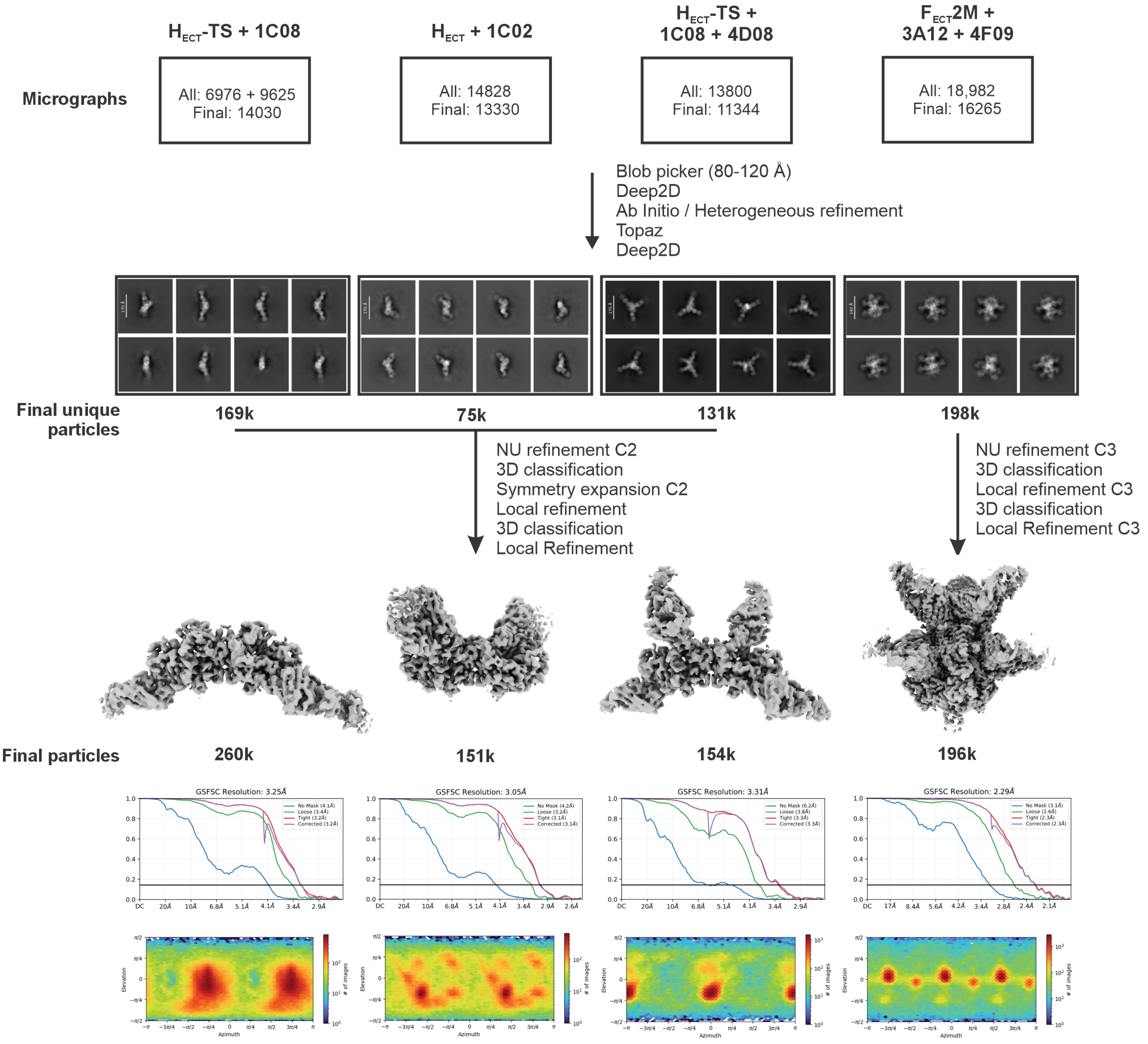

$H_{ECT}$-TS + 1C08
$H_{ECT}$ + 1C02
$H_{ECT}$-TS + 1C08 + 4D08
$F_{ECT}$2M + 3A12 + 4F09
Micrographs
All: 6976 + 9625
Final: 14030
All: 14828
Final: 13330
All: 13800
Final: 11344
All: 18,982
Final: 16265
Blob picker (80-120 Å)
Deep2D
Ab Initio / Heterogeneous refinement
Topaz
Deep2D
Final unique particles
169k
75k
131k
198k
NU refinement C2
3D classification
Symmetry expansion C2
Local refinement
3D classification
Local Refinement
NU refinement C3
3D classification
Local refinement C3
3D classification
Local Refinement C3
Final particles
260k
151k
154k
196k
GSFSC Resolution: 3.25Å
GSFSC Resolution: 3.05Å
GSFSC Resolution: 3.31Å
GSFSC Resolution: 2.29Å
Elevation
Azimuth
### of images

### Figure S6.

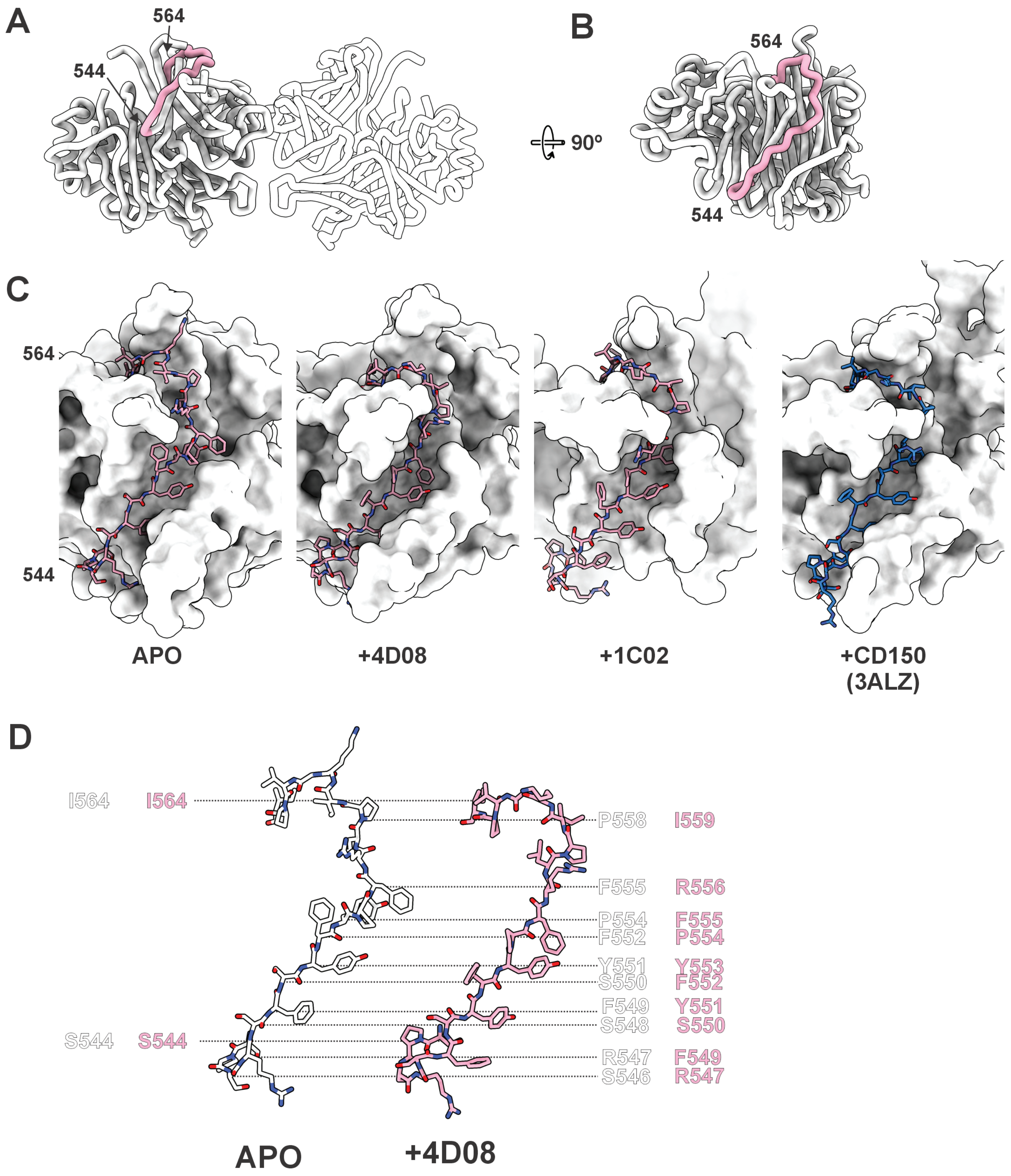
A
564
544
B
564
544
90º
C
564
544
APO
+4D08
+1C02
+CD150
(3ALZ)
D
I564
I564
P558
I559
F555
R556
P554
F555
F552
P554
Y551
Y553
S550
F552
F549
Y551
S548
S550
S544
S544
R547
F549
S546
R547
APO
+4D08

### Figure S7.

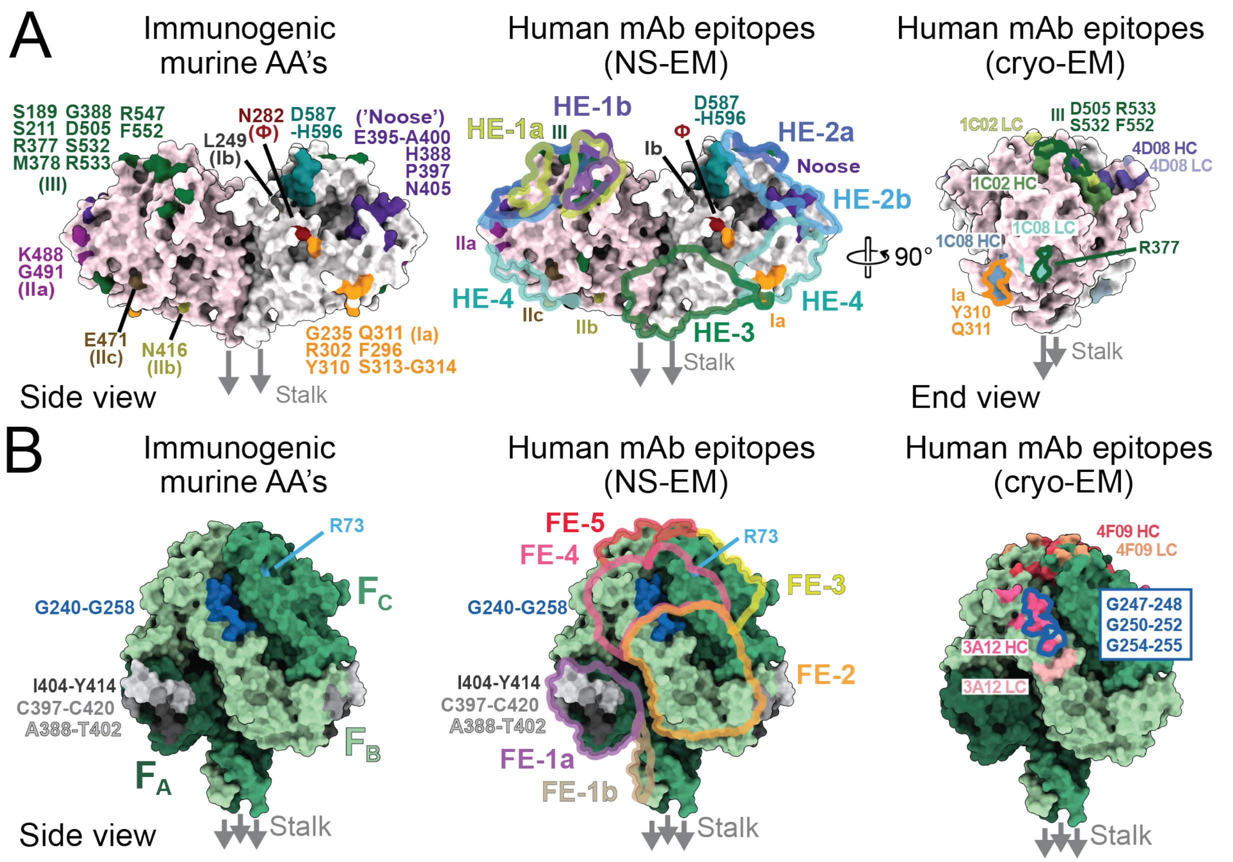


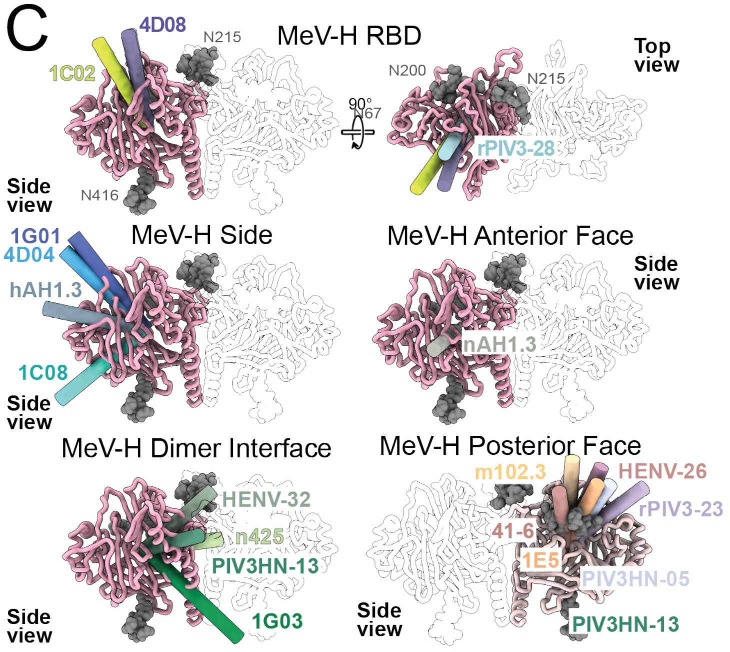


| mAb Name | Species | Antigen | Epitope on MeV-H | PDB ID |
|---|---|---|---|---|
| 1C02 | Human | MeV-H | RBD (HE-1a) | This study |
| 4D08 | Human | MeV-H | RBD (HE-1b) | This study |
| rPIV3-28 | Human | PIV3-HN | RBD | 8TQI |
| 1G01 | Human | MeV-H | Side (upper, HE-2a) | This study |
| 4D04 | Human | MeV-H | Side (upper, HE-2B) | This study |
| hAH1.3 | Mouse | HeV-G | Side (middle) | 7SYY |
| 1C08 | Human | MeV-H | Side (lower, HE-4) | This study |
| HENV-32 | Human | HeV-G | Dimer interface | 6VY4 |
| n425 | Human | NiV-G | Dimer interface | 8XPS |
| PIV3HN-13 | Human | PIV3-HN | Dimer interface | 9DZQ |
| 1G03 | Human | MeV-H | Dimer interface (HE-3) | This study |
| nAh1.3 | Mouse | NiV-G | Anterior face | 7TXZ |
| m102.3 | Human | HeV-G | Posterior face | 6CMI |
| 1E5 | NHP | NiV-G | Posterior face | 8XC4 |
| 41-6 | Human | NiV-G | Posterior face | 8K3C |
| HENV-26 | Human | HeV-G | Posterior face | 6VY6 |
| rPIV3-23 | Human | PIV3-HN | Posterior face | 8TQi |
| PIV3HN-05 | Human | PIV3-HN | Posterior face | 9DZQ |
