## Supplementary material for "Uncovering the features of Measles-targeting human antibodies elicited by the MMR vaccine": Table S1

| Name | Epitope | VH[a] Gene | VL[a] Gene | Ka[b] (1/Ms) | Kd[b] (1/Ms) | KD[b] (nM) | EC50[c] (ug/mL) | Protective? | CDRH-3 AA Sequence[d] |
|---|---|---|---|---|---|---|---|---|---|
| 1D02 | HE-1a | IGHV1-69 | IGKV2D-28 | 3.1E+05 | 2.1E-04 | 0.65 | 0.176 | No | AREMATISLLYYYGLDV |
| 1C02 | HE-1a | IGHV1-69 | IGLV1-40 | 1.9E+05 | 2.7E-05 | 0.18 | 0.019 | No | ARERITVAGINYYYGMDV |
| 4D08 | HE-1b | IGHV1-2 | IGKV1D-39 | 1.6E+05 | 3.0E-05 | 0.19 | 0.025 | Yes | ARDAYDFWSGYGNWGSNYYYFYMDA |
| mAb55 | HE-1b | IGHV1-66 (mouse) | IGKV5-37 (mouse) | 1.0E+05 | 1.4E-04 | 1.20 | 0.069 | No | ASGTDRYFDY |
| 1G01 | HE-2a | IGHV3-23D | IGLV1-51 | 5.5E+04 | 1.0E-05 | 0.18 | 0.113 | No | ARDGYSSYWYLDN |
| 4D04 | HE-2b | IGHV3-23 | IGLV1-51 | 1.1E+05 | 2.7E-05 | 0.27 | 0.121 | No | ARALYSSRGSTSFVQ |
| 1G03 | HE-3 | IGHV1-69 | IGLV1-51 | 5.5E+04 | 2.8E-05 | 0.53 | ND | No | ATAGIGVAAKRKPSVRTFDL |
| 3H06 | HE-4 | IGHV3-33 | IGLV2-14 | 1.0E+05 | 3.0E-05 | 0.29 | ND | No | ARDRAPYNSYYYYMDV |
| 1C08 | HE-4 | IGHV3-15 | IGKV2D-28 | 2.5E+05 | 2.9E-05 | 0.12 | 0.013 | Yes | TTDLFRFGFGDV |
| 2D07 | FE-1a | IGHV3-33 | IGKV3-11 | 1.3E+05 | 4.8E-04 | 3.66 | 3.347 | No | ARDGVAYGLDV |
| 2B11 | FE-1b | IGHV3-30-5 | IGLV3-21 | 6.6E+04 | 8.3E-05 | 1.26 | 64.84 | No | AKDFRAYSYGDTFDY |
| 2G05 | FE-2 | IGHV3-21 | IGKV3-15 | 1.8E+05 | 4.5E-05 | 0.24 | 0.237 | No | ARDQLEGESYFYLMDA |
| mAb77 | FE-2 | IGHV4-30-4 (mouse) | IGKV16-104 (mouse) | 1.6E+05 | 2.5E-05 | 0.16 | 0.041 | Yes | ARSGWLLPYWYFDV |
| 3D01 | FE-2 | IGHV3-9 | IGLV2-23 | 1.9E+05 | 5.1E-05 | 0.27 | ND | No | AKGVQWLVRGYFDY |
| 3D04 | FE-3 | IGHV4-4 | IGKV3-15 | 1.0E+05 | 2.2E-05 | 0.22 | 0.067 | Yes | ARFTADLCFDY |
| 3B10 | FE-4 | IGHV1-69 | IGKV1D-39 | 2.6E+04 | 2.5E-04 | 9.85 | 0.029 | Yes | AKEVHGYMDV |

| | | | | | | | | | |
|---|---|---|---|---|---|---|---|---|---|
| 3A12 | FE-4 | IGHV4-31 | IGKV1D-39 | 2.9E+04 | 9.6E-05 | 3.29 | 0.054 | Yes | ARATSKDASGRHLLDY |
| 3B05 | FE-5 | IGHV1-69 | IGKV1D-39 | 1.7E+04 | 9.1E-05 | 5.26 | 0.031 | Yes | ARQAHGYMDV |
| 4F09 | FE-5 | IGHV4-31 | IGKV1-5 | 9.0E+04 | 3.0E-05 | 0.34 | 0.118 | Yes | ARGIQGPYYYDRSGYYWPPYFDY |

**Table S1. Properties of MeV mAbs used to define human epitopes on H and F.** [a]VH= heavy chain variable region, VL= light chain variable region. [b]MAb kinetics were determined by measuring mAb binding interactions with MeV $H_{ECT}$ or $F_{ECT}$ via surface plasmon resonance, and data was fit to a 1:1 Languir model to determine the Ka (on-rate), Kd (off-rate), and KD (equilibrium dissociation rate) for each mAb using the Kinetics software package (Carterra). [c]$EC_{50}$= half-maximal effective dose. [d]The amino acid sequence of the complementarity-determining region-3 on the mAb heavy chain.
