## Supplementary material for "Uncovering the features of Measles-targeting human antibodies elicited by the MMR vaccine": Table S2

| **$H_{ECT}$ and 1C08** | | | | | | | | |
|---|---|---|---|---|---|---|---|---|
| H chain & residue | Location | 1C08 residue | Location | Interaction Type | Atom Interaction | % H residue frequency | H residue conservation index | Variants |
| A I308 | β2 | D R103 | CDRH-3 | Hydrogen bond | Backbone - Side chain | 97.37 | -1.537 | V (59) M (1) N (1) T (1) |
| A Y310 | β2 | D D55 | CDRH-2 | Hydrogen bond | Backbone - Side chain | 99.92 | -0.464 | D (1) H (1) |
| A Q311 | β2 | D R51 | CDRH-2 | Hydrogen bond | Backbone - Side chain | 99.45 | -1.085 | H (9) R (2) K (2) |
| A I346 | β3 | D R103 | CDRH-3 | Hydrogen bond | Backbone - Side chain | 84.13 | -0.551 | V (373) M (1) |
| A D347 | β4 | D R30 | CDRH-1 | Salt bridge | Side chain - Side chain | 100 | 1.482 | |
| A D373 | β3 | E Y32 | CDRL-1 | Hydrogen bond | Side chain - Side chain | 99.96 | 0.321 | A (1) |
| A R377 | β3 | E Y32 | CDRL-1 | Hydrogen bond | Side chain - Side chain | 100 | 1.482 | |
| **$H_{ECT}$ and 4D08** | | | | | | | | |
| H chain & residue | Location | 4D08 residue | Location | Interaction Type | Atom Interaction | H residue frequency | H residue conservation index | Variants |
| A G190 | β1 | B N109 | CDRH-3 | Hydrogen bond | Backbone - Side chain | 100 | 1.482 | |
| A T192 | β1 (RBD) | B G108 | CDRH-3 | Hydrogen bond | Backbone - Backbone | 99.7 | -1.607 | A (3) I (3) P (1) |
| A I194 | β1 (RBD) | B G106 | CDRH-3 | Hydrogen bond | Backbone - Backbone | 99.92 | -0.463 | I (3) |
| A S532 | β5 (RBD) | B N55 | CDRH-2 | Hydrogen bond | Backbone - Side chain | 98.56 | 0.308 | F (1) |
| A S532 | β5 (RBD) | B N55 | CDRH-2 | Hydrogen bond | Side chain - Side chain | 98.56 | 0.308 | F (1) |
| A R533 | β5 (RBD) | B F103 | CDRH-3 | Hydrogen bond | Side chain - Backbone | 98.6 | 1.482 | |

| | | | | | | | | |
|---|---|---|---|---|---|---|---|---|
| A T607 | β6 | C S52 | CDRL-2 | Hydrogen bond | Backbone - Side chain | 98.13 | -1.543 | P (4) A (3) S (1) I (1) |
| **$H_{ECT}$ and 1C02** | | | | | | | | |
| H chain & residue | Location | 1C02 residue | Location | Interaction Type | Atom Interaction | H residue frequency | H residue conservation index | Variants |
| D V504 | β5 | C R100 | CDRH-3 | Hydrogen bond | Backbone - Side chain | 98.51 | -0.484 | A (1) M (1) |
| D D505 | β5 (RBD) | C N107 | CDRH-3 | Hydrogen bond | Side chain - Side chain | 98.34 | -0.488 | G (4) Y (2) |
| D D505 | β5 (RBD) | C R100 | CDRH-3 | Salt bridge | Side chain - Side chain | 98.34 | -0.488 | G (4) Y (2) |
| D D507 | β5 (RBD) | C N107, C Y110 | CDRH-3, CDRH-3 | Hydrogen bond | Side chain - Side chain | 98.6 | 1.482 | |
| D S532 | β5 (RBD) | C N107 | CDRH-3 | Hydrogen bond | Side chain - Side chain | 98.56 | 0.308 | F (1) |
| D Y541 | β5 (RBD) | C I106 | CDRH-3 | Hydrogen bond | Side chain - Backbone | 98.56 | 0.306 | N (1) |
| D Y543 | β5 (RBD) | C A104 | CDRH-3 | Hydrogen bond | Side chain - Backbone | 98.51 | 0.307 | H (2) |
| D S548 | β5 (RBD) | C Y60 | FRH-3 | Hydrogen bond | Backbone - Backbone | 98.56 | 0.308 | P (1) |
| D S548 | β5 (RBD) | C R65 | CDRH-2 | Hydrogen bond | Backbone - Side chain | 98.56 | 0.308 | P (1) |
| D F549 | β5 (RBD) | C A58 | CDRH-2 | Hydrogen bond | Backbone - Backbone | 98.47 | 0.307 | L (3) |
| D G190 | β1 | E S116 | CDRL-3 | Hydrogen bond | Backbone - Side chain | 100 | 1.482 | |
| D R533 | β5 (RBD) | E D55 | CDRL-1 | Salt bridge | Side chain - Side chain | 98.6 | 1.482 | |
| D S550 | β5 (RBD) | E S119, E Y114 | CDRL-3, CDRL-3 | Hydrogen bond | Backbone - Side chain | 98.26 | -0.488 | F (5) A (3) |
| D F552 | β5 (RBD) | E Y114, E Y54 | CDRH-3, CDRL-1 | Hydrogen bond | Backbone - Side chain | 98.39 | -0.494 | S (4) L (1) |

| | | | | | | | | |
|---|---|---|---|---|---|---|---|---|
| D Y553 | β5 (RBD) | E A52 | CDRL-1 | Hydrogen bond | Backbone - Side chain | 98.47 | 0.305 | F (3) |
| D R556 | β5 (RBD) | E G53 | CDRL-1 | Hydrogen bond | Side chain - Backbone | 98.6 | 1.482 | |
| D R556 | β5 (RBD) | E D55 | CDRL-1 | Salt bridge | Side chain - Side chain | 98.6 | 1.482 | |
| **F<sub>ECT</sub> and 3A12** | | | | | | | | |
| F chain & residue | Location | 3A12 residue | Location | Interaction Type | Atom Interaction | H residue frequency | H residue conservation index | Variants |
| B Q52 | F2 | D S59 | CDRH-2 | Hydrogen bond | Side chain - Side chain | 99.65 | -1.361 | H (4) P(1) |
| B R165 | F1 HRN | D Y56 | CDRH-2 | Hydrogen bond | Side chain - Backbone | 99.44 | -1.963 | G (4) K (3) I (1) |
| B E247 | F1 D1 | D Y55, D Y56 | CDRH-2, CDRH-2 | Hydrogen bond | Side chain - Side chain | 99.79 | -1.362 | G (2) K (1) |
| B K248 | F1 D1 | D S57, D S59 | CDRH-2, CDRH-2 | Hydrogen bond | Backbone - Side chain | 100 | 0.829 | |
| B G250 | F1 D1 | D Y53 | CDRH-2 | Hydrogen bond | Backbone - Side chain | 100 | 0.829 | |
| B Y251 | F1 D1 | D Y36 | CDRH-1 | Hydrogen bond | Backbone - Side chain | 100 | 0.829 | |
| B S252 | F1 D1 | D Y53 | CDRH-2 | Hydrogen bond | Side chain - Backbone | 99.93 | -0.529 | G (1) |
| B S252 | F1 D1 | D H110, D Y53 | CDRH-3, CDRH-2 | Hydrogen bond | Side chain - Side chain | 99.93 | -0.529 | G (1) |
| B D255 | F1 D1 | E Y93 | CDRL-3 | Hydrogen bond | Backbone - Backbone | 99.93 | -0.529 | G (1) |
| B D255 | F1 D1 | E T95 | CDRL-3 | Hydrogen bond | Backbone - Side chain | 99.93 | -0.529 | G (1) |
| B E290 | F1 D1 | E E28, E S29 | CDRL-1, CDRL-1 | Hydrogen bond | Side chain - Backbone | 100 | 0.829 | |
| B E339 | F1 D1 | E S31, E Y93 | CDRL-1, CDRL-3 | Hydrogen bond | Side chain - Side chain | 100 | 0.829 | |

| | | | | | | | | |
|---|---|---|---|---|---|---|---|---|
| B E339 | F1 D1 | E Q69 | FRL-3 | Hydrogen bond | Backbone - Side chain | 100 | 0.829 | |
| B G254 | F1 D1 | E S92, E Y93 | CDRL-3, CDRL-3 | Hydrogen bond | Backbone - Backbone | 85.52 | -1.371 | D (207) N (1) |
| B K44 | F2 | E S29 | CDRL-1 | Hydrogen bond | Side chain - Backbone | 100 | 0.829 | |
| B S289 | F1 D1 | E D1 | FRL-1 | Hydrogen bond | Side chain - Side chain | 100 | 0.829 | |

### FECT and 4F09

| F chain & residue | Location | 4F09 residue | Location | Interaction Type | Atom Interaction | H residue frequency | H residue conservation index | Variants |
|---|---|---|---|---|---|---|---|---|
| A Q192 | F1 DIII | F Y106 | CDRH-3 | Hydrogen bond | Backbone - Backbone | 99.86 | -0.529 | R (2) |
| A Q200 | F1 DIII | F Y108 | CDRH-3 | Hydrogen bond | Side chain - Side chain | 100 | 0.829 | |
| A S194 | F1 DIII | F Y106 | CDRH-3 | Hydrogen bond | Backbone - Backbone | 100 | 0.829 | |
| B N158 | F1 HRN | F D109 | CDRH-3 | Hydrogen bond | Side chain - Backbone | 100 | 0.829 | |
| B N183 | F1 DIII | F Y106 | CDRH-3 | Hydrogen bond | Backbone - Side chain | 99.79 | -0.541 | D (3) |
| B N183 | F1 DIII | F Y113 | CDRH-3 | Hydrogen bond | Side chain - Side chain | 99.79 | -0.541 | D (3) |
| B N184 | F1 DIII | F Y108 | CDRH-3 | Hydrogen bond | Side chain - Side chain | 99.93 | -0.531 | T (1) |
| B N489 | F2 (N61) | F S111 | CDRH-3 | Hydrogen bond | Glycan - Backbone | 100 | 0.829 | |
| B N489 | F2 (N61) | F K67 | FRH-3 | Hydrogen bond | Glycan - Side chain | 100 | 0.829 | |
| A C195 | F1 DIII | G N93 | CDRL-3 | Hydrogen bond | Backbone - Side chain | 99.93 | -0.529 | S (1) |
| A C195 | F1 DIII | G S29 | CDRL-1 | Hydrogen bond | Backbone - Side chain | 99.93 | -0.529 | S (1) |

| | | | | | | | | |
|---|---|---|---|---|---|---|---|---|
| A N191 | F1 DIII | G K51 | CDRL-2 | Hydrogen bond | Side chain - Side chain | 99.72 | -1.361 | S (3) K(1) |
| A N66 | F2 | G R32, G S31, G S68 | CDRL-1, CDRL-1, | Hydrogen bond | Side chain - Side chain | 99.58 | -1.367 | D (4) K (2) |

**Table S2. Summary of H- and F-mAb contact residues as determined by cryo-EM.** Shown are the interacting residues between $H_{ECT}$ and protective mAbs 1C08 and 4D08, and less potent mAb 1C02, and $F_{ECT}$ and protective mAbs 3A12 and 4F09. Contact residues for each mAb are separated into heavy chain (HC) and light chain (LC) residues. The first letter of each residue refers to the chain, followed by amino acid and position. Interaction types, atom interaction state (glycoprotein - antibody), and conservation scores are also provided. Conservation scores were calculated and normalized using AL2CO. The maximum score for a fully conserved residue is 1.48 for H residues and 0.83 for F residues.
